# Lateral Hypothalamic Glutamate and GABA Neurons Cooperatively Shape Striatum-Wide Dopamine Dynamics During Consumption

**DOI:** 10.1101/2025.04.09.648052

**Authors:** Adam Gordon-Fennell, Barbara M. Benowitz, Joumana M. Barbakh, Isabella Montequin, Anthony Campuzano, Hannah P. Stevenson, Elena Lu, Madelyn M. Hjort, Ethan Ancell, Madalyn Critz, Daniela Witten, Garret D. Stuber

## Abstract

The lateral hypothalamic area (LHA) contains GABAergic and glutamatergic neurons that converge on the midbrain dopamine system and exert opposing influences on consummatory feeding behavior. However, the activity dynamics of these populations during consumption and their impact on striatal dopamine release remain poorly understood. Here, we show that LHA GABAergic and glutamatergic neurons independently scale their activity during the consumption of rewarding and aversive solutions while cooperatively regulating dopamine release along the anterior-posterior axis of the striatum in mice. Dopamine release exhibited widespread modulation during consumption, with anterior striatal regions showing stronger representation of solution value and its recent history—an effect dependent on LHA activity. These findings suggest that the LHA acts as a central regulator of striatal dopamine release during consumption, providing a mechanism through which hypothalamic circuits influence motivational and consummatory behaviors.

## INTRODUCTION

Consummatory behavior is necessary to obtain calories and nutrients that sustain life. The regulation of food and fluid consumption relies on a distributed neural network that integrates homeostatic need, sensory cues, and learned associations. Dopamine (DA) signaling regulates consumption behavior^1,2^ and is central to motivation^3–5^, yet while DA responses to predictive cues are well-characterized^6–8^, the spatiotemporal DA dynamics during consumption remain poorly characterized. Understanding how and where dopamine is released during consumption is necessary to isolate future interventional sites to reduce aberrantly motivated consumption. It is also vital to characterize candidate neural circuitry that modulates dopamine release during consumption and could serve as an additional therapeutic target. The lateral hypothalamic area (LHA) is a key integrator of metabolic state and feeding-related cues^9–11^, with projections to midbrain DA neurons^12–15^ that could modulate DA release throughout the striatum during consumption. Therefore, the purpose of this study is to test the hypothesis that LHA circuits influence striatal DA signaling during consumption-related behaviors.

The LHA contains genetically distinct GABAergic (LHA^GABA^) and glutamatergic (LHA^Glut^) neurons^16–19^ that exert opposing effects on feeding and reinforcement: stimulation of LHA^GABA^ neurons enhance consumption and drive positive valence^20–23^, while stimulation of LHA^Glut^ neurons suppress consumption and drive negative valence^18,21,22,24,25^. Despite these opposing roles, both populations are display increases in their calcium activity during consummatory behavior^18,20,23,26^, suggesting they work in tandem to regulate consummatory behavior. One potential site of convergence for these distinct populations is the midbrain DA system, where LHA^GABA^ and LHA^Glut^ neurons send dense projections to the ventral tegmental area (VTA) that shapes DA release in the nucleus accumbens caudal core (NAcCC)^12^. Additionally, the LHA projects directly to DA neurons in the VTA and substantia nigra pars compacta (SNc)^14,15^ and influences DA activity through polysynaptic pathways, including the lateral habenula^24,27,28^, periaqueductal gray^29,30^, raphe ^31,32^, and the peri-locus coeruleus^1,33^. Notably, the LHA also synapses on DA neurons that project to tail of the striatum (TS) ^34^, a region implicated in aversive processing and avoidance reinforcement ^7,35,36^. Given the complexity of the anatomical connectivity between the LHA and midbrain DA system, this study specifically sought to establish how LHA^GABA^ and LHA^Glut^ populations regulate DA release throughout the striatum during consummatory behavior.

Although DA release throughout the striatum encodes reward, motivation, and aversion, its dynamics during consumption remain critically underexplored—even though consumption is integral to most reward learning paradigms. Supporting the need for characterization of striatum-wide DA dynamics, DA levels in the ventral striatum correlate with sucrose concentration^37^ and reward volume^38,39^, but consumption-driven DA signals vary across ventral striatum subregions^22,40,41^. In addition to its implications for aberrant food drive, a deeper understanding of DA in this context will facilitate additional interpretation of previous studies that incorporate reward learning. This advance hinges upon disentangling DA representations of solution value (e.g., concentration) from those of consummatory behavior (e.g., licking), since both often co-occur in time. Adapting the Davis Rig traditionally used to quantify taste-driven consumption by delivering a range of discrete solutions^42^ for head-fixed mice^22^ greatly reduces signal confounds from approach behaviors^20,43–45^ and higher-order cognitive processes^46–48^, streamlining the dissociation of solution properties from consummatory actions. Specifically, we isolate motoric aspects of consumption through fluid licking, which offers high temporal resolution by capturing each lick, and characterize solution ‘value’ through the concentration of a single appetitive or aversive taste (e.g., sucrose or NaCl). This tightly-controlled task allowed us to characterize how LHA populations influence striatum-wide DA signaling during consumption.

Here, we combine dual-color fiber photometry, optogenetics, and multi-site DA recordings to investigate how LHA^GABA^ and LHA^Glut^ neurons regulate striatal DA release during consumption. Dual-color fiber photometry in LHA^GABA^ and LHA^Glut^ neurons revealed their simultaneous but distinct activity patterns during consumption of both rewarding and aversive solutions. Optogenetic manipulations of LHA^GABA^ and LHA^Glut^ neurons, combined with multi-site fiber photometry throughout the striatum, established the coupling between LHA activity and DA release across striatal subregions. Finally, using a generalized linear model, we identified spatiotemporal encoding of DA release during consumption, distinguishing the contributions of motoric aspects of consummatory behavior and solution value. Altogether, these findings demonstrate how bidirectional LHA encoding shapes spatially organized DA release to regulate motivation and consummatory behavior.

## RESULTS

### LHA^GABA^ and LHA^Glut^ exhibit unique activity scaling patterns during consummatory behavior

Before characterizing how LHA influences striatum-wide DA release during consummatory behavior, we first assessed the simultaneous dynamics of LHA^GABA^ and LHA^Glut^ populations during consumption. To this end, we utilized the Open-Source Head-fixed Rodent Behavioral Experimental Training System (OHRBETS) head-fixed brief-access taste task^22^ in which head-fixed mice consume one of five solutions presented in a pseudorandom order (2 trials per solution every 10 trials) in 3s windows (**Figure 1A-E**). To elicit a range of consumption, we modulated homeostatic demand by food-restriction (FR) or water restricting (WR) mice and providing a range of rewarding sucrose (Suc) concentrations or aversive NaCl concentrations. We used 3 primary restriction / solution configurations:

- WR:Suc: thirst-driven high homeostatic demand for all sucrose concentrations
- FR:Suc: hunger-driven high demand for high sucrose concentrations and low aversion for low sucrose concentrations
- WR:NaCl: thirst-driven high demand for low NaCl concentrations and high aversion for high NaCl concentrations

**Figure 1.**
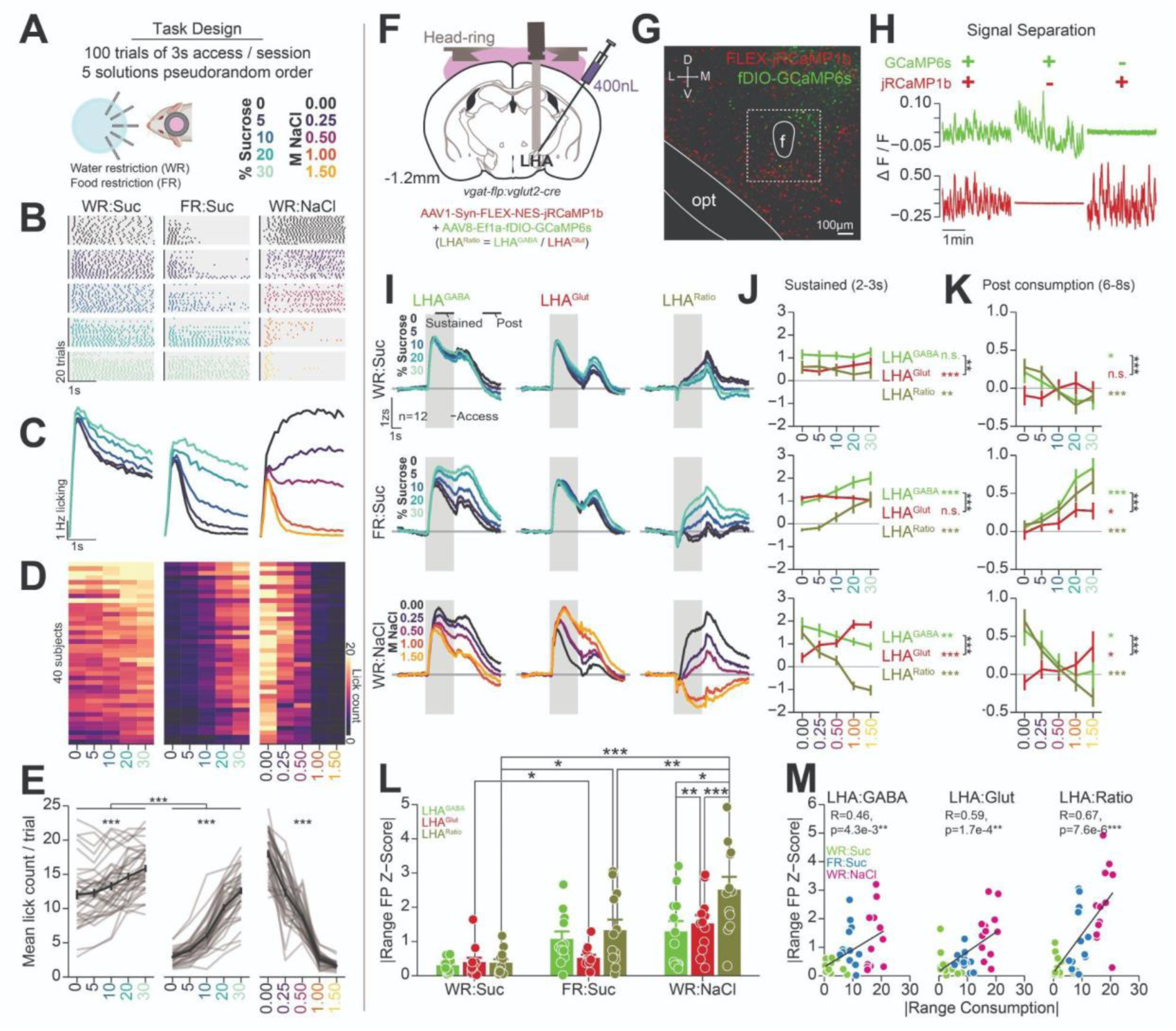
Tandem LHA^GABA^ and LHA^Glut^ dynamics during consummatory behavior. (A) Task design: mice underwent the multi-spout brief-access taste task where they were given 100 trials of 3 s access to one of five solutions presented in pseudorandom order (**see methods**). Mice were water restricted and provided access to five concentrations of sucrose (WR:Suc), food restricted and provided access to five concentrations of sucrose (FR:Suc), water restricted and provided to multiple concentrations of NaCl (WR:NaCl). Throughout the figure, color depicts the sucrose or NaCl concentration provided on the corresponding trial. (B) Licking raster for a representative subject in one session for each restriction/solution configuration. Trials are sorted by solution and trial number (earlier trials are lower), the vertical line indicates spout extension, and each dot indicates a lick. (C) Binned lick rate (100ms / bin) across all trials for all mice across each restriction/solution configuration (n trials=WR:Suc, 14,100; FR:Suc, 14,500; WR:NaCl 14,200). (D) Heat map depicting the mean trial lick count for each solution for subjects included throughout this manuscript (n=40), with subject identity consistent across rows. (E) Line plot depicting the mean trial lick count for each solution for subjects included throughout this manuscript, showing enhanced licking for higher concentrations of sucrose and reduced licking for higher concentrations of NaCl. Note that the range of licking is greater for sucrose under food restriction compared to water restriction. Asterisks over line plots indicate results from independent one-way repeated measures [RM] ANOVA performed on each restriction/solution configuration (within subject solution effect: WR:Suc: F(4, 156)=35.16, ***p=6.47e-21; FR:Suc: F(4, 156)=389.53, ***p=4.62e-80; WR:NaCl: F(4, 156)=387.30, ***p=6.95e-80, all Tukey Honest Significant [HSD] post hoc comparisons between solution concentrations are significant except WR:Suc 0 vs 5%). Asterisks between WR:Suc and FR:Suc indicate result from a two-way RM ANOVA with restriction state and solution within subject factors (F(4, 156)=55.15, ***p=6.51e-29, all HSD post hoc comparisons between restriction states with the same solution are significant) (F) Cartoon depicting approach for performing simultaneous dual-color imaging for LHA^GABA^ and LHA^Glut^ neurons. (G) Histology image showing expression of jRCaMP1b and fDIO-GCaMP6s in the LHA (annotations, f: fornix; opt: optic tract). (H) Validation of dual-color separation using lock-in demodulated fiber photometry. Top row displays the demodulated signal from the 465 nm channel while the bottom row displays the red 560 nm channel. Columns display signals from mice expressing combinations of GCaMP6s and jRCaMP1b as indicated by icons above. (I) LHA^GABA^ and LHA^Glut^ bulk-calcium photometry signals during the multi-solution brief-access taste task (n=12 mice). Bulk calcium activity for LHA^GABA^ (left column), LHA^Glut^ (center column), and the LHA^GABA^ / LHA^Glut^ ratio photometry signal (LHA^Ratio^, left column), during consumption for each restriction/solution configuration (rows). Grey rectangle indicates the 3 s access window, and annotations indicate the time epoch for the sustained and post windows summarized in (J) and (K). (J-K) Mean photometry signals in the LHA^GABA^, LHA^Glut^, and LHA^Ratio^ during sustained (J) and post consumption (K) time windows (n=12 mice, 3 LHA signal IDs per mouse [LHA^GABA^, LHA^Glut^, LHA^Ratio^]). Colored asterisks indicate main within subject solution effects results from independent one-way RM ANOVA performed on each restriction/solution configuration, and black asterisks indicate interaction effects from a two-way RM ANOVA with LHA subpopulation and solution within subject factors. (J) Sustained window: Top row shows WR:Suc (one-way RM ANOVA solution effect, LHA^GABA^: F(4, 44)=2.01, p=0.11; LHA^Glut^: F(4, 44)=5.96, ***p=6.36e-4; LHA^Ratio^: F(4, 44)=4.53, **p=3.74e-3; two-way RM ANOVA interaction F(4, 44)=4.69, **p=3.10e-3), middle row shows FR:Suc (one-way RM ANOVA solution effect, LHA^GABA^: F(4, 44)=19.69, ***p=2.38e-9; LHA^Glut^: F(4, 44)=0.81 p=0.52; LHA^Ratio^: F(4, 44)=16.12, ***p=3.36e-8; two-way RM ANOVA interaction, F(4, 44)=13.06, ***p=4.29e-7), and bottom row shows WR:NaCl (one-way RM ANOVA solution effect, LHA^GABA^: F(4, 44)=3.88, **p=8.75e-3; LHA^Glut^: F(4, 44)=17.37, ***p=1.28e-8; LHA^Ratio^: F(4, 44)=37.36, ***p=1.28e-13; two-way RM ANOVA interaction, F (4, 44)=29.11, ***p=7.40e-12) (K) Post consumption window: Top row shows WR:Suc (one-way RM ANOVA solution effect, LHA^GABA^: F(4, 44)=3.10, *p=0.026; LHA^Glut^: F(4, 44)=1.18, p=0.33; LHA^Ratio^: F(4, 44)=13.59, ***p=2.72e-7; two-way RM ANOVA interaction: F(4, 44)=12.36, ***p=8.04e-7), middle row shows FR:Suc (one-way RM ANOVA solution effect, LHA^GABA^: F(4, 44)=15.77, ***p=4.43e-8; LHA^Glut^: F(4, 44)=3.06, **p=0.026; LHA^Ratio^: F(4, 44)=6.87, ***p=2.19e-4; two-way RM ANOVA interaction : F(4, 44)=6.54, p=3.20e-4), and bottom row shows WR:NaCl (one-way RM ANOVA solution effect, LHA^GABA^: F(4, 44)=2.78, *p=0.038; LHA^Glut^: F(4, 44)=3.69, *p=0.011; LHA^Ratio^: F(4, 44)=10.21, ***p=6.21e-6; two-way RM ANOVA interaction : F(4, 44)=9.97, ***p=7.84e-6) (L) Comparison of the absolute range of the mean photometry signal (response during highest concentration minus response during lowest concentration) for each LHA signal ID (LHA^GABA^, LHA^Glut^, and LHA^Ratio^) and restriction/solution configuration (columns). Asterisks indicate significant HSD post hoc comparisons (two-way RM ANOVA, within factor LHA signal: F(2, 22)=15.67, ***p=5.87e-5; within factor configuration, F(2, 22)=13.69, ***p=1.37e-4; Interaction: F(4, 44)=30.20, ***p=4.14e-12) (M) Linear correlation between the absolute range of the consummatory response and the absolute range of the fiber photometry response for each subject across LHA signal ID (columns). Text depicts the results of a Pearson’s correlation test. Across all panels, statistics are displayed using the following convention: *p < 0.05, **p < 0.01, and ***p < 0.01. In fiber photometry traces (I), solid line depicts mean and shaded line depicts SEM calculated across all trials with licking.

Mirroring previous results^22^, mice consumed more on trials with higher concentrations of sucrose, with a greater range of consumption under FR:Suc than WR:Suc, likely due to weaker homeostatic demand for water in FR mice. Mice consumed less solution volume on trials with higher concentrations of NaCl (**Figure 1E**). These results validate that our approach effectively captures a range of consumption behaviors driven by both solution valence and homeostatic state.

Previous studies have shown that despite opposing effects on consumption and valence, LHA^GABA^ and LHA^Glut^ neurons increase activity during reward consumption^18,20^. However, it was unclear how these neuronal populations respond *simultaneously* when animals consume stimuli that span a range of reward values. We used dual-color fiber photometry to simultaneously record LHA^GABA^ and LHA^Glut^ activity while mice performed the OHRBETS brief-access taste task (**Figure 1F-M**). We used double transgenic mice expressing Cre recombinase in LHA^Glut^ neurons (*Slc17a6^tm2(cre)Lowl^/J* #016963 [*Vglut2-cre*]^49^) Flp recombinase in LHA^GABA^ neurons (*B6.Cg-Slc32a1^tm1.1(flpo)Hze^/J* #029591 [*Vgat-flp*]^49^), injected the LHA with a virus cocktail of a cre-dependent red calcium indicator (AAV1-Syn-FLEX-NES0-jRCaMP1b^50^) and a Flp-dependent green calcium indicator (AAV8-Ef1a-fDIO-GCaMP6s^51^), and implanted an optical fiber overlaying the LHA (**Figure 1F**). This approach produced robust, non-overlapping expression of both indicators in the LHA^16,18,19^ (**Figure 1G**) and enabled clear separation of green (LHA^Glut^) and red (LHA^GABA^) photometry signals (**Figure 1H**). We then analyzed each population independently as well as their tandem signaling by quantifying the ratio of calcium activity between LHA^GABA^ and LHA^Glut^ neuronal populations (LHA^Ratio^).

LHA^GABA^ and LHA^Glut^ calcium photometry signals exhibited distinct activity scaling during consumption across restriction and solution configuration (**Figure 1I-M**). Although both populations showed increased signals at consumption onset, calcium activity during the sustained portion of the access window varied depending on restriction and solution conditions. Under WR:Suc, both LHA^GABA^ and LHA^Glut^ signals increased during consumption as expected^18,26^, but displayed relatively weak scaling during the sustained and post-consumption windows (**Figure 1I-K, top row**). Despite the low-magnitude of scaling, the LHA^Glut^ signal during the sustained window positively correlated with solution value and consumption amount, while the LHA^GABA^ signal in the post-consumption window negatively correlated with the previous trial’s solution value. Under FR:Suc, which elicited a broader range of consumption responses, the LHA^GABA^ signal displayed strong scaling that positively correlated with solution value and consumption amount, while the LHA^Glut^ signal increased activity during consumption but did not scale (**Figure 1I-K, middle row**). In the post-consumption window, both signals scaled in response to the preceding trial’s solution value, with LHA^GABA^ neurons exhibiting a stronger effect. Under WR:NaCl, the LHA^GABA^ signal scaled positively and LHA^Glut^ activity scaled negatively with both solution value and consumption amount during the sustained and post-consumption windows (**Figure 1I-K, bottom row**).

The LHA^Ratio^ exhibited distinct scaling patterns across restriction and solution conditions. Under FR:Suc, the LHA^Ratio^ displayed positive scaling, increasing with solution value and consumption, and showing minimal reductions below baseline for less rewarding solutions. Under WR:NaCl, the LHA^Ratio^ displayed bidirectional scaling, increasing for the most valuable solutions (water; LHA^GABA^ > LHA^Glut^) and decreasing for the most aversive solutions (high-molar NaCl; LHA^Glut^ > LHA^GABA^). To assess how scaling of each signal related to consumption, we compared the absolute range of activity during the sustained window by subtracting responses to the least valuable solution from those to the most valuable solution (**Figure 1L**). The LHA^Ratio^ tracked the range of consumption across configurations better than LHA^GABA^ or LHA^Glut^ alone (**Figure 1L, M**).

In summary, LHA^GABA^ and LHA^Glut^ bulk calcium signals exhibited distinct patterns during consumption that were shaped by homeostatic demand and solution value. LHA^GABA^ neurons exhibited scaling of activity that was positively correlated with solution value, while LHA^Glut^ neurons exhibited scaling of activity negatively correlated with solution value during consumption of aversive solutions, producing enhanced and bidirectional LHA^Ratio^ dynamics. These findings suggest that the tandem activity of LHA^GABA^ and LHA^Glut^, as reflected in the LHA^Ratio^, cooperatively regulates downstream circuits to coordinate appropriate consummatory responses.

### LHA^GABA^ and LHA^Glut^ neurons collectively regulate DA dynamics across the striatum

Our next step in understanding how LHA influences striatum-wide DA release during consummatory behavior was to characterize the casual impacts of LHA cell type specific stimulation on striatum-wide DA dynamics. To investigate how LHA^GABA^ and LHA^Glut^ activity influences DA dynamics throughout the striatum, we used a multi-fiber approach combining optogenetic stimulation with simultaneous DA recordings (**Figure 2A**). We expressed a Cre-dependent, red-shifted excitatory opsin (AAV5-Syn-FLEX-rc[ChrimsonR-tdTomato] ^52^) bilaterally in the LHA of *Vgat-Cre* mice to stimulate LHA^GABA^ neurons and in *Vglut2-Cre* (*Slc17a6^tm2(cre)Lowl^/J* #016963^49^) mice to stimulate LHA^Glut^ neurons. As a control, we expressed a static red fluorophore (DIO-mCherry) in the LHA of a subset of *Vgat-Cre* and *Vglut2-Cre* mice. To measure DA levels, we expressed the fluorescent DA sensor GRAB-DA2m^53^ and implanted optical fibers in six striatal regions spanning the anterior-posterior axis: nucleus accumbens core rostral (NAcCR), NAcCC, nucleus accumbens ventral lateral shell (NAcShL), dorsal medial striatum (DMS), dorsolateral striatum (DLS), and tail of the striatum (TS) (**see Methods**).

**Figure 2.**
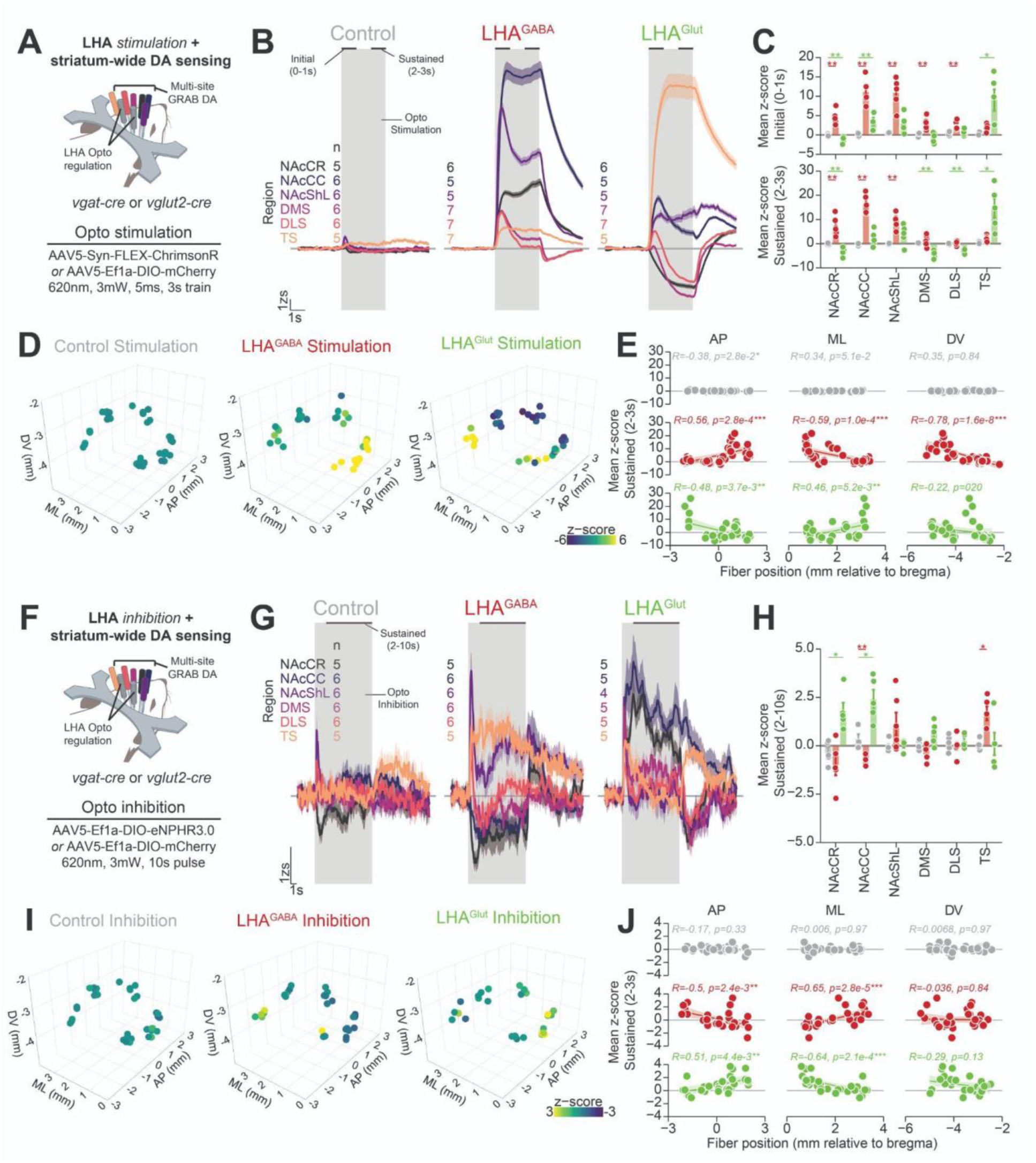
LHA^GABA^ and LHA^Glut^ collectively regulate DA release across the striatum (A-E) Striatal DA response to optogenetic stimulation of LHA^GABA^ or LHA^Glut^ neurons. (A) Cartoon depicting experimental approach for optogenetic stimulation of the LHA^GABA^, LHA^Glut^, or control (mCherry in *vgat-cre* or *vglut2-cre* mice) groups. (B) GRAB-DA response to laser emission in control (left), LHA^GABA^ Chrimson (middle), and LHA^Glut^ Chrimson (right) groups. The grey rectangle indicates the 3 s stimulation window, line color indicates striatal region, horizontal black lines indicate time epochs from which means were calculated for (C), and numerical values indicate the number of fibers per striatal region within each group. (C) Mean GRAB-DA response during the initial window (top) or sustained window (bottom). Asterisks indicate results from a Wilcoxon rank sum test (Bonferroni adjusted for multiple comparisons to the same control). (D) 3D map of striatal GRAB-DA response to laser light delivery during the sustained window (See Figure 2 **supplement 1** A-D for 2D representations). (E) Correlation between striatal position in the AP (left), ML (middle), and DV (right) dimensions versus the mean GRAB-DA response to laser emission during the sustained window. Text depicts the results of a Pearson’s correlation test. (F-J) Striatal DA response to optogenetic inhibition of LHA^GABA^ or LHA^Glut^ neurons. Figure annotations and statistics follow convention in corresponding panels in (A-E). (F) Cartoon depicting experimental approach for optogenetic inhibition of the LHA^GABA^, LHA^Glut^, or control (mCherry in *vgat-cre* or *vglut2-cre* mice) groups. (G) GRAB-DA response to laser light delivery. Transient increase in GRAB-DA signal observed across all groups while sustained changes observed only in LHA^GABA^ and LHA^Glut^ groups. (H) Mean GRAB-DA response during the sustained window. (I) 3D map of striatal GRAB-DA response to laser emission during the sustained window (See Figure 2 **supplement 1** E-H for 2D representations). (J) Correlation between striatal position and mean GRAB-DA response to laser emission. Across all panels, statistics are displayed using the following convention: *p < 0.05, **p < 0.01, and ***p < 0.01. In fiber photometry traces, solid line depicts mean and shaded line depicts SEM calculated across all trials (10-15 trials per fiber).

Optogenetic stimulation of LHA^GABA^ and LHA^Glut^ neurons induced widespread changes in DA dynamics across the striatum relative to controls (**Figure 2B-E**). Brief optogenetic stimulation at 20 Hz for 3 s produced time-dependent DA responses that varied by striatal subregion and consisted of two distinct components: a rapid DA rise within the first second of stimulation (“initial”) and a plateau during the final second (“Sustained”, **Figure 2B**). Stimulating LHA^GABA^ neurons increased DA levels in the NAcCR, NAcCC, NAcShL, DMS, and DLS during the initial phase but had no effect in the TS (**Figure 2C, top red**). During the sustained phase, DA remained elevated in the NAcCR, NAcCC, and NAcShL (**Figure 2C, bottom red**). In contrast, stimulating LHA^Glut^ neurons increased DA levels in the TS during the initial phase (**Figure 2C, top green**), while during the sustained phase, DA decreased in the NAcCR, DMS, and DLS and increased in the TS (**Figure 2C, bottom green**).

To assess how DA dynamics during LHA stimulation varied across the striatum, we analyzed the relationship between the position of the recording site and the effect of LHA^GABA^ or LHA^Glut^ stimulation (**Figure 2D-E, Figure 2 supplement 1A-D**). DA levels in response to LHA^GABA^ stimulation correlated positively with the anterior-posterior axis and negatively with the medial-lateral and dorsal-ventral axes (**Figure 2E, middle row**). The DA response to LHA^Glut^ stimulation showed the opposite pattern, correlating negatively with the anterior-posterior axis and positively with the medial-lateral axis (**Figure 2E, bottom row**). Control animals also exhibited significant correlations between laser stimulation and striatal position, but their DA responses were far weaker than those observed in LHA^GABA^ and LHA^Glut^ stimulation groups (**Figure 2E, bottom row, Figure 2B-C**).

We next examined how LHA^GABA^ and LHA^Glut^ neurons constitutively regulate striatal DA levels by optogenetically inhibiting these populations while recording DA levels. To achieve inhibition, we bilaterally expressed a Cre-dependent inhibitory opsin (DIO-eNPHR3.0^54^) in the LHA of *Vgat-Cre* mice to target LHA^GABA^ neurons and in *Vglut2-Cre* mice to target LHA^Glut^ neurons. As a control, we expressed a static red fluorophore (DIO-mCherry) in a subset of *Vgat-Cre* and *Vglut2-Cre* mice. Optogenetic inhibition of LHA^GABA^ and LHA^Glut^ neurons differentially shaped DA levels in a subset of striatal regions relative to controls (**Figure 2F-J**). Inhibiting LHA^GABA^ neurons caused a sustained DA decrease in the NAcCC and an increase in the TS (**Figure 2H**). Inhibiting LHA^Glut^ neurons led to a sustained DA increase in the NAcCR and NAcCC (**Figure 2H**).

To assess how DA responses to LHA inhibition varied across the striatum, we analyzed the relationship between recording site position and the effects of LHA^GABA^ or LHA^Glut^ inhibition (**Figure 2I-J, Figure 2 supplement 1E-H**). The DA response to LHA^GABA^ inhibition correlated negatively with the anterior-posterior axis and positively with the medial-lateral axis (**Figure 2J, middle row**). The DA response to LHA^Glut^ inhibition showed the opposite pattern, correlating positively with the anterior-posterior axis and negatively with the medial-lateral axis (**Figure 2J, bottom row**). In control animals, laser illumination had no correlation with striatal axes (**Figure 2J, top row**).

These findings demonstrate that LHA^GABA^ and LHA^Glut^ neurons modulate striatal DA dynamics in distinct patterns along the anterior-posterior and medial-lateral axes: LHA^GABA^ coupling with DA level is *positively* correlated with the anterior-posterior axis (positively coupled in anterior subregions and negatively coupled in posterior regions) while LHA^Glut^ is *negatively* correlated with the anterior-posterior axis (negatively coupled in anterior subregions and positively coupled in posterior subregions). The interaction of these coupling gradients suggests a broader role for the LHA^Ratio^ in coordinating DA levels across the striatum.

### Evoked DA release in one striatal region produces minimal DA release in other striatal regions

After establishing that LHA^GABA^ and LHA^Glut^ neurons regulate DA dynamics across the striatum, we tested whether this wide-spread regulation occurs through interactions between DA release at subsets of striatal subregions that influence DA release at other subregions, as previously hypothesized^55–58^. Towards this end, we recorded DA levels at four striatal subregions within the same hemisphere while selectively evoking DA release at each subregion through the same optical fiber (**Figure 3**). To evoke DA release, we expressed a red-shifted excitatory opsin (AAV5-Syn-FLEX-rc[ChrimsonR-tdTomato]) in the midbrain of *DAT-Cre* (*Slc6a3^tm1(cre)Xz/^J #020080*) mice, resulting in targeted opsin expression within midbrain DA regions, including the VTA and SNc (**Figure 3B**). We then recorded DA levels using multi-site fiber photometry in mice expressing the fluorescent DA sensor GRAB-DA2m in four striatal subregions along the anterior-posterior axis: NAcCR, DMS, DLS, and TS (representative placements from a single subject shown in **Figure 3C**). This approach allowed us to simultaneously record DA levels at all four subregions while optogenetically evoking DA release at each subregion individually, permitting investigation of whether DA release at one subregion influenced DA levels in other subregions.

**Figure 3.**
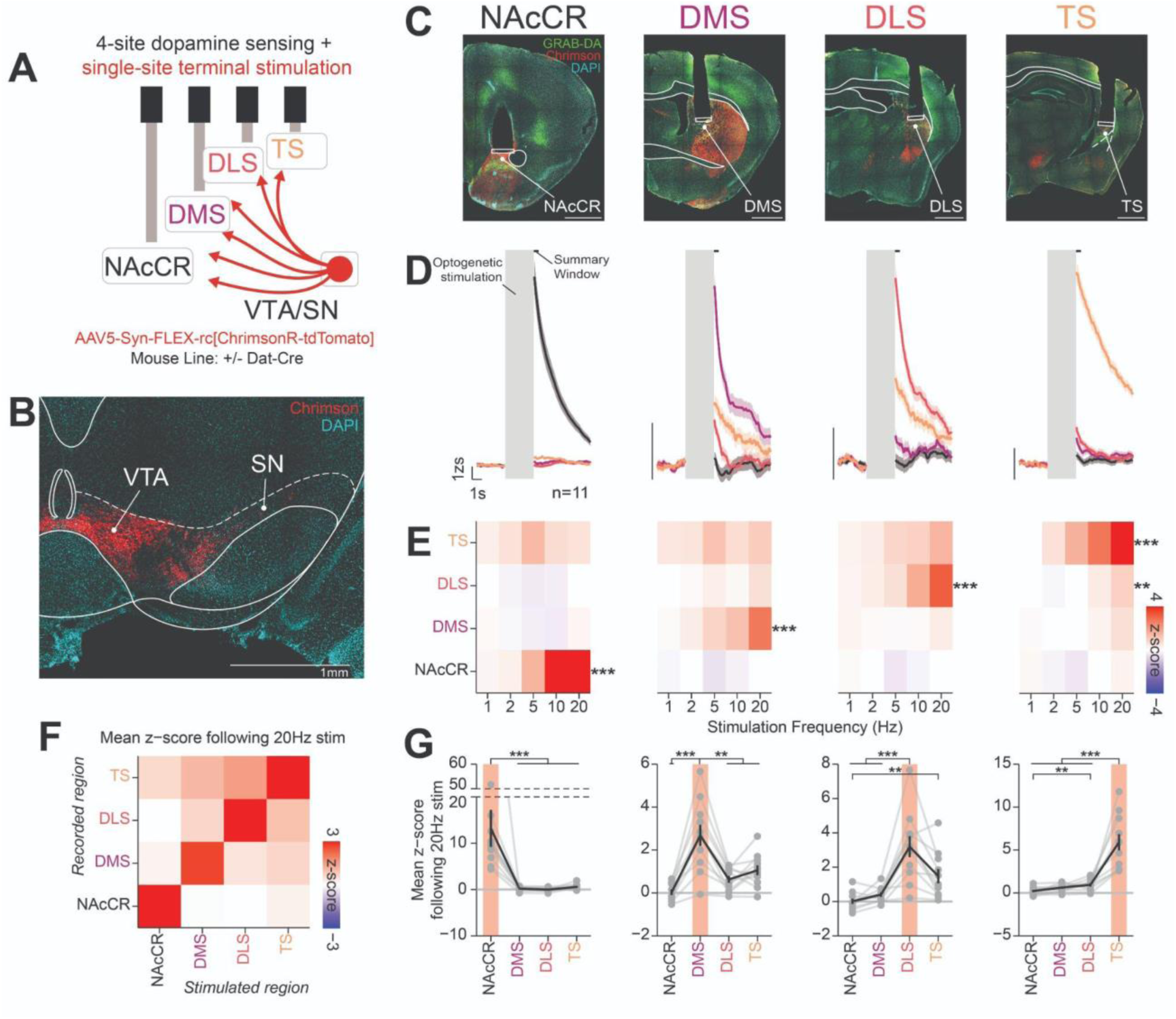
Evoked DA release in one striatal region produces minimal DA release in other striatal regions. (A) Cartoon depicting experimental approach for optogenetic stimulation of DA release at one striatal region while recording DA release across multiple subregions. (B) Expression of Chrimson-tdTomato in the VTA and SN in *Dat-Cre* mice. (C) Histology in a representative mouse (scale bar = 1mm). (D) GRAB-DA response across 4 striatal regions during 3s 20 Hz stimulation of DA release in a single subregion (n=11 mice, 10 stimulations per subregion). Each column represents the striatal region that was stimulated while line color indicates the striatal region that was recorded. Grey rectangle indicates optogenetic stimulation period, and horizontal black line indicates summary window used for means in (E-G). (E) Heatmap depicting the GRAB-DA response to optogenetic stimulation at each striatal region. Asterisks denote significant results from a Friedman rank sum test for stimulation frequency effect, adjusted using the Bonferroni correction for four striatal regions recorded (NAcCR stim:NAcCR response, Q=38.98, ***p=2.81e-7; DMS stim:DMS response, Q=25.16, ***p=1.87e-4; DLS stim: DLS response, Q=31.42, ***p=1.01e-5; TS stim: TS response, Q=41.82, ***p=7.28e-8; TS stim: DLS response, Q=21.24, **p=1.14e-3). (F) Heatmap showing the GRAB-DA response to 20 Hz stimulation, with the stimulated region represented on the x-axis and the recorded region on the y-axis. (G) Line plot showing the GRAB-DA response to 20 Hz stimulation, with striatal region that was recorded on the x-axis and red rectangle indicating the striatal region that was stimulated. Evoked DA in each subregion produced differential DA levels across striatal subregions, as determined using a Friedman rank sum test (NAcCR: Q=23.73, ***p=2.85e-5, DMS: Q=26.45, ***p=7.66e-6, DLS: Q=21.22, ***p=9.48e-5, TS: Q=26.02, ***p=9.45). Asterisks denote significant post hoc comparisons using a pairwise Wilcoxon rank sum exact test, adjusted using the Bonferroni correction. Across all panels, statistics are displayed using the following convention: *p < 0.05, **p < 0.01, and ***p < 0.01. In fiber photometry traces, solid line depicts mean and shaded line depicts SEM calculated across all trials (11 subjects x 10-15 stimulations per stimulated region).

Optogenetically evoked DA release produced frequency-dependent changes in DA levels only within the stimulated subregion for the NAcCR, DMS, and DLS (**Figure 3E**). In contrast, evoking DA release in the TS resulted in a small but significant frequency-dependent increase in DA within the DLS. We next examined the effect of 20 Hz stimulation alone to compare how stimulating one subregion influenced DA levels in other subregions (**Figure 3F-G**). A matrix visualizing stimulation and recording subregions revealed a modest relationship between DA levels, primarily dependent on the proximity between the evoked and recorded subregions (**Figure 3F**). Stimulation at each subregion resulted in significantly greater DA levels within the stimulated subregion compared to all others. However, in the DLS, DA levels following stimulation did not significantly differ between the DLS and TS (**Figure 3G**). Some interactions between subregions emerged following DA terminal stimulation in the DLS and TS, as indicated by greater DA levels in the TS compared to the NAcCR when stimulating the DLS, and greater DA levels in the DLS compared to the NAcCR when stimulating the TS.

These results demonstrate that evoked DA release at one striatal subregion does not substantially shape DA levels at other subregions, apart from a modest relationship between the DLS and TS. This localized and modest interaction suggests that DA release within the striatum is largely independent across subregions. In the context of the overarching question regarding how the LHA regulates DA release across the striatum, these results suggest that the LHA influences DA release at distinct subregions directly rather than indirectly modulating DA in a subset of striatal subregions that propagates throughout the striatum.

### Spontaneous DA transients and their kinetics are correlated with position within the striatum

Before understanding how LHA influences DA release dynamics during consumption, we sought to characterize striatum-wide dopamine dynamics unique to consumption, which itself required comparisons to dopamine release at rest, defined here as a baseline period 2-minutes prior to every behavioral session (**Figure 4**). We recorded DA from six striatal subregions (NAcCR, NAcCC, NAcShL, DMS, DLS, and TS) using multi-site fiber photometry paired with the DA sensor GRAB-DA2m while mice were head-fixed (**Figure 4C**, representative placements from a single subject shown in **Figure 4A**, all analyzed placements shown in **Figure 4B**, 3d model of fiber implant shown in **Figure 4 supplement 1**; 3D striatal maps of transient rate and kinetics shown in **Figure 4 supplement 2**; correlations between position and kinetics shown in **Figure 4 supplement 3**).

**Figure 4.**
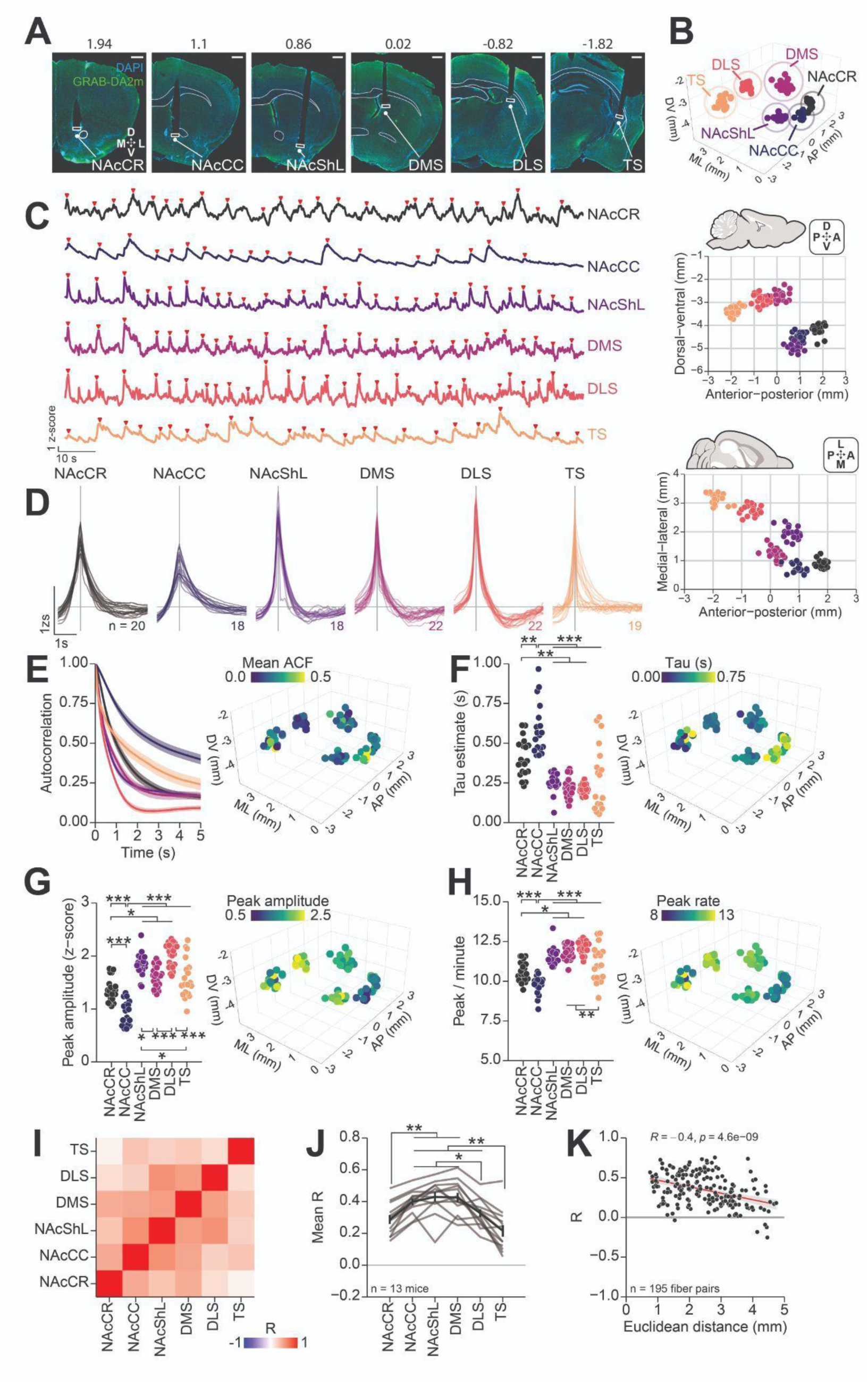
Spontaneous DA dynamics across the striatum. (A) Histology in a representative mouse. NAcCC and NAcShL fibers were located in the opposite hemisphere and images have been inverted horizontally (see Figure 4 **supplement 1** for implant design) (B) Position of 119 fiber tips in the striatum used in analysis represented in 3D (top), striatal (middle), and horizontal (bottom) orientations with color indicating the striatal region. (C) GRAB-DA trace at rest during head-fixation in a representative mouse with red triangles indicating detected peaks. (D) Mean GRAB-DA trace of spontaneous transients centered on time of peak. (E-H) Spontaneous GRAB-DA dynamics across striatal regions. Autocorrelation over a 5s window (E), tau estimates based on decay following spontaneous transients (F), peak amplitude of spontaneous transients (G), and the rate of spontaneous transients (H). 3D maps depicting the location of fiber tips color coded with the corresponding variable (see Figure 4 **supplement 2** for 2D representations and Figure 4 **supplement 3** for correlation between position and each variable). (I-J) Correlation of GRAB-DA dynamics across striatal regions in mice with six accurately placed fibers. Heatmap illustrates the mean correlation between each pair of striatal regions (I), line plot shows the mean correlation of each region with all other regions (J, grey lines depict individual subjects, black line depicts mean across subjects), and scatter plot depicts the relationship between the Euclidean distance between fiber pairs and their corresponding correlation (K, points depict pairs of fibers within the same mouse). Across all panels, asterisks denote results from HSD post hoc tests using the following convention: *p < 0.05, **p < 0.01, and ***p < 0.01.

To analyze spontaneous DA kinetics, we applied a peak-detection algorithm (see methods) to identify DA transients (mean peri-peak time histograms in **Figure 4D**). Assessing DA signal autocorrelations at each subregion revealed that ventral subregions exhibited greater mean autocorrelation than dorsal subregions, with particularly high autocorrelation in the NAcCC (**Figure 4E, Figure 4 supplement 2B, Figure 4 supplement 3A**), as previously reported^59^. The decay rate, tau, varied significantly across subregions, with longer tau estimates in anterior, medial, and ventral portions of the striatum, as previously described^59,60^ (**Figure 4F, Figure 4 supplement 2C, Figure 4 supplement 3B**). The spontaneous DA transient rate and peak amplitude showed an inverse relationship with tau, exhibiting lower values in anterior, medial, and ventral subregions (**Figure 4G-H, Figure 4 supplement 2D-E, Figure 4 supplement 3C-D**). Among all subregions, the NAcCC displayed the highest autocorrelation and tau estimates but the lowest transient rate and peak amplitude, possibly reflecting unique regulation of DA release and reuptake in this subregion^61,62^. Altogether, these data reveal a structured relationship between spontaneous DA transients, their kinetics, and position within the striatum.

We next assessed the baseline correlation of DA levels across subregions in mice with accurate placements for all six fibers (**Figure 4I-K**). Visualizing the correlation matrix of DA dynamics across subregions (**Figure 4I**) revealed broad positive correlations, with stronger correlations observed between neighboring subregions compared to more distant ones, as reflected in the higher correlation values along the diagonal (**Figure 4I**). The mean R, computed as the mean correlation between each subregion and all others, exhibited an “inverted-U” pattern, with lower correlations in the most anterior (NAcCR) and posterior (DLS and TS) subregions and higher correlations in the mid-striatal axis (NAcShL and DMS). To determine whether this pattern resulted from the proximity of recorded subregions, we analyzed the relationship between Euclidean distance and DA signal correlation for all recorded subregion pairs (**Figure 4K**). We found a negative relationship, indicating that subregions closer in space showed higher DA correlation compared to those farther apart. These findings suggest that the baseline correlation of DA levels across the striatum is strongly influenced by spatial proximity, with neighboring subregions exhibiting greater correlation than distant ones.

### DA dynamics during consumption show distinct patterns that are correlated with position within the striatum

Having characterized striatal DA dynamics at rest, we next examined DA signaling during the consumption of rewarding and aversive solutions. Like our investigations of the LHA, we examined DA activity during consumption in the OHRBETS brief-access taste task under the three restriction/solution configurations (**Figure 1A-E, Figure 5**). To increase the number of striatal subregions analyzed, we incorporated a subset of data from a previous study^22^, which included simultaneous recordings from the NAcShM and NAcShL in mice using the same DA sensor and behavioral approach. We analyzed DA responses within three time-windows relative to trial onset (**Figure 5C**): licking onset (0.0-0.3 s; preceding the onset of licking in most trials with licking), sustained (2-3 s), and post (6-8 s following the offset of the access period).

**Figure 5.**
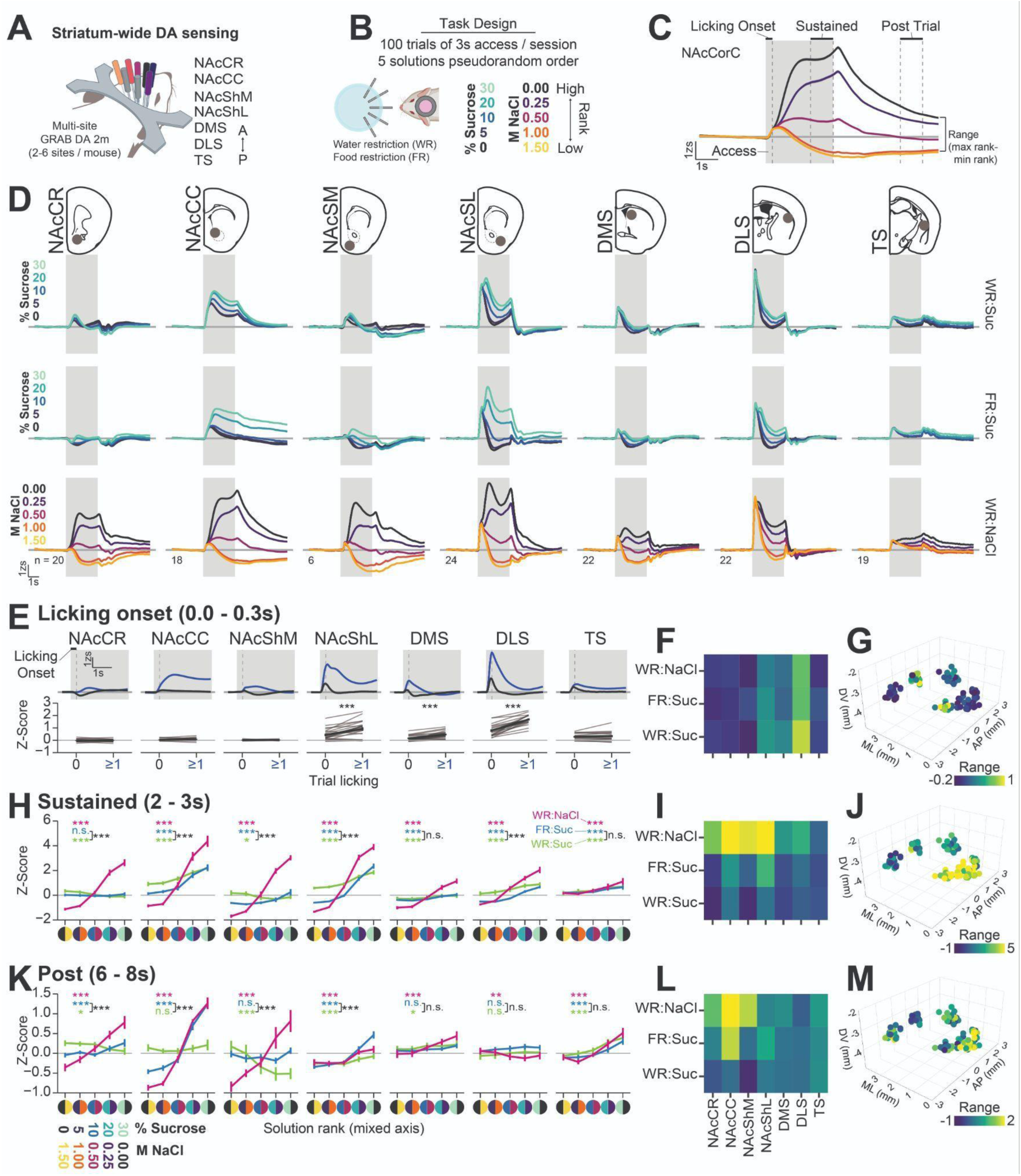
DA dynamics across the striatum during consumption. (A) Cartoon depicting the approach for multi-site striatum GRAB-DA recordings in head-fixed mice. (B) Brief-access taste task design (see Figure 1). (C) Diagram showing time epochs used in analysis (D) Perievent time histograms of GRAB-DA activity across 7 striatal regions (columns) and three solution/restriction configurations (rows). Grey rectangle indicates the 3s access period and color indicates the solution for the corresponding trial. (E-M) Analysis of GRAB-DA dynamics during consumption behavior across each striatal region divided based on time epoch: licking onset (E-G), sustained (H-J), and post consumption (K-M). Each epoch displays the mean GRAB-DA responses (E, H, and K), a heatmap depicting the mean GRAB-DA response across all solution/restriction configurations (F, I, and L), and a 3D map depicting the location of fiber tips color coded with the range of the corresponding variable (G, J, and M, see Figure 5 **supplement 2** for 2D representations and correlation between position and each variable). In fiber photometry traces, solid line depicts mean and shaded line depicts SEM calculated across all trials. In (H) and (K) the axis depicts the solution rank indicated by the color code (0→30% sucrose or 1.50→0.00M NaCl). In (E), (H), (K) centers depict mean and error bars depict SEM calculated across subjects and grey lines in (E) depict means for individual subjects. in (H) and (K) colored asterisks indicate one-way RM ANOVA solution effect, in (E) asterisks denote results of a paired T-Test, grey lines depict individual subjects and black line depicts mean across subjects), and black asterisks indicate two-way RM ANOVA solution and restriction interaction (see **Table S1** for the results from each statistical test).

Consumption of rewarding and aversive solutions coincided with dopamine scaling across a broad expanse of the striatum, with time-dependent changes varying by subregion and restriction/solution configuration (**Figure 5D-M**, representative subject shown in **Figure 5 supplement 1**). During the initial window, before licking onset in most trials, DA significantly increased in the NAcShL, DMS, and DLS, with greater increases on trials that included licking (mean across all trials shown in **Figure 5E**, divided by restriction/solution configuration in **Figure 5 supplement 3A**). Minimal differences in DA responses between trials with and without licking were observed across restriction/solution configurations (**Figure 5F**). 3D mapping of recording positions revealed relatively weak linear correlations between striatal position and DA responses during licking onset. Instead, a band of subregions along the anterior-posterior axis exhibited distinct DA dynamics prior to licking onset (**Figure 5, Figure 5 supplement 2A, D**).

Differences in the GRAB-DA rise-time and maximum derivative during consumption followed an anterior-posterior relationship (**Figure 5 supplement 5**). Posterior subregions displayed significantly faster rise times and maximum derivatives compared to anterior subregions^59^, extending previous reports of wave-like dynamics in DA release in the dorsal striatum^63,64^. The rapid rise time in these posterior subregions may facilitate the encoding of sensory components before behavior onset and could reflect a role in reinforcing behavioral initiation.

During the sustained window, in the final second of the access period, DA responses scaled with solution rank across all restriction/solution configurations (**Figure 5H-J**, individual subjects shown in **Figure 5 supplement 3B**). Higher-ranked solutions (water in WR:NaCl or 30% sucrose in FR:Suc and WR:Suc) elicited greater DA signals in nearly all striatal subregions, with two exceptions: the NAcCR, which showed a negative relationship with solution rank in WR:Suc and no relationship in FR:Suc, and the NAcShM, which displayed a negative relationship in WR:Suc. An interaction test confirmed that DA responses to different sucrose concentrations varied between WR and FR conditions in every recorded subregion except the DMS and TS. Under FR, DA responses exhibited stronger scaling, with more subregions showing a negative response to low sucrose concentrations. Additionally, consumption under WR:NaCl produced substantially stronger DA scaling across most subregions (**Figure 5I**). 3D mapping of recording positions revealed strong linear correlations between striatal position and DA scaling during the sustained window in WR:NaCl, with more anterior, medial, and ventral subregions exhibiting greater scaling (**Figure 5J, Figure 5 supplement 2B, E**).

During the post-consumption window, DA responses exhibited similar scaling patterns to the sustained window but with reduced magnitude in the posterior and dorsal striatum (**Figure 5K-M**, individual subjects shown in **Figure 5 supplement 3B**). Under the WR:NaCl configuration, DA responses maintained a positive relationship with solution rank in all subregions except the DLS, which exhibited a negative relationship. Under FR:Suc, DA responses maintained a positive relationship with solution rank in the NAcCR, NAcCC, NAcShL, and TS. Under WR:Suc, DA responses showed a positive relationship in the NAcShL and TS but a negative relationship in the NAcCR and NAcShM (**Figure 5K**). We observed restriction-dependent differences in DA responses to sucrose consumption in anterior and ventral striatal subregions (NAcCR, NAcCC, NAcShM, and NAcShL), but not in posterior and dorsal subregions (DMS, DLS, and TS). The WR:NaCl configuration produced substantially stronger DA scaling in anterior subregions, including the NAcCR, NAcCC, and NAcShM (**Figure 5L**). 3D mapping of recording positions revealed strong linear correlations between striatal position and DA scaling during the post-consumption window under WR:NaCl, with more anterior, medial, and ventral subregions exhibiting greater scaling (**Figure 5M, Figure 5 supplement 2C, F**).

In summary, DA responses in striatal subregions scaled robustly with solution value over time, revealing pronounced differences in dynamics along the anterior–posterior axis: posterior subregions exhibited a rapid enhancement before consumption, scaling then spread throughout the striatum and peaked in anterior subregions, and a prolonged separation in DA levels persisted in a subset of these anterior areas. However, these conclusions are limited due to temporal co-occurrence of key task parameters including the motoric (licking) and value (solution concentration) components of the consumption task.

### Spatial gradients of DA encoding across the striatum reflect both consummatory behavior and solution concentration

To disentangle the relative contributions of solution value and consummatory behavior to DA signaling, we conducted two initial analyses on the data described above (**Figure 5 supplement 4**). First, we computed linear correlations between DA signals and consummatory behavior across all trials (**Figure 5 supplement 4A-B**). These analyses revealed strong positive correlations between DA levels and lick count in most striatal subregions across restriction/solution configurations (**Figure 5 supplement 4A**), with stronger correlations observed under the WR:NaCl configuration compared to FR:Suc and WR:Suc in most subregions (**Figure 5 supplement 4B**). However, because trials were interleaved across multiple solutions, it was unclear whether these correlations reflected consumption behavior *per se* or were driven by solution concentration parameters. To address this, we binned trials by both consumption and solution concentration, creating a two-dimensional matrix of mean DA responses (**Figure 5 supplement 4C**). This heat map showed that, at a fixed solution concentration, greater consumption correlated with elevated DA signals in most subregions. Conversely, at a fixed level of consumption, more preferred solutions elicited higher DA activity, supporting independent contributions of consummatory behavior and solution value to DA signaling.

To formally separate these influences, we implemented a generalized linear model (GLM) to predict DA signals across subregions in the striatum^65–70^ using the following predictor sets: smoothed kernels for licking behavior, solution concentration (as a continuous variable ranging from 0 to 1), solution history (mean solution concentration in the previous three trials), and a “time in trial” capturing other sensory or timing-related effects independent of solution and behavior (**Figure 6A-F**, see **Methods**). We generated independent models for each combination of fiber and restriction/solution configuration. For the sake of simplicity, WR:NaCl is shown in **Figure 6A-F** and results from all models are shown in **Figure 6 supplement 1**. We assessed the contribution of each of the predictor sets to DA variability across striatal subregions by refitting the GLMs without each predictor set and computing the change in proportion of explained variance (ΔR^2^). We found broad encoding of licking across most subregions, with the strongest effects in the NAcCC and NAcShL (**Figure 6D**). In contrast, solution concentration (**Figure 6E**) and its history (**Figure 6F**) were most strongly represented in anterior and ventral regions, consistent with their established role in value-related processing. Lastly, the dorsal striatum, particularly the DLS, showed a pronounced contribution from the trial-aligned kernel, suggesting that non-solution sensory information or other task-related factors are encoded more prominently in these regions (**Figure 6C**).

**Figure 6.**
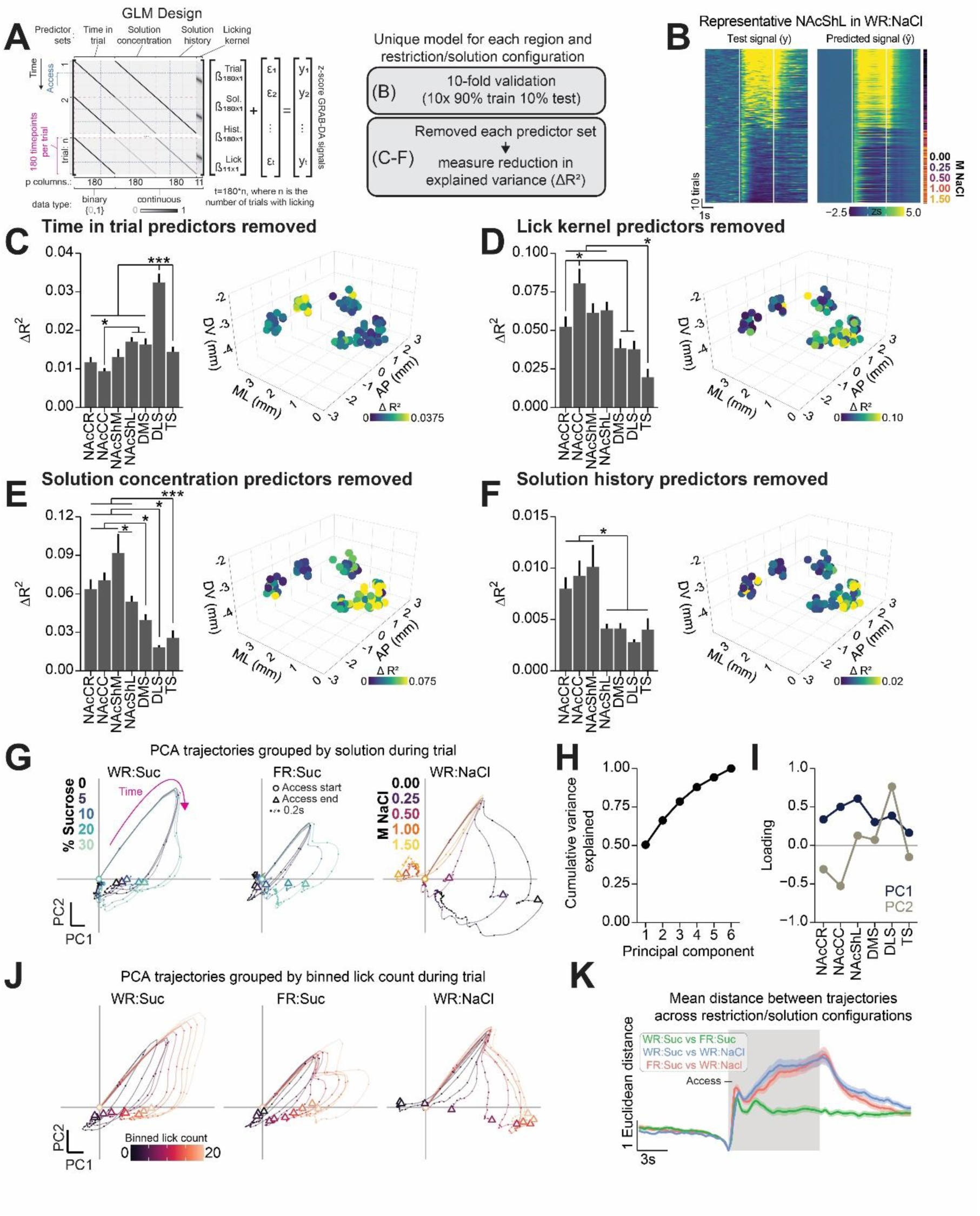
Unique representation of subcomponents of consumption in DA release across the striatum. (A-F) Generalized linear model analysis of striatal DA dynamics during consumption under the WR:NaCl restriction/solution configuration shown in Figure 5 (See Figure 6 **supplement 1** for results from all configurations). (A) Cartoon of approach to analysis (see **Methods**): For each fiber, and in each restriction/solution configuration, a separate linear model was fit to predict the GRAB-DA signal using predictor sets corresponding to time in trial, scaled solution concentration, scaled solution concentration history (mean of concentration over the last 3 trials), and licking (smooth kernel). (B) Test signals and predicted signals from 10-fold cross-validation for a representative fiber in the NAcShL during WR:NaCl. (C-F) Change in proportion of GRAB-DA variance explained (ΔR^2^) due to the removal of each predictor set: Time in trial predictor set (C, ANOVA, F(6,120)=21.48, ***p=5.11e-17), licking predictor set (D, ANOVA, F(6,120)=32.00, ***p=9.35e-23), solution concentration predictor set (E, ANOVA, F(6,120)=11.18, ***p=6.89e-10), and solution concentration history predictor set (F, ANOVA, F(6,120)=11.10, ***p=7.96e-10). Bar plots depict the ΔR^2^ across striatal subregions and 3D maps depict the location of fiber tips color coded with ΔR^2^ (see Figure 6 **supplement 2**, **3** for 2D representations and correlation between position and each variable). (G-K) PCA analysis of striatum-wide DA dynamics during consumption. (G) Mean 2D PCA trajectories from the onset of the access period to 8s post access, averaged across trials with a particular solution concentration (color) for a given solution/restriction configuration (horizontal facet). Circles indicate the beginning of access period, triangles indicate end of access period, each point within the trajectory is separated by 0.2s; see Figure 6 **supplement 4** for representative subject). (H) Scree plot corresponding to PCA analysis. (I) PC loadings across striatal regions for the first two PCs. (J) Mean 2D PCA trajectories during the access period, averaged across trials with a particular binned lick count for a given solution/restriction configuration. Additional details as in (G). (K) Euclidean distance in PC space between trajectories of the same trial lick count across solution/restriction configurations (solid line depicts mean and shaded line depicts SEM calculated across subjects). (K) The Euclidean distance between PC trajectories with the same trial lick count for each pair of solution/restriction configurations (solid line depicts the mean and shaded line depicts the SEM, across 10 subjects).

### Dimensionality reduction reveals distinct population-level DA dynamics when consuming rewarding vs. aversive solutions

We examined how striatal DA levels across multiple subregions vary with solution value and consummatory behavior by applying principal component analysis (PCA, see **Methods**) to concatenated data from all animals and restriction/solution configurations (**Figure 6G-L**). Plotting the mean trajectory of the first two principal components (PC1 and PC2) over time for each solution revealed a consistent clockwise rotation during consumption, with an initial rapid shift in PC space followed by a divergence that tracked solution value (**Figure 6G**). This divergence was particularly pronounced under the WR:NaCl configuration, suggesting that population-level DA signals collectively encode solution value during consumption.

To determine whether differences in consummatory behavior alone accounted for variations in PCA trajectories, we regrouped trials by lick count (**Figure 6J, K**). Under both WR:Suc and FR:Suc configurations, trials with the same lick count followed similar trajectories in PC space during the access period (**Figure 6K**). In contrast, under the WR:NaCl configuration, trajectories remained distinct even when matched for lick count, suggesting that aversive solutions introduce unique features into the striatum-wide DA response beyond differences in consumption behavior.

### Transient LHA inhibition reveals bidirectional modulation of consumption and striatal dopamine

With all pieces in place, we could finally determine how LHA populations influence striatum-wide DA signaling during consumption. We therefore asked whether transiently inhibiting LHA^GABA^ or LHA^Glut^ neurons would alter both behavior and striatum-wide DA signals during consumption of an aversive (1M NaCl) or rewarding (water) solutions under water restriction. To test this, we applied continuous optogenetic inhibition of LHA^GABA^ or LHA^Glut^ neurons (mCherry as control) beginning six seconds before each trial and extending through the six-second post-access window, while recording DA across multiple striatal subregions. Behaviorally, LHA^GABA^ inhibition suppressed water consumption, whereas LHA^Glut^ inhibition increased NaCl consumption (**Figure 7B–D**). These effects coincided with changes in DA dynamics: LHA^GABA^ inhibition reduced DA in the NAcCR, NAcCC, and DMS (depending on the solution), while LHA^Glut^ inhibition increased DA in the NAcCR and NAcShL (**Figure 7F**). Further analysis of the spatial distribution of these effects revealed correlations with the anterior-posterior and medial-lateral axes of the striatum: DA changes from LHA^GABA^ inhibition negatively correlated with anterior-posterior position and positively correlated with medial-lateral position, whereas LHA^Glut^ inhibition showed the opposite pattern (**Figure 7G-H**). Even after accounting for baseline DA shifts introduced by LHA inhibition before consumption (**Figure 7 supplement 1, Figure 7 supplement 2A**), these manipulations continued to produce bidirectional modulation of DA signals across multiple striatal subregions (**Figure 7 supplement 2B**).

**Figure 7.**
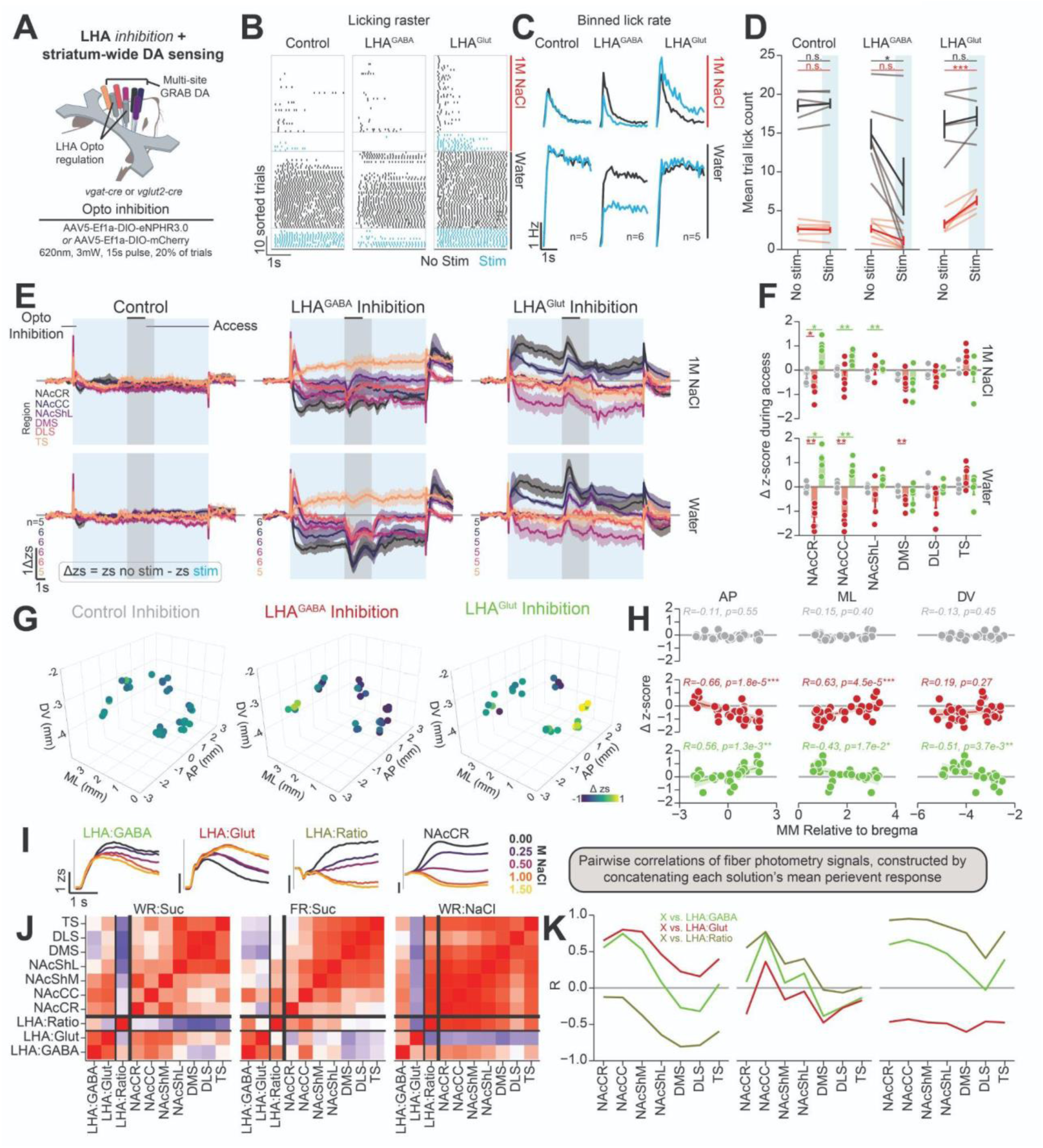
The LHA^GABA^ and LHA^Glut^ regulation of striatal DA during consumption. (A) Cartoon depicting the approach for optogenetic regulation of LHA neurons during multi-site striatum GRAB-DA recordings in head-fixed mice. (B) Licking raster for a representative subject from each group in one session. Trials are sorted by solution, laser stimulation, and trial number (earlier trials are lower), and each dot indicates a lick. (C) Binned lick rate across all trials for each combination of group, solution, and laser stimulation. (D) Line plot depicting the mean trial lick count for each group (columns), solution (indicated by color), and trial laser stimulation (x-axis, indicated by blue rectangle) (rmANOVA 3-way group x solution x laser interaction, F(2,14)=4.76, *p=0.027). Asterisks indicate results from HSD post hoc test comparing no-stim and stim trials for the solutions as indicated by the color. (E) Perievent time histograms of the difference in GRAB-DA activity between laser stimulation trials and non-stimulated trials at each time point (Δzs) across groups (columns) and solution (rows). Grey rectangle indicates the 3s access period, blue rectangle indicates the 15s laser stimulation window, color indicates striatal region GRAB-DA signal was recorded from, and numerical values indicate number of fibers for each striatal region. (F) Mean Δzs in the early access window during consumption of 1M NaCl (top) or water (bottom). Asterisks indicate results from a Wilcoxon rank sum test (Bonferroni adjusted for multiple comparisons to the same control). (G) 3D map of striatal position and Δzs in the early access window. (H) Correlation between striatal position in the AP (left), ML (middle), and DV (right) dimensions versus the mean Δzs. Text depicts the results of a Pearson’s correlation test. (I-K) Correlation of mean perievent time histogram traces across LHA and striatal signals. (I) Example of data included in the analysis. The mean fiber photometry perievent time histogram over the access window for each solution within each solution/restriction configuration for each region was correlated with every other region. (J) Correlation matrix comparing the mean fiber photometry perievent time histograms during access to each solution across all signals. (K) Line plot depicting the correlation of LHA^GABA^, LHA^Glut^, and LHA^Ratio^ signals with the GRAB-DA signal recorded in each striatal region.

Together, these findings demonstrate that LHA activity influences both consumption behavior and DA dynamics across the striatum. While our data reveals a clear alignment between LHA function, DA signaling, and behavioral responses to different solutions, they do not determine whether changes in DA directly drive behavior or if behavioral modifications influence DA levels. Instead, these results suggest a broader role for LHA^GABA^ and LHA^Glut^ in shaping both striatal DA dynamics and consumption behavior.

We next examined whether the ratio of LHA^GABA^ to LHA^Glut^ activity better predicted striatal DA responses than either population alone. To test this, we computed linear correlations between mean fiber photometry traces for LHA^GABA^, LHA^Glut^, LHA^Ratio^, and striatal DA signals across restriction/solution configurations (**Figure 7I–K**). Visual inspection of the correlation matrix revealed strong correlations among striatal subregions, particularly during FR:Suc and WR:NaCl (**Figure 7J**). Quantifying correlation strength (Pearson’s R) showed that LHA^Ratio^ more closely aligned with DA responses in the ventral striatum under FR:Suc and exhibited broader alignment across the striatum under WR:NaCl. Additionally, correlation patterns followed an anterior-posterior gradient that paralleled the spatial organization observed with optogenetic manipulations: correlations were strongest in anterior regions and declined in more posterior subregions. Taken together, these findings suggest that LHA^Ratio^ represents a distinct dimension of LHA output that is preferentially coupled to striatal DA signals, particularly in anterior and ventral subregions. This highlights how the balance of LHA^GABA^ and LHA^Glut^ activity may serve as a crucial mechanism for fine-tuning striatal DA signaling and consummatory behaviors.

## DISCUSSION

Here, we investigated how lateral hypothalamic LHA^GABA^ and LHA^Glut^ neurons cooperatively regulate DA dynamics across the striatum during consummatory behavior. Using dual-color fiber photometry, we show that LHA^GABA^ and LHA^Glut^ neurons exhibit distinct but complementary activity patterns that scale with solution value and homeostatic demand. Optogenetic manipulations reveal that LHA^GABA^ neurons broadly enhance DA levels across anterior and ventral striatal regions, whereas LHA^Glut^ neurons suppress DA levels in these regions while enhancing DA in the TS. Multi-site fiber photometry demonstrates that DA levels during consumption follow an anterior-posterior gradient, with anterior subregions more strongly encoding solution value and its history, and posterior subregions more strongly encoding task related features (e.g. sensory information) independent of value and consummatory behaviors. Finally, optogenetic inhibition of LHA^GABA^ or LHA^Glut^ neurons bidirectionally alters both consumption behavior and DA dynamics, revealing a cooperative mechanism through which LHA circuits shape motivated consumption, namely that LHA^GABA^ cells promote consumption and are coupled to DA release in a manner positively correlated with the anterior–posterior axis of the striatum, whereas LHA^Glut^ cells inhibit consumption and are coupled to DA release in a negatively correlated manner. Together, these findings provide a refined mechanism of hypothalamic-striatal interactions, where LHA^GABA^ and LHA^Glut^ neurons cooperatively regulate DA dynamics throughout the striatum to influence consummatory behavior. This mechanistic dissection also positions both populations of the LHA as a potential future intervention point in therapeutic manipulations of the DA system to ameliorate maladaptive motivated behaviors.

### LHA^GABA^ and LHA^Glut^ neurons cooperatively represent consumption behavior and regulate DA dynamics throughout the striatum

Previous studies established that LHA^GABA^ neurons promote feeding while LHA^Glut^ neurons suppress it, yet both populations increase their activity during consumption. Here, we demonstrate that under conditions where solution value and homeostatic state elicit a broad range of consumption (FR:Suc and WR:NaCl), LHA^GABA^ and LHA^Glut^ exhibit distinct activity scaling. LHA^GABA^ activity positively correlates with consumption, whereas LHA^Glut^ activity scales negatively, specifically when mice encounter aversive solutions (**Figure 1**). This suggests that these populations function cooperatively to shape adaptive consumption: LHA^GABA^ neurons facilitate greater intake, while LHA^Glut^ neurons selectively *scale* in response to aversive solutions, potentially limiting harmful ingestion. One interpretation is that LHA^GABA^ neurons function as a “universal consumption promoter”, whereas LHA^Glut^ neurons function as a "homeostatic-risk monitor," raising their activity to drive aversion when hypertonic NaCl intake (under WR) could be detrimental. Supporting this, we found that inhibiting LHA^Glut^ neurons enhanced 1.0M NaCl consumption (**Figure 7B-D**). Consistent with previous reports, LHA^Glut^ activity is not limited to aversive solutions, as LHA^Glut^ neurons exhibit increased activity during the onset of consumption regardless of the homeostatic state or solution on offer. This raises the possibility that LHA^Glut^ neurons provide tonic regulation or fine-tuning of LHA^GABA^ output to refine consumption. Previous studies identified LHA neurons whose activity scaled with rewarding or aversive stimuli^71^, but the neurotransmitter identity of these populations was unknown; our findings address this by linking opposing value-scaling signals to genetically defined LHA^GABA^ and LHA^Glut^ neurons. Consistent with this dual involvement, we observed that LHA^GABA^ and LHA^Glut^ jointly modulate DA dynamics across the anterior-posterior axis of the striatum, suggesting that these populations converge on DA circuits to cooperatively guide appropriate intake across varying motivational and homeostatic states. Notably, we focused on fluid consumption tasks that emphasize licking and swallowing; it remains possible that different motor action patterns, such as chewing for solid foods, engage partially distinct LHA circuitry to regulate feeding under different conditions.

The LHA integrates inputs from a broad range of homeostatic, value-related, and taste-processing regions, which could collectively shape the neural dynamics we observed across restriction and solution conditions. For homeostatic signaling, the LHA is downstream of key ingestive centers, including direct projections from hunger promoting agouti-related protein neurons in the arcuate nucleus^72–74^ and thirst promoting neurons in the median preoptic nucleus^75^. Additionally, thirst related signals are conveyed to the LHA from osmolarity sensing neurons of the subfornical organ^76,77^ and organum vasculosum of the lamina terminalis subfornical organ^76^ indirectly through the bed nucleus of the stria terminalis^78^. Value-related modulation arises from descending input from the nucleus accumbens, including D1-expressing neurons that promote feeding^79^. Taste-related signals may reach the LHA through multiple pathways, including a brainstem route via the nucleus of the solitary tract^80^ and parabrachial nucleus^81–83^, as well as a forebrain pathway from the gustatory cortex through the amygdala^84^. The integration of homeostatic and taste-related information by LHA^GABA^ and LHA^Glut^ neurons must be distinct to generate the divergent activity patterns we observed. However, whether and how these two populations directly influence each other remains unclear. Paired electrophysiological recordings indicate that local synaptic connectivity between LHA^GABA^ and LHA^Glut^ neurons is sparse^85^, but alternative pathways may enable cross-regulation. For instance, interactions via the midbrain DA system and reciprocal connections with structures such as the bed nucleus of the stria terminalis, lateral septum, and lateral preoptic area^33^ could provide polysynaptic routes for functional coordination. Additionally, neuropeptide signaling within the LHA may shape intra-LHA activity^17^. Future research should explore how these distinct information streams converge onto the LHA and how its neuronal subpopulations interact through local and polysynaptic mechanisms to regulate motivated behavior.

Multiple lines of evidence support the LHA’s ability to modulate midbrain DA neuron activity. Early studies demonstrated that LHA stimulation supports intracranial self-stimulation in a DA-dependent manner, while more recent work shows that LHA^GABA^ and LHA^Glut^ differentially influence VTA GABA and DA neuron activity^21^. Upregulating LHA^GABA^ decreases VTA GABA activity and increases DA in the NAcCC, whereas LHA^Glut^ stimulation suppresses DA in the NAcCC. Broadly, this descending disynaptic regulation of DA release through midbrain GABA neurons seems to be a common circuit motif shared by other hypothalamic and forebrain populations^86^. However, subsequent work revealed further complexity; for instance, stimulating the LHA^Glut^→VTA pathway increased DA levels in the ventromedial shell (NAcShM) but not the lateral shell (NAcShL), and monosynaptic tracing shows that LHA neurons directly innervate GABAergic, glutamatergic, and DAergic neurons in the VTA that project throughout the striatum. Our results build upon this framework, demonstrating that stimulating LHA^GABA^ neurons produces widespread increases in DA levels, particularly in anterior striatal regions, while stimulating LHA^Glut^ neurons produces more nuanced effects—broad DA suppression in most subregions, mixed effects in some, and robust enhancement in the TS. This partially contrasts with de Jong et al. 2018^87^, as we observed both increased and decreased DA levels in the NAcShL under LHA^Glut^ stimulation, suggesting additional circuit complexity or engagement of alternative pathways (e.g., LHA^Glut^ projections to the lateral habenula^24^). Moreover, LHA^Glut^ stimulation and LHA^GABA^ inhibition both increased DA in the TS, indicating a cooperative influence on this subregion. However, DA levels in the TS did not consistently mirror LHA activity during consumption, suggesting that distinct subsets of LHA neurons, separate from those engaged during consumption, may regulate DA in this subregion. Together, these results establish the TS as a functionally distinct striatal subregion that is differentially modulated by LHA^GABA^ and LHA^Glut^ neurons, reinforcing its unique role within hypothalamic-striatal circuits during consummatory behavior

### Evoked DA release in one striatal subregion minimally propagates to other subregions

Previous studies^55–58^ have proposed that DA levels in the ventral striatum can influence DA release in the dorsal striatum via polysynaptic, reciprocal circuits, forming a theoretical framework for how DA might facilitate the shift from goal-directed to habitual behaviors. However, our findings at least partially challenge this notion, as brief optogenetic stimulation at one striatal subregion did not induce frequency-dependent DA release, aside from DA release in the TS modestly enhancing DA in the DLS (**Figure 3**). Together, these findings suggest that DA release during LHA stimulation (**Figure 2**) and DA scaling during consumption (**Figures 5–7**) likely arise from independent modulation of DA levels across subregions of the striatum, rather than from a cascade in which DA release at one subregion propagates throughout the striatum.

A key distinction between our study and prior work proposing ventral-to-dorsal DA interactions is the time-scale of measurement. Previous studies used pharmacological manipulations and microdialysis, capturing DA release over minutes, which could allow for indirect, slower-acting pathways to coordinate DA levels across regions. By contrast, our second-to-second optogenetic and fiber-photometric approach primarily captures rapid, local changes in DA. Another possibility is that DA release in certain striatal subregions influences release in other subregions under specific behavioral contexts, particularly when accompanied by coincident glutamatergic input (e.g., from the thalamus, ventral hippocampus, basolateral amygdala, or prefrontal cortex), or in subregions we did not examine (e.g., the NAc medial shell). Repeated pairing of evoked DA release may also shape DA levels in other structures. Future work should examine how DA dynamics at longer timescales, across different behavioral states, and within additional striatal regions contribute to the broader coordination of DA release.

### Spatiotemporal dynamics of striatal dopamine levels during consummatory behavior

Our findings reveal that striatal DA responses during consumption follow a posterior-to-anterior progression. Before licking begins, DA levels increase in posterior regions such as the NAcShL, DMS, and DLS on trials with licking compared to those without, coinciding with an early DA spike that does not scale with solution value. This may serve to reinforce the initiation of consumption in response to sensory input (spout extending) or act as a salience signal. Once consummatory behavior diverges for different solutions, DA responses show strong scaling in anterior regions, with the largest effects observed under the WR:NaCl condition. After the access period, DA signals remain distinct according to the preceding trial’s solution in anterior sites (NAcCR, NAcCC, NAcShL). This posterior-to-anterior temporal organization is further supported by shorter DA latencies in posterior compared to anterior regions (**Figure 5 supplement 5**), echoing previous reports of DA waves propagating along this axis during movement and tone presentation^63,64^. Using multi-site photometry, we extend this framework to show that the posterior-to-anterior gradient extends to value representation and sustained motivational signals in anterior/ventral striatal regions. While our experiments focused on the consumption phase, DA release throughout the brain also exhibits post-ingestive responses^88^. Future studies should investigate how feed-forward and feedback ingestive mechanisms^89^ shape DA responses during consumption.

Although dopamine shows widespread scaling during consumption, our computational analyses indicate that different factors contribute to these signals across the anterior-posterior axis. Specifically, our general linear model included predictor sets for “time in trial” (sensory components and trial timing consistent across trials), licking, solution identity/value, and solution history. We observed broad representation of licking across much of the striatum, except in the TS, as well as gradients in the representation of trial-related components (stronger in posterior regions) and solution information (stronger in anterior regions). This aligns with theories positing that posterior/dorsal striatal DA shapes sensorimotor components of behavior, whereas anterior/ventral striatal DA influences value and motivation^90^. Notably, the NAcShL displayed features mirroring both anterior and posterior regions, suggesting it may integrate elements of both movement and value functions.

The TS exhibited DA responses distinct from the broader anterior-posterior gradients observed elsewhere. While most striatal subregions displayed robust scaling tied to solution value and licking behavior, the TS demonstrated only subtle scaling and weaker representations of both licking and solution information. Optogenetic manipulations further highlighted its uniqueness: inhibiting LHA^GABA^ or activating LHA^Glut^ increased DA in the TS, in contrast to the opposite effects seen in most other striatal regions.

Correlation analyses also underscored its distinct profile, as DA levels in the TS showed substantially lower correlations with other striatal areas—particularly during consumption, when DA levels show clear scaling elsewhere in the striatum. These functional results may reflect the unique set of inputs onto TS-projecting DA neurons^34^. Intriguingly, our optogenetic data suggest that these TS-projecting DA neurons receive direct innervation from LHA^GABA^ and LHA^Glut^ populations rather than relying on intermediary GABAergic relays that are evident with DA populations projecting to the anterior/ventral and posterior/dorsal striatum. The difference in coupling between descending GABA and glutamate regulation of TS-projecting DA neurons could partially explain the reduced correlation between DA levels in the TS and the rest of the striatum during consumption. More broadly, the TS’s differential response pattern emphasizes its role as a specialized node within the striatal network, with a unique function in shaping behavior.

Midbrain DA neurons projecting to distinct downstream targets receive largely similar monosynaptic inputs, with the notable exception of TS-projecting DA neurons^34^. Prior to our study, it was unclear if this difference in connectivity was related to differences in spontaneous DA dynamics. Our study enabled us to simultaneously record DA levels throughout six striatal subregions and assess the correlation of DA levels between them to determine if TS DA levels are functionally distinct. We found that DA levels were positively correlated throughout the striatum, and the degree of correlation was dependent on the Euclidean distance between recording sites—subregions closer in space exhibited stronger correlations in DA dynamics (**Figure 4**). However, during consumption, the correlation of DA levels in the TS with other subregions was substantially lower than correlations observed across all other striatal sites. Notably, during consumption, correlations in DA levels were enhanced across most of the striatum but remained low between the TS and other regions (**Figure 5 supplement 6**). These results suggest that during consumption—especially of aversive solutions—inputs onto the DA system produce coherent dynamics in DA levels that span the striatum while largely excluding the TS.

It remains unclear whether distinct representations in DA dynamics across the striatum arise from the same or different DA neurons, and whether the behavioral function of DA during consumption also follows this spatial gradient. One possibility is that unique subpopulations of DA neurons encode different task components, with distinct genetic profiles or input patterns shaping their representation of consummatory behavior. Recording individual DA cell bodies in the midbrain^67,91^ or individual axons in the striatum^92^ would help determine whether these representations reflect functionally distinct DA populations. Alternatively, differences in DA signals across the striatum could arise from local presynaptic mechanisms regulating DA release or from differential post-release modulation, such as through acetylcholine-dependent signaling. Future studies should examine presynaptic release dynamics and local regulatory mechanisms to determine whether these processes contribute to the DA representations observed during consumption. Additionally, causal manipulations of DA at these timescales across different striatal regions will be necessary to establish whether its influence on consumption behaviors follows this organizational gradient.

The role of striatal DA in consummatory behavior is complex and multifaceted serving unique functions across multiple time-scales^1,93,94^. While we observe widespread scaling of DA levels across the striatum during consumption, this does not necessarily imply that DA in each of these structures directly drives consummatory behavior. Instead, DA levels likely shape consumption through different functions dependent on the striatal subregion. In the ventral striatum, DA may contribute to incentive sensitization, motivation, and reinforcement learning, processes that evolve over longer timescales and may only become evident through repeated experience or learning-based manipulations. In contrast, DA levels in the dorsal striatum may serve to reinforce specific action sequences, including the initiation of consumption. Future studies should continue to probe the regional function of DA during consumption in shaping behavior by systematically manipulating DA levels at discrete time points during consumption in behavioral assays designed to model the wide range of DA functions.

### Limitations, considerations, and future directions

Our findings reveal a strong coupling between LHA neuronal activity and spatially organized DA dynamics during consumption. However, it is important to note that GRAB-DA photometry signals do not provide a direct readout of midbrain DA neuron spiking or absolute DA concentrations. Prior studies have shown that VTA firing rates do not always predict ventral striatal DA levels^95^, highlighting the role of presynaptic mechanisms in shaping DA signals^96,97^. Additionally, differences in DA kinetics across striatal subregions^61,98^ could contribute to the sustained DA elevations observed in anterior regions (NAcCR, NAcCC, and NAcShM) under the WR:NaCl condition. These prolonged signals may reflect extended tonic firing of DA neurons or unique presynaptic adaptations of DA terminals in response to aversive solutions or broad value distributions. Finally, while fiber photometry with GRAB-DA provides high-temporal-resolution recordings of DA fluctuations, it may not fully capture slower DA changes that occur across prolonged behavioral states. Techniques such as microdialysis^95^ or fluorescence lifetime imaging^99^ could help resolve tonic DA shifts and their impact on long-term motivational states.

An important consideration of our study is the use of head-fixation and randomized solution access, which attempts to isolate consummatory behavior by eliminating approach behaviors and minimizing learning-related influences and normalizing post-ingestive factors^88^. While this design enables precise measurement of LHA^GABA^ and LHA^Glut^ activity and striatum-wide DA levels during consumption, it excludes key processes such as decision-making, motivation, and experience-dependent adaptations. To gain deeper insights into how the LHA and striatum-wide DA dynamics contribute to naturalistic behaviors, future studies should systematically build upon these findings by incorporating predictive cues, approach behaviors, and operant conditioning. These approaches would provide a more comprehensive understanding of how LHA-striatal circuits shape DA dynamics and regulate adaptive feeding behaviors across different motivational contexts.

## Conclusions

The goal of this study was to test whether LHA circuits influence striatal dopamine signaling during consumption-related behavior. We found that the activity ratio of LHA^GABA^ to LHA^Glut^ neurons dynamically scales with the value and valence of consumed solutions (**Figure 1**), and that both populations exert time-locked effects on striatal dopamine release (**Figures 2, 7**) that do not rely on dopamine propagation between subregions. Instead, dopamine levels appear to be independently modulated across spatially organized striatal subregions along the anterior-posterior axis, each of which exhibits semi-discrete representations of task variables (**Figures 3–6**). This work provides a circuit mechanism for lateral hypothalamic modulation of the DA system during consumption behaviors that can be leveraged in future work to intervene when this circuitry goes awry and produces maladaptive motivation and consummatory behaviors.

## Lead contact

Further information and requests for resources and reagents should be directed to and will be fulfilled by the lead contact, Garret D. Stuber.

## Data and code availability

- All original code has been deposited in a public reposition at GitHub [https://github.com/agordonfennell/gordon-fennell_et.al._2025] and is publicly available as of the date of publication.
- All data will be made available at reasonable request.
- Any additional information required to reanalyze the data reported in this paper is available from the lead contact upon reasonable request.

## ACKNOWLEDGMENTS

This work was supported by R37DA032750 (G.D.S.), R01DA038168 (G.D.S.), R21DA050868 (G.D.S.), P30DA048736 (G.D.S.), NSF DMS-2322920 (D.M.), and Office of Naval Research N000142312589 (Program in Mathematical Data Science, D.W.). A.G.F. was supported by K99DA059612, F32DA054719, T32DA7278-27. M.M.H. was supported by F31DA053706.

## AUTHOR CONTRIBUTIONS

Conceptualization, A.G.F. and G.D.S.; methodology, A.G.F., M.M.H., G.D.S.; Investigation, A.G.F., B.M.B., J.M.B., I.M., A.C., H.S., E.L., M.M.H., and M.C.; writing—original draft, A.G.F. and G.D.S.; writing—review & editing, D.W., E.A., M.M.H., I.M., and H.P.S.; funding acquisition, A.G.F. and G.D.S.

## DECLARATION OF INTERESTS

All authors declare no competing interests.

**Figure 2 Supplement 1:**
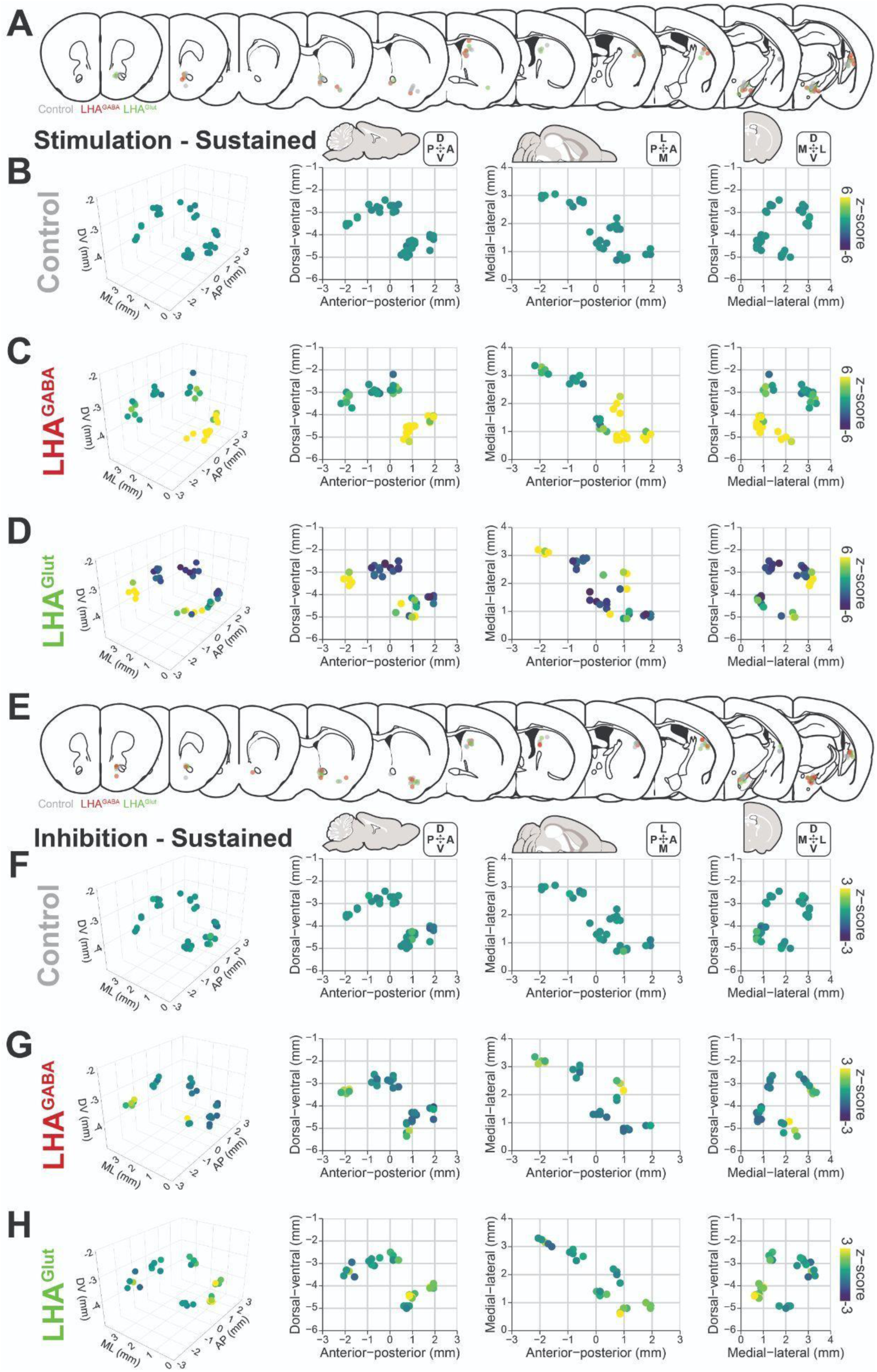
Histological analysis of LHA^GABA^ and LHA^Glut^ regulation of striatal DA release. (A-D) Histological analysis of striatal DA response to optogenetic *stimulation* of LHA^GABA^ or LHA^Glut^ neurons during the sustained windows defined in Figure 2. (A) Coronal atlas sections depicting the location of fiber placements in the striatum to record GRAB-DA signals and in the lateral hypothalamus to regulate the activity of LHA neurons. Color depicts the experimental group (Grey: control, Red: LHA^GABA^, Green: LHA^Glut^). (B-D) from left to right: representation of striatal fiber placement in 3d, sagittal, horizontal, and coronal orientations across the control (B), LHA^GABA^ (C), and LHA^Glut^ (D) groups. Color depicts the mean z-score response. Fiber positions have been jittered by 0.1mm to reduce overlap. (E-H) Histological analysis of striatal DA response to optogenetic *inhibition* of LHA^GABA^ or LHA^Glut^ neurons. Figure annotations and statistics follow convention in corresponding panels in (A-D).

**Figure 2 Supplement 2:**
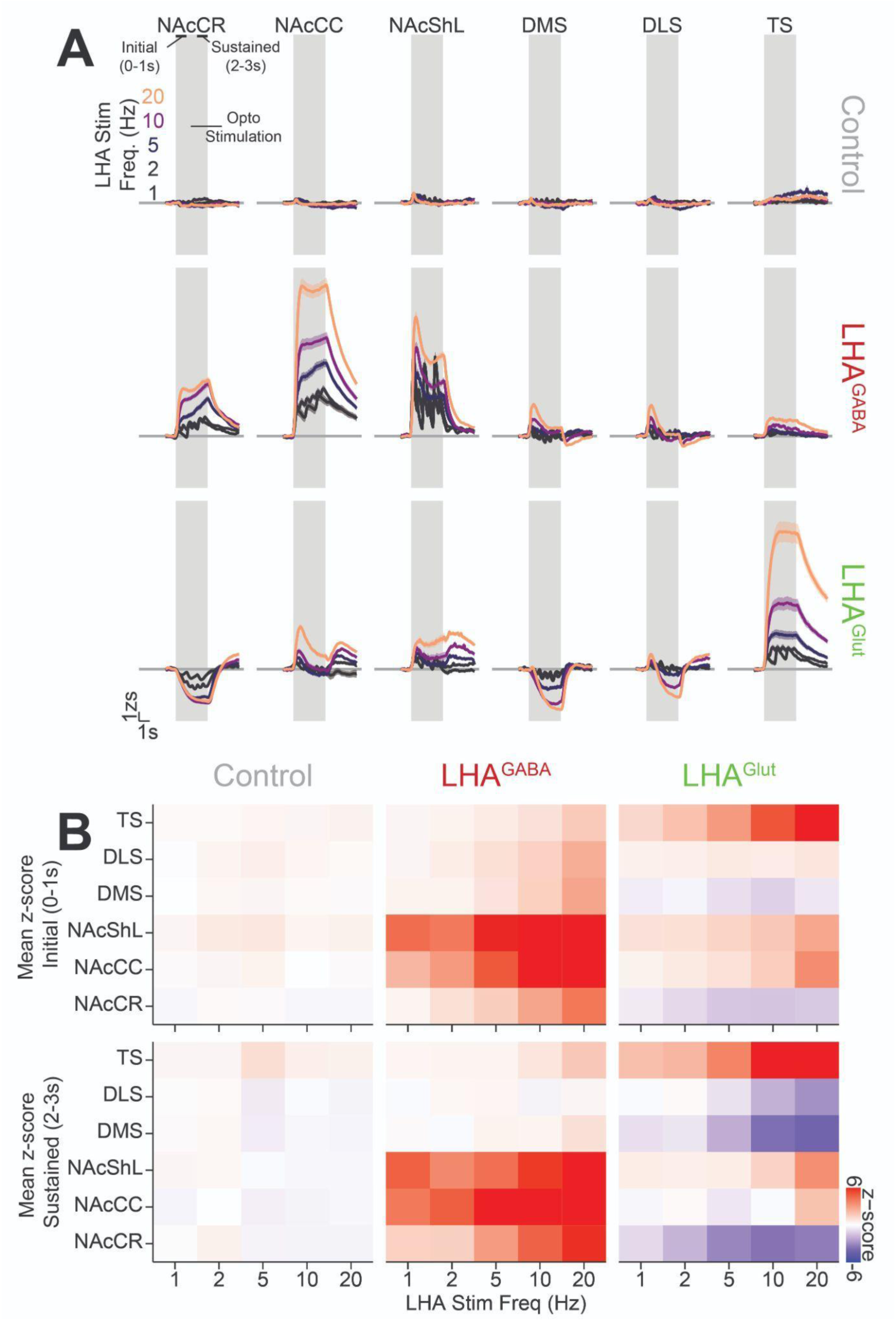
Frequency dependence of LHA^GABA^ and LHA^Glut^ regulation of striatal DA release. (A) Striatal GRAB-DA response to optogenetic stimulation of LHA^GABA^ or LHA^Glut^ neurons at a range of frequencies. Horizontal facets depict striatal region, vertical facets depict experimental group, and color depicts laser stimulation frequency. (B) Heat map displaying the mean z-score response. Horizontal facets depict experimental group, vertical facets depict time-window of mean, and color depicts mean z-score.

**Figure 3 Supplement 1:**
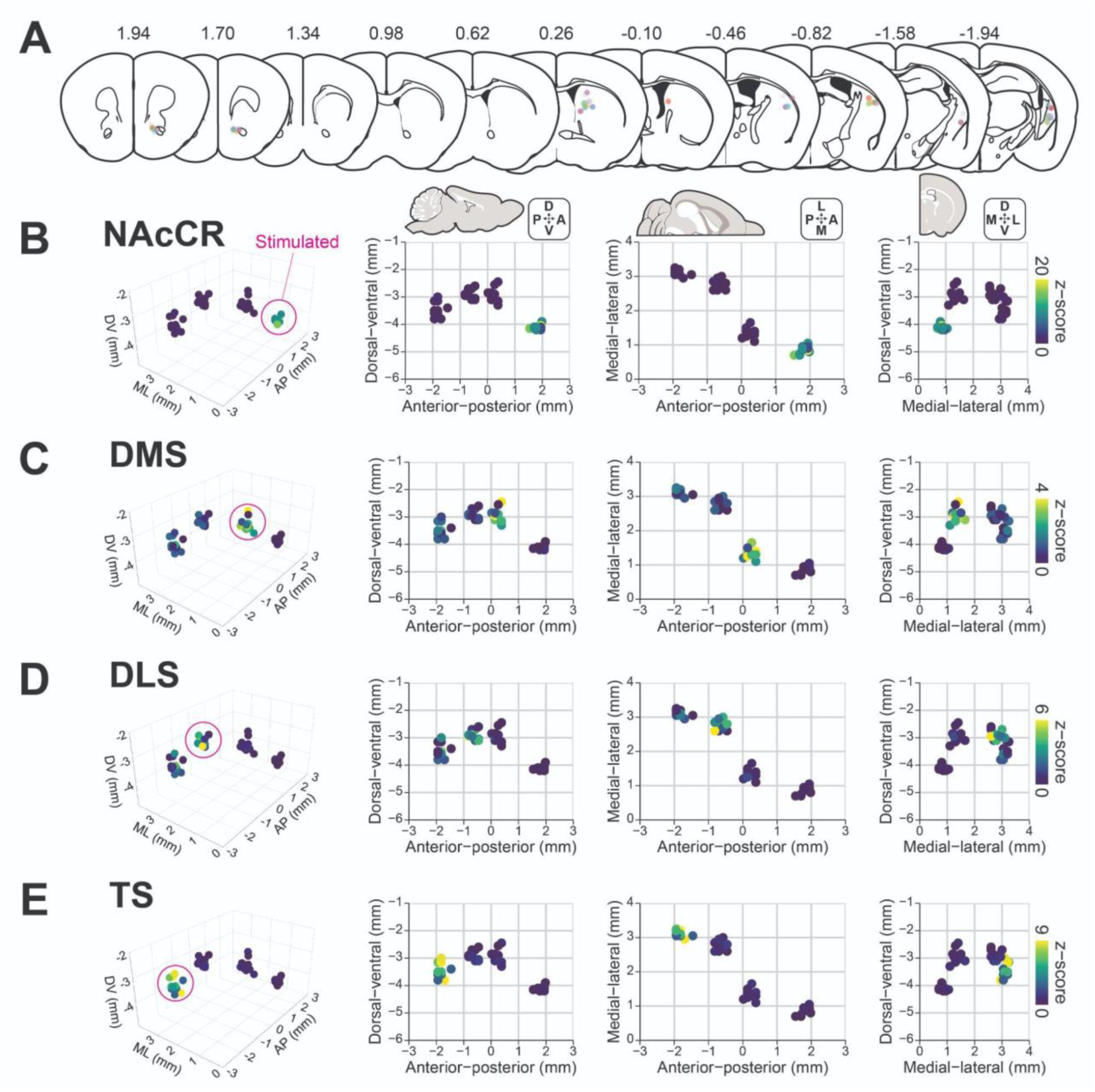
Histological analysis of cross-regulation of DA release between striatal regions. (A) Coronal atlas sections depicting the location of fiber placements in the striatum to record GRAB-DA signals and optogenetically stimulate DA release. Color depicts subject identity. (B-E) from left to right: representation of striatal fiber placement in 3d, sagittal, horizontal, and coronal orientations. Rows depict the striatal subregion in which DA terminals were stimulated: NAcCR (B), DMS (C), DLS (D), and TS (E). Color depicts the mean z-score response. Fiber positions have been jittered by 0.1mm to reduce overlap.

**Figure 4 Supplement 1:**
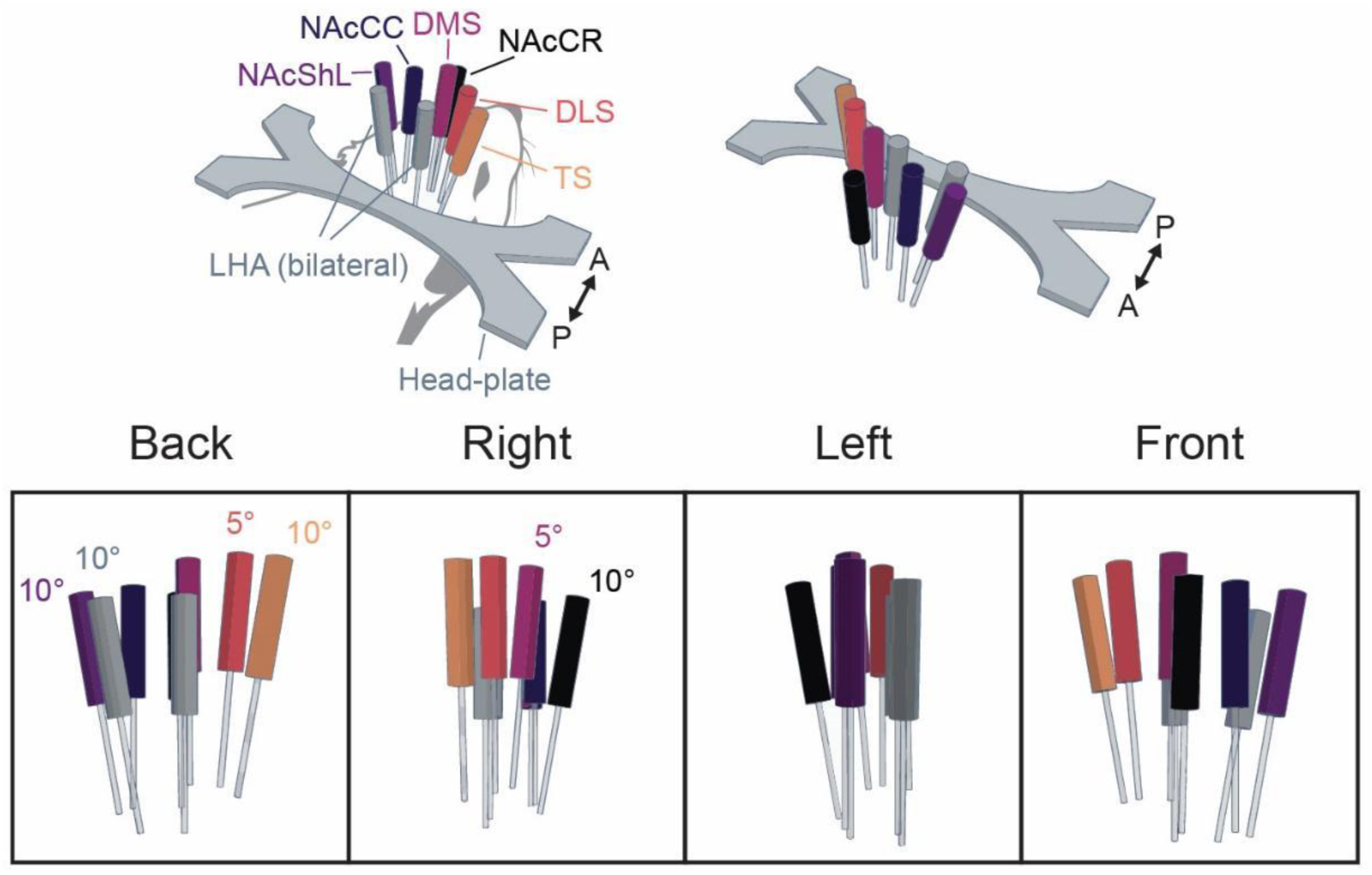
Design of multi-fiber approach. 3d diagram of the 8-fiber implant configuration allowing for simultaneous optogenetic regulation of the LHA bilaterally and recording of DA responses in 6 striatal subregions.

**Figure 4 Supplement 2:**
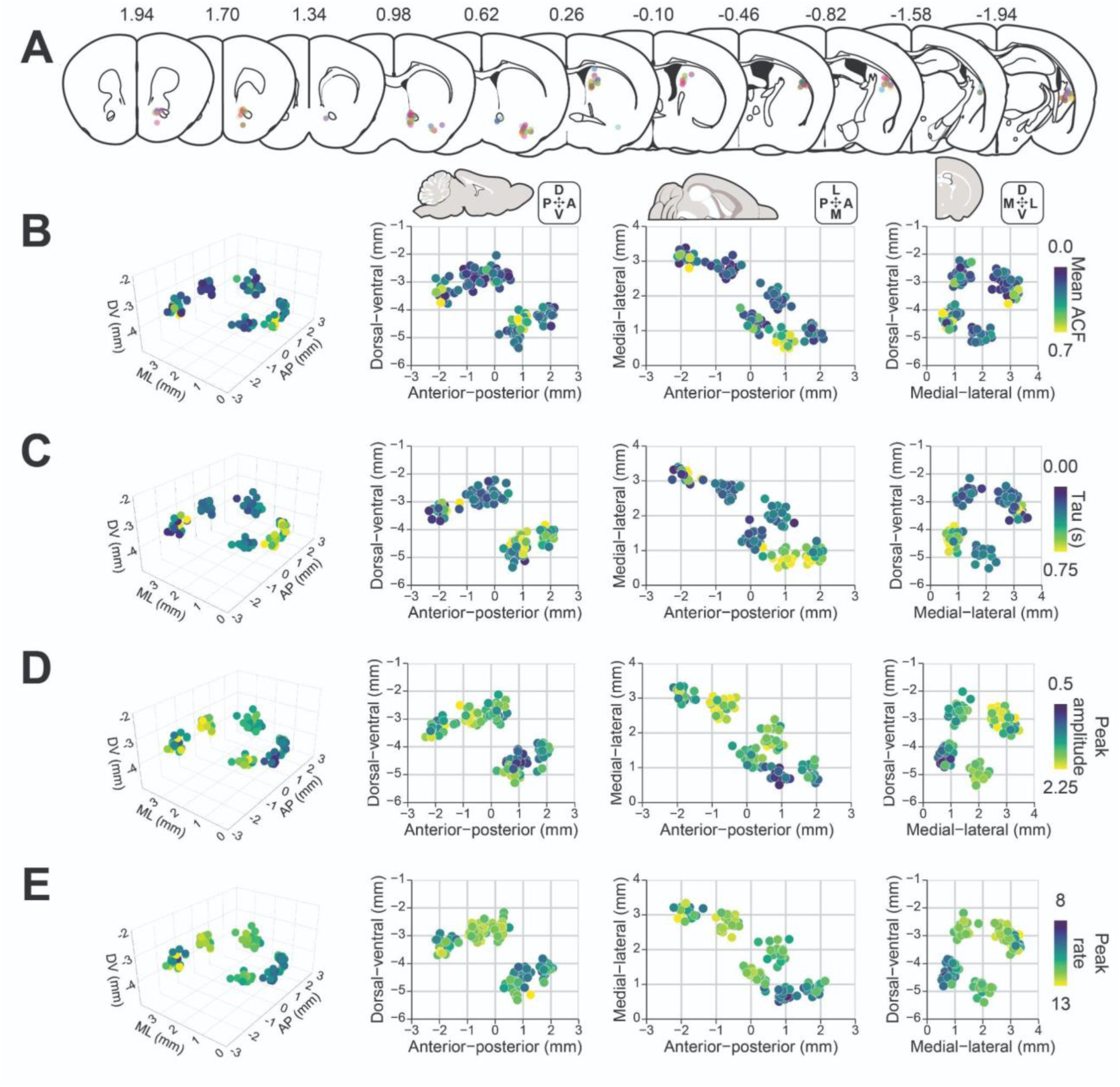
Histological analysis of spontaneous DA dynamics across the striatum. (A) Coronal atlas sections depicting the location of fiber placements in the striatum to record GRAB-DA signals. Color depicts subject identity. (B-E) from left to right: representation of striatal fiber placement in 3d, sagittal, horizontal, and coronal orientations. Rows depict the summary variable in Figure 4 used as the color range: Mean ACF over 0-5 seconds (B), tau estimate (C), peak amplitude of transient (D), and spontaneous peak rate per 10s (E). Fiber positions have been jittered by 0.1mm to reduce overlap.

**Figure 4 Supplement 3:**
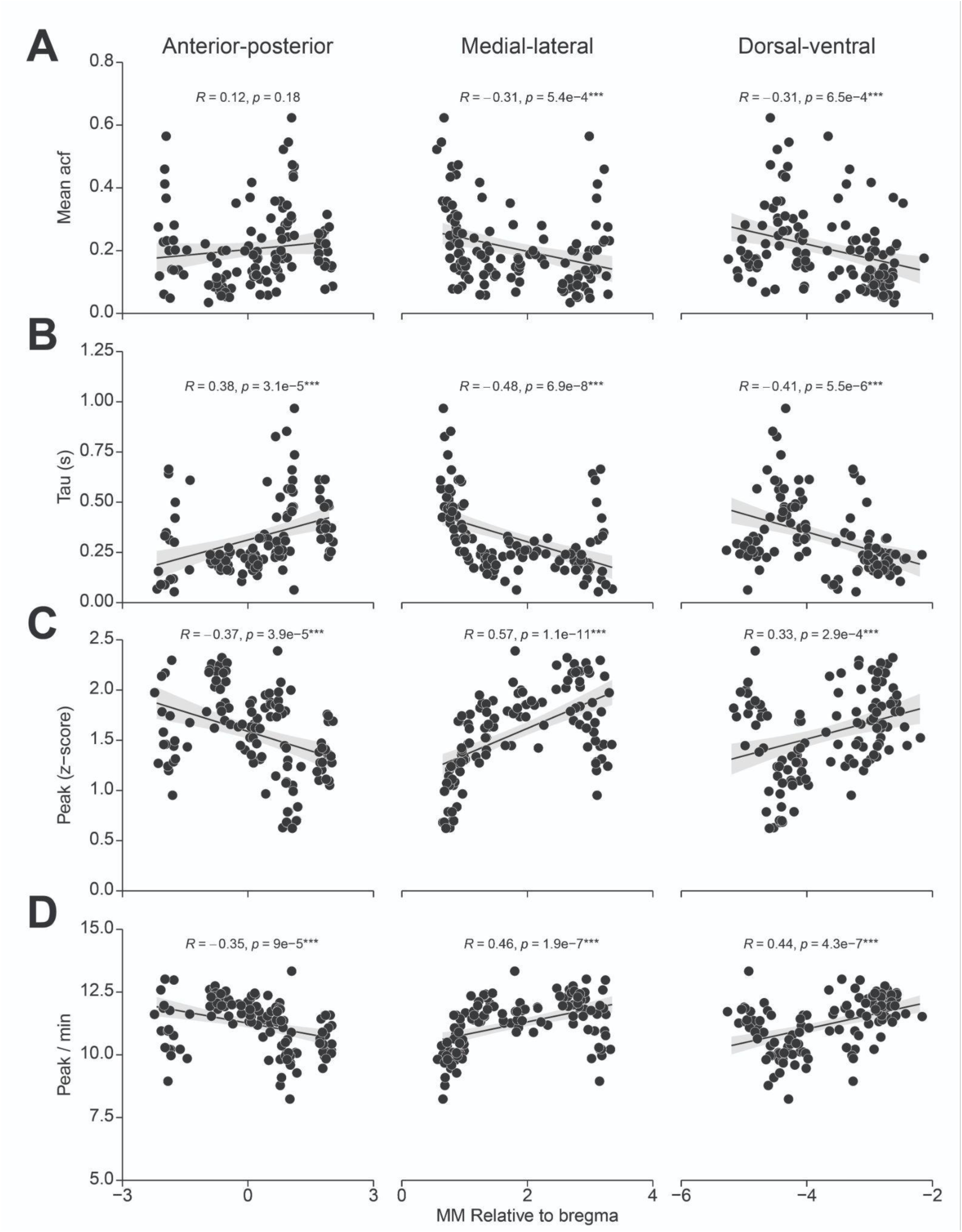
Correlation between atlas axes and spontaneous DA dynamics across the striatum. Correlation between the position in anterior-posterior (left), medial-lateral (middle), and dorsal-ventral (right) axes of the mouse brain against the summary variables in Figure 4: Mean ACF over 0-5 seconds (B), tau estimate (C), peak amplitude of transient (D), and spontaneous peak rate per 10s (E). Text depicts the results of a Pearson’s correlation test. Fiber positions have been jittered by 0.1mm to reduce overlap.

**Figure 5 Supplement 1:**
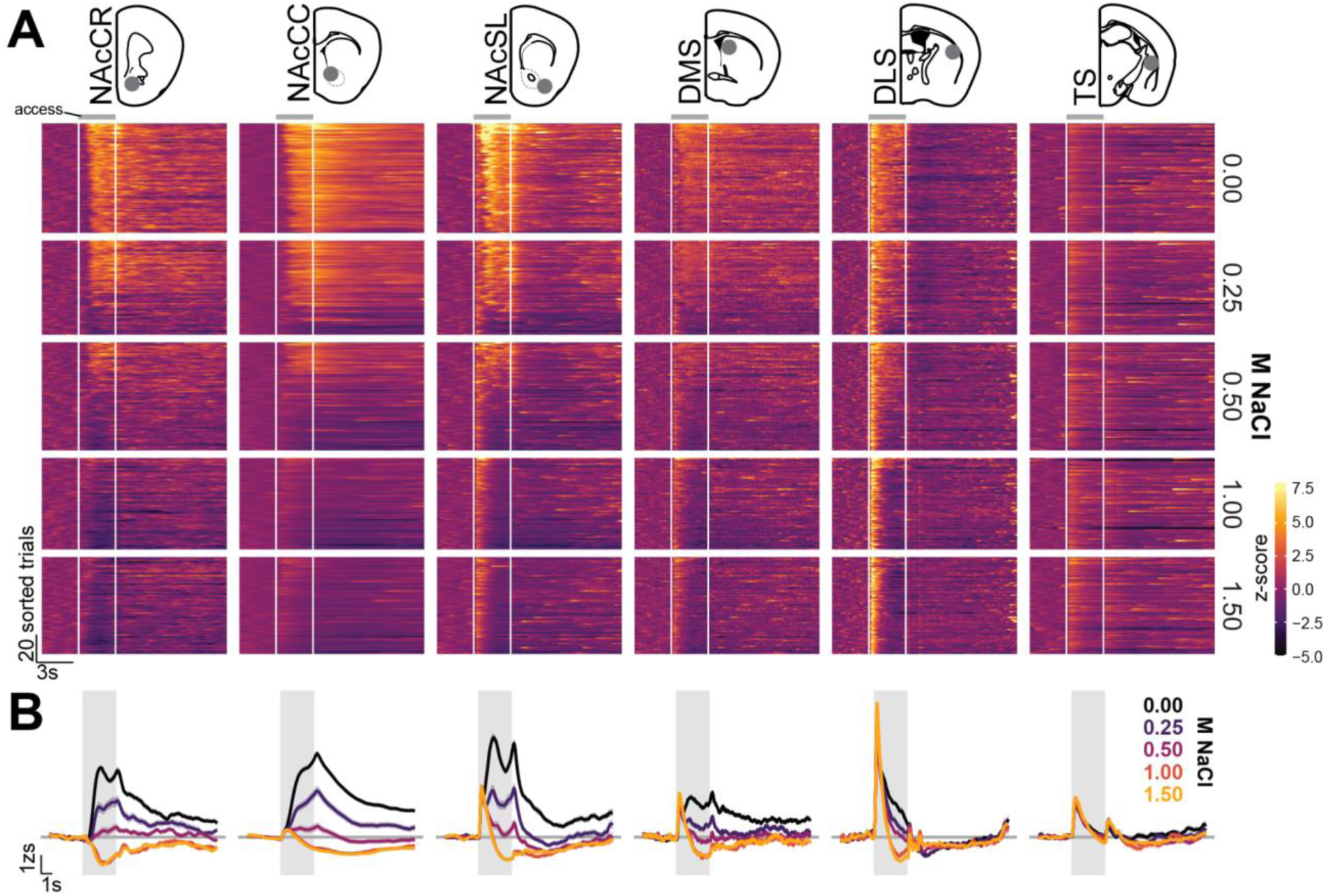
Representative subject DA dynamics across the striatum during consumption. GRAB-DA dynamics from a representative subject across 4 sessions of the brief-access taste task under the WR:NaCl configuration (same subject as Figure 4A**, C**). (A) heat-maps displaying the GRAB-DA dynamics across all trials faceted vertically by solution concentration and horizontally by brain area. Trials were sorted based on the global mean z-score during the access period for each solution individually– each row represents a single trial across all striatal subregions. (B) Perievent time histograms displaying the mean GRAB-DA response during access to each solution concentration (indicated by color) faceted horizontally by brain region. Solid line depicts mean and shaded line depicts SEM calculated across all trials.

**Figure 5 Supplement 2:**
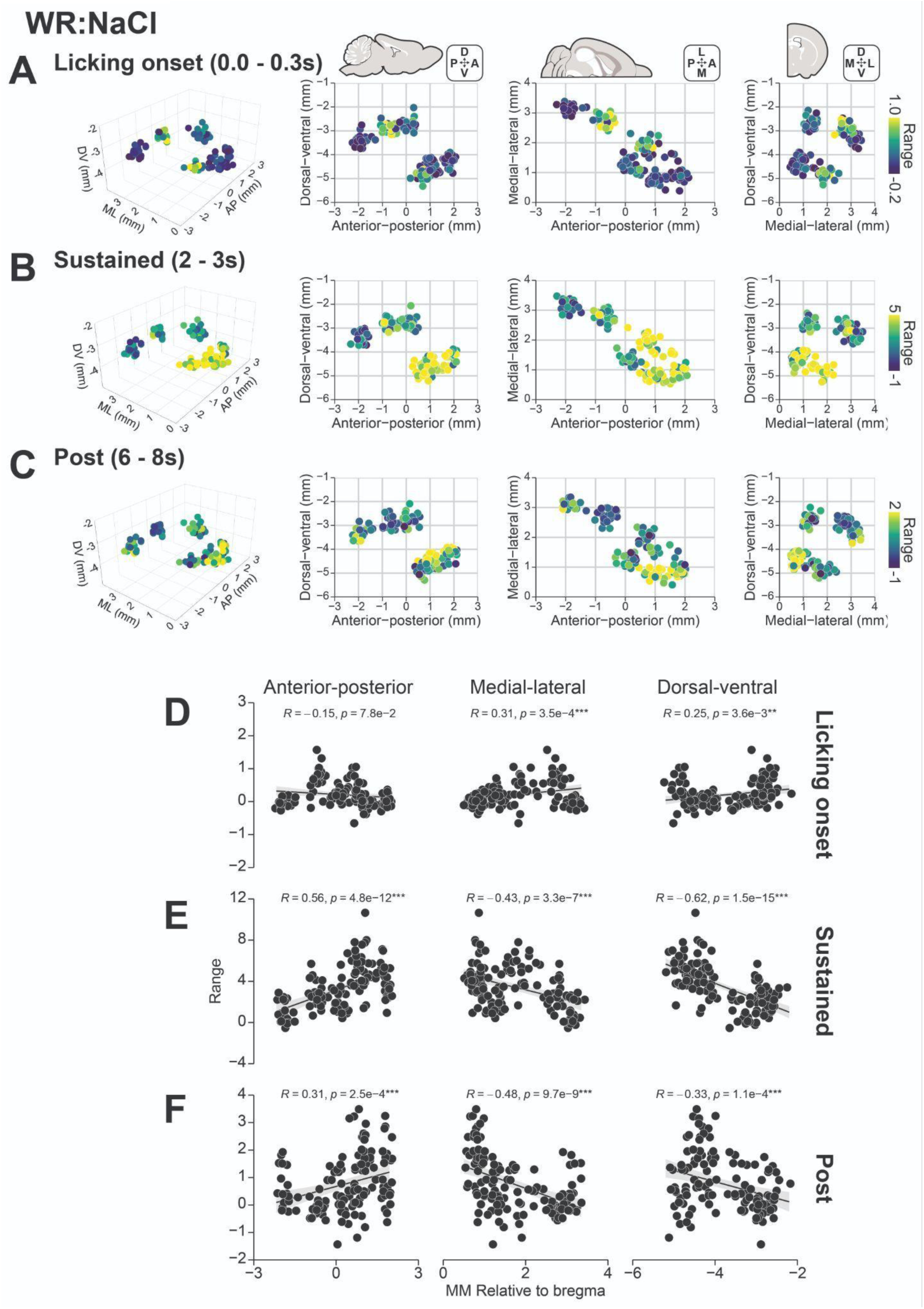
Histological analysis of DA dynamics across the striatum during consumption. (A-C) from left to right: representation of striatal fiber placement in 3d, sagittal, horizontal, and coronal orientations. Rows depict the GRAB-DA mean response range under the WR:NaCl configuration during different time-windows as defined in Figure 5 used as the color range: Licking onset (A), sustained consumption (B), and post consumption (C). Fiber positions have been jittered by 0.1mm to reduce overlap. (D-F) Correlation between the position in anterior-posterior (left), medial-lateral (middle), and dorsal-ventral (right) axes of the mouse brain against the GRAB-DA response range during different time-windows as defined in Figure 5: Licking onset (D), sustained consumption (E), and post consumption (F). Text depicts the results of a Pearson’s correlation test. Fiber positions have been jittered by 0.1mm to reduce overlap.

**Figure 5 Supplement 3:**
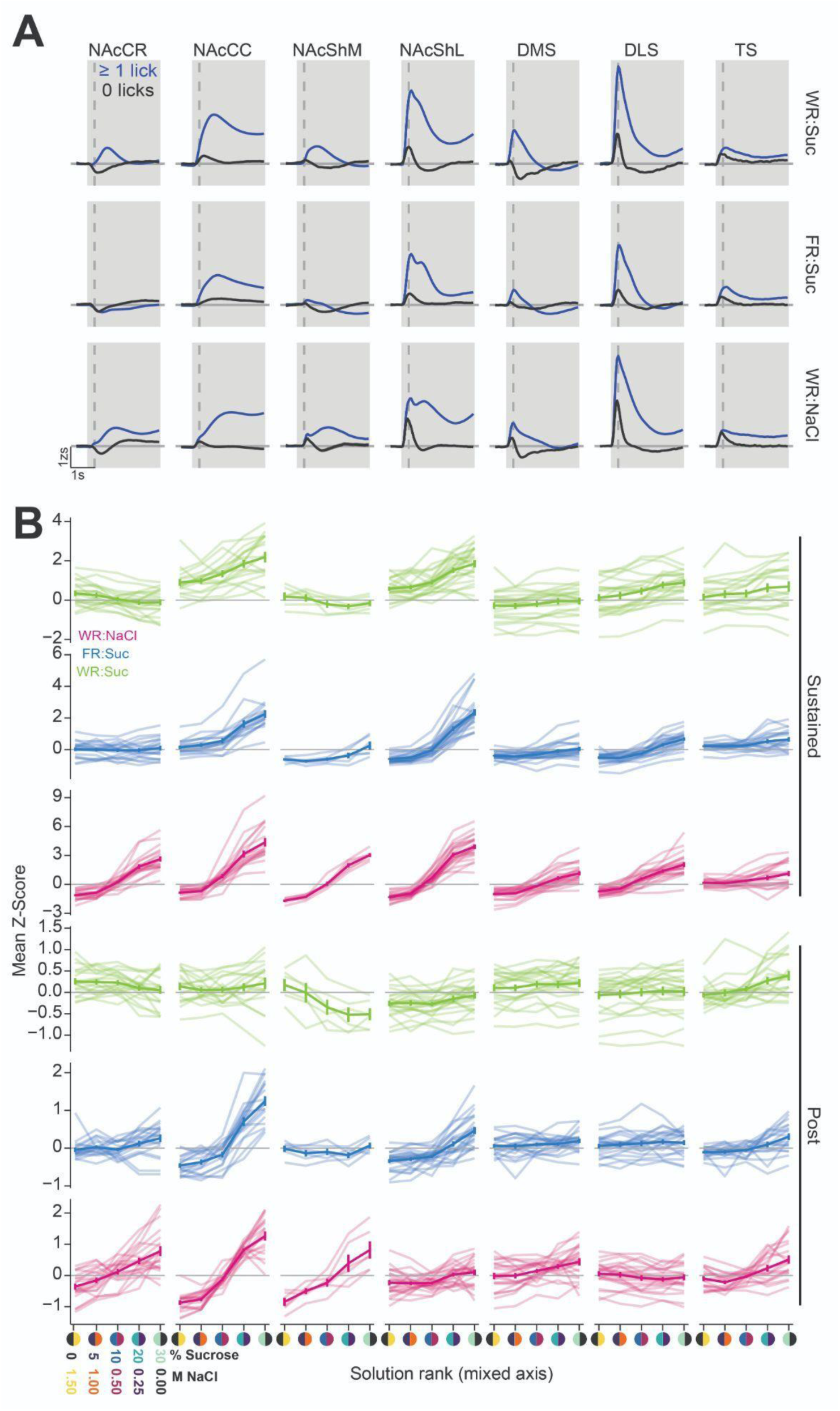
of response dynamics DA dynamics across the striatum during consumption. (A) Mean GRAB-DA trace for trials with and without licking across all striatal subregions and restriction/solution configurations. Solid line depicts mean and shaded line depicts SEM calculated across all corresponding trials. (B) Single subject representations of data shown in Figure 5H, K. Each faded line depicts the mean response for a single subject, dark lines indicate mean and SEM calculated across subjects.

**Figure 5 Supplement 4:**
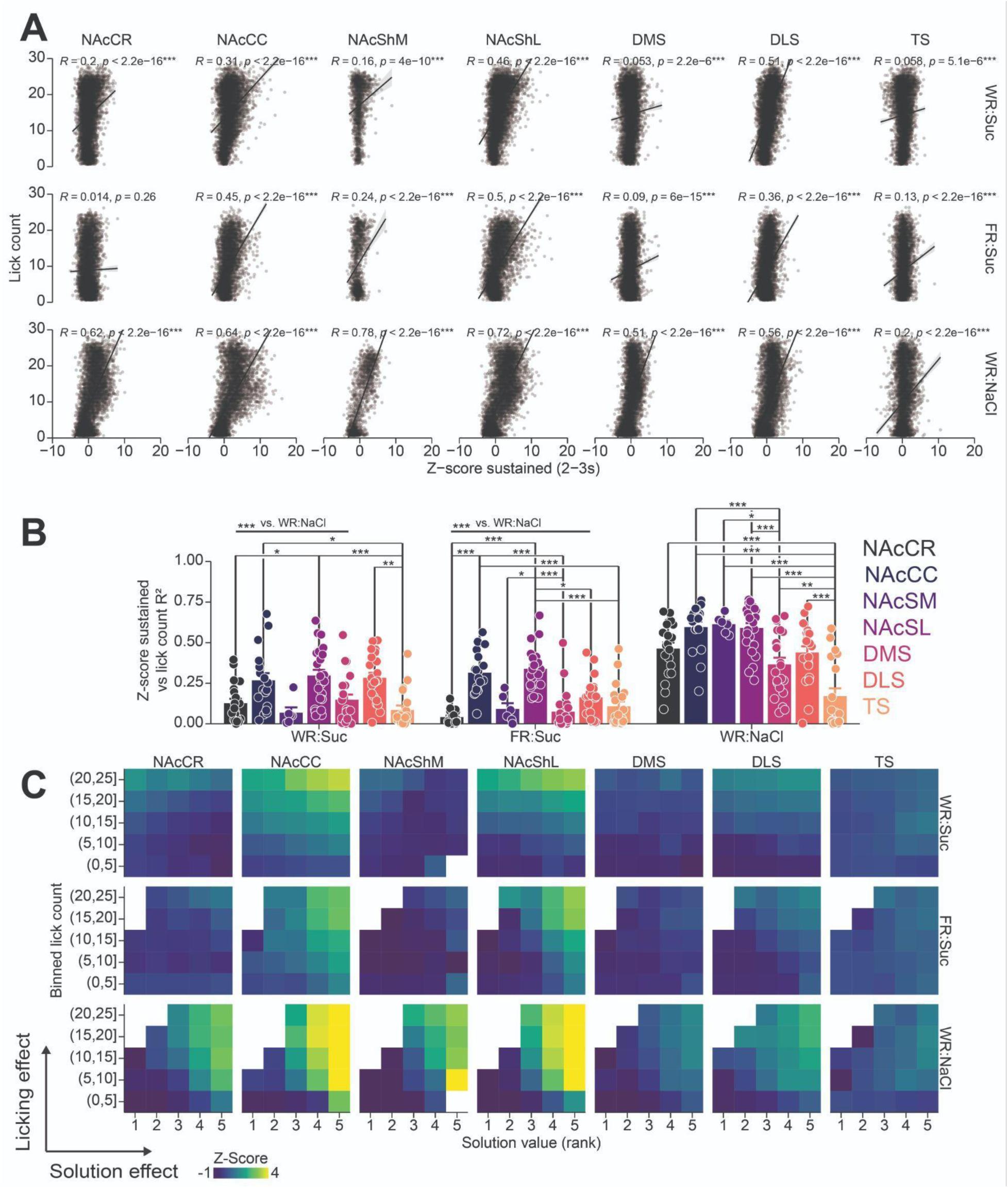
Striatal DA varies with licking and solution rank. (A) Correlation between the mean GRAB-DA photometry signal during the sustained time-window and the number of licks on the corresponding trial. Plots are faceted by striatal subregions horizontally and restriction/solution configuration vertically. Lick count capped at 30 for display purposes. Text depicts the results of a Pearson’s correlation test. (B) Comparison of the correlation strength for each fiber calculated individually. R^2^ was calculated using a Pearson’s correlation test computed on the mean GRAB-DA photometry signal during the sustained time-window and the number of licks on the corresponding trial. Asterisks denote significant HSD post hoc comparisons. (C) Heatmap displaying the mean GRAB-DA photometry signal during the sustained time-window separated by the solution (x-axis) and the binned number of licks (y-axis) on the corresponding trial. Plots are faceted by striatal subregion horizontally and restriction/solution configuration vertically. Data was filtered on a subject basis to combinations of solution rank and binned lick counts with greater than 4 trials per combination.

**Figure 5 Supplement 5:**
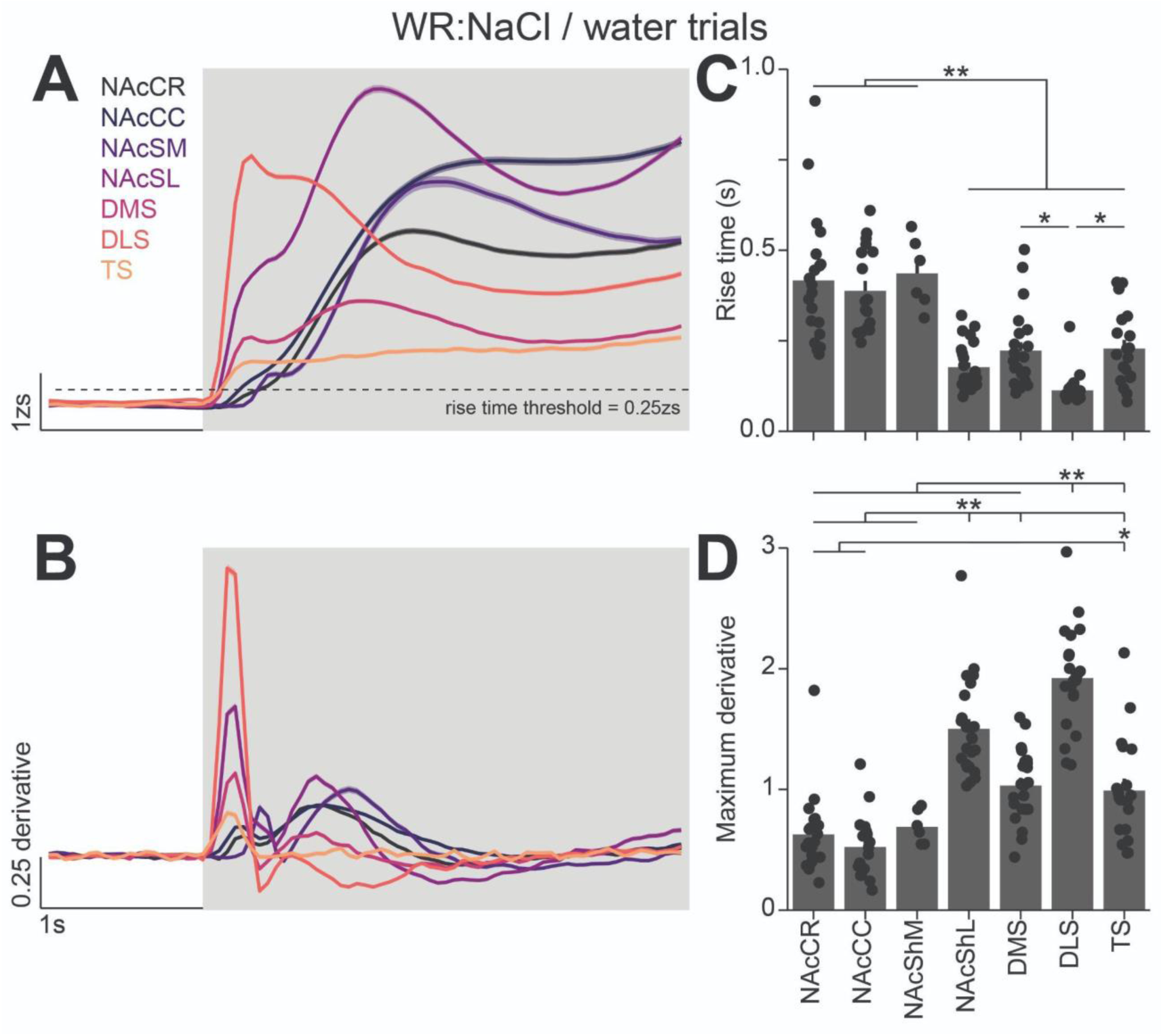
Onset kinetics of GRAB-DA photometry signals across the striatum during consumption. (A-B) Perievent time histograms of GRAB-DA photometry signals (A) and the derivative (B) during consumption of water under the WR:NaCl restriction/solution configuration. Color indicates the striatal subregion the signal was recorded from, solid line depicts mean and shaded line depicts SEM calculated across all corresponding trials. (C-D) Comparison of the rise time (C, time to reach a z-score of 0.25, ANOVA, region effect, F(6,124)=23.79, p=1.41e-18) and maximum derivative during the access window (D, ANOVA, region effect, F(6,124)=37.86, p=8.06e-26). Bar plot depicts the mean value for each striatal region (bar depicts mean, error bar depicts SEM, and individual points depict single striatal sites).

**Figure 5 Supplement 6:**
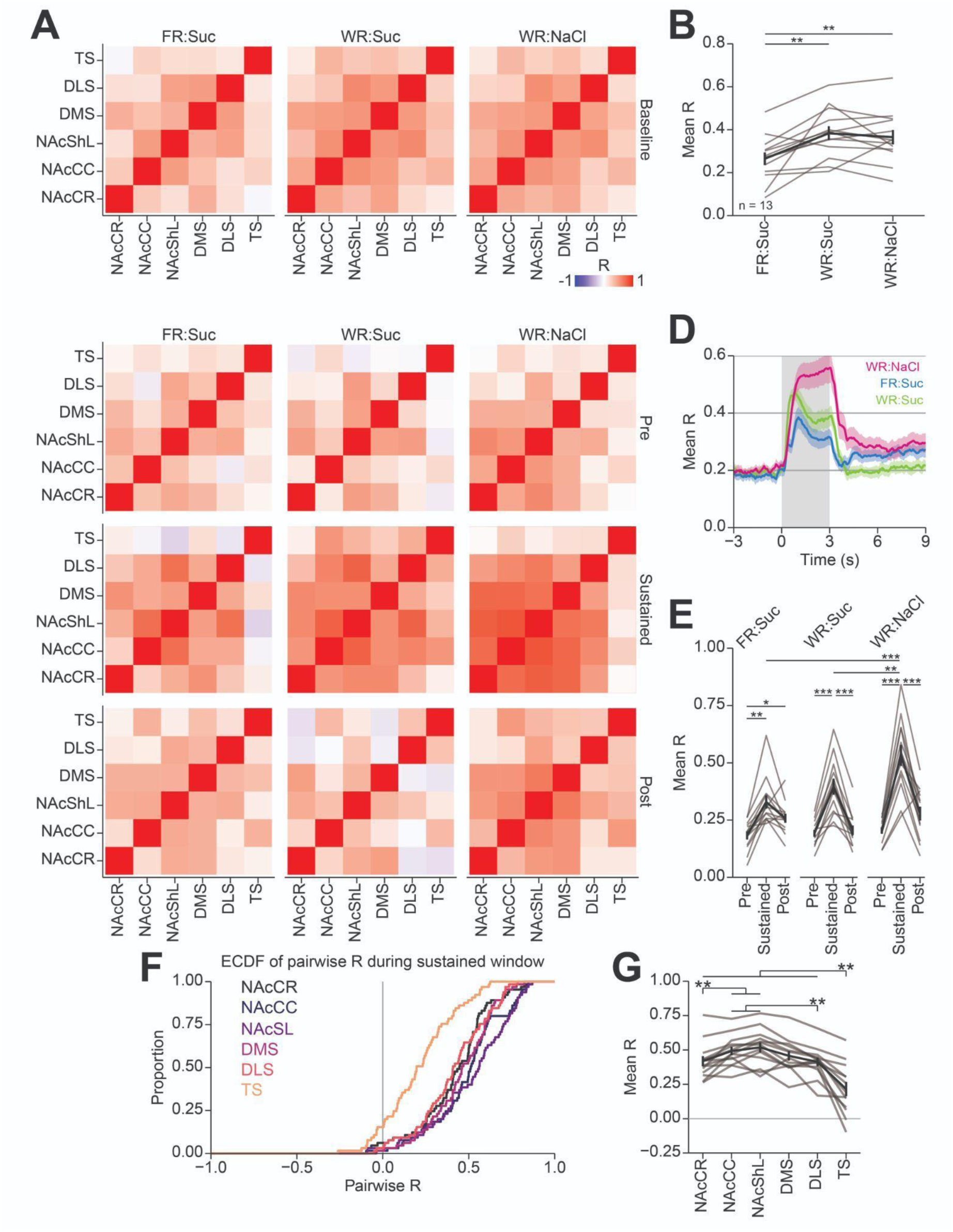
Differential correlations of striatum-wide DA dynamics during consumption behavior across homeostatic demand and solution sets. Analysis of the correlation of GRAB-DA levels across striatal regions in mice with six accurately placed fibers (n=13). (A-B) Correlation of GRAB-DA levels during a 2-minute period prior to session start indicating heightened correlation under water-restriction compared to food-restriction. (A) Correlation matrix with color depicting Pearson’s R for each pair of striatal subregions. (B) Mean Pearson’s R across all striatal regions. (C-H) Correlation of GRAB-DA levels during consumption in the brief-access taste task. (C) Correlation matrices with color depicting Pearson’s R of GRAB-DA levels between each region in 2 s windows prior (“Pre”, -2-0s relative to access start; top row), during consumption (“Sustained”, 1-3s relative to access start; middle row), and following (“Post”, 3-5s following access end; bottom row) the access period under each restriction/solution configuration (columns). (D) Mean Pearson’s R over the peri-access time window showing increased correlation of DA levels during the access period for all configurations. (E) Mean Pearson’s R over each 3 s time window. (F) Empirical cumulative distribution of the pairwise correlation of DA levels during the sustained window across all restriction/solution configurations. (G) Line plot of the mean correlation of each region with all other regions (left). Across all panels, faded lines depict individual subjects, black lines depict mean across subjects, error bars / ribbons depict SEM, asterisks denote results from HSD post hoc tests using the following convention: *p < 0.05, **p < 0.01, and ***p < 0.01.

**Figure 6 Supplement 1:**
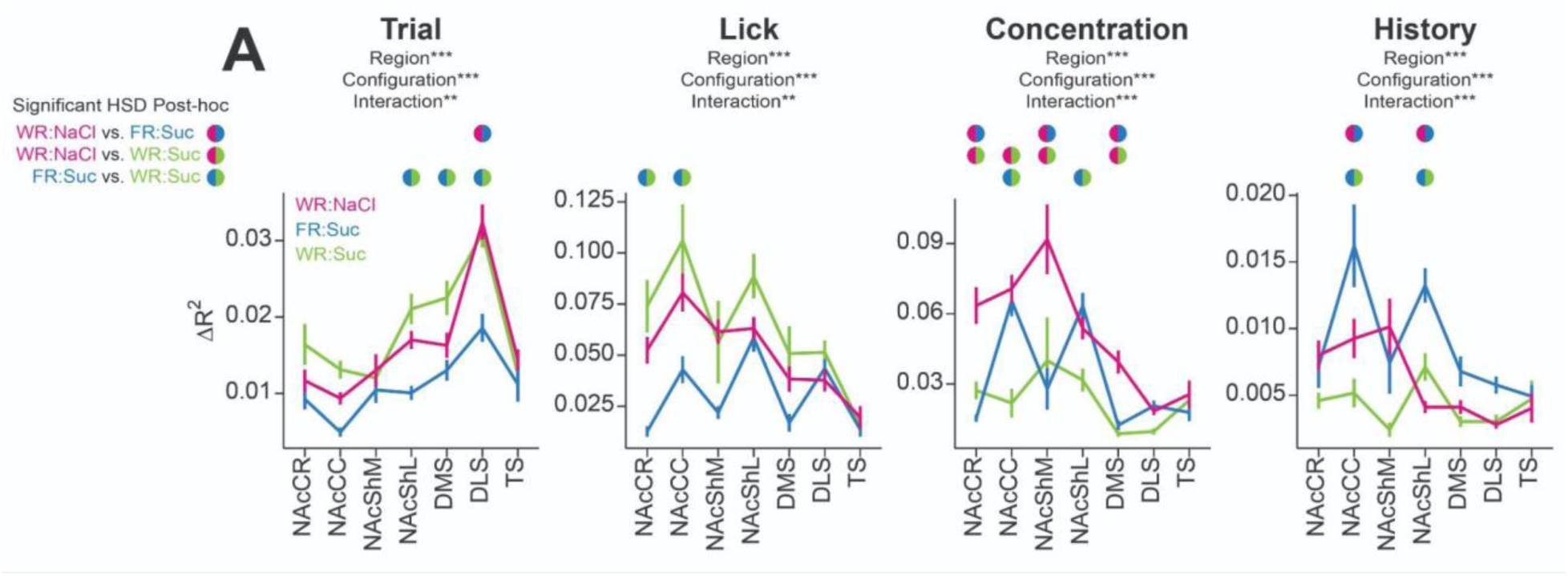
Unique representations of consumption behavior across homeostatic demand and solution set. (A) Change in GRAB-DA signal prediction (ΔR^2^) across striatal subregions with removal of each predictor set (columns) for all three restriction/solution configurations (depicted by color). Significant effects of region, configuration, and region / configuration interaction on ΔR^2^ were observed for all predictor sets. Significant HSD post hoc comparisons between each solution set within the same striatal subregion are shown above the line plot where the color depicts the corresponding comparison as indicated in the legend to the left of the plot.

**Figure 6 Supplement 2:**
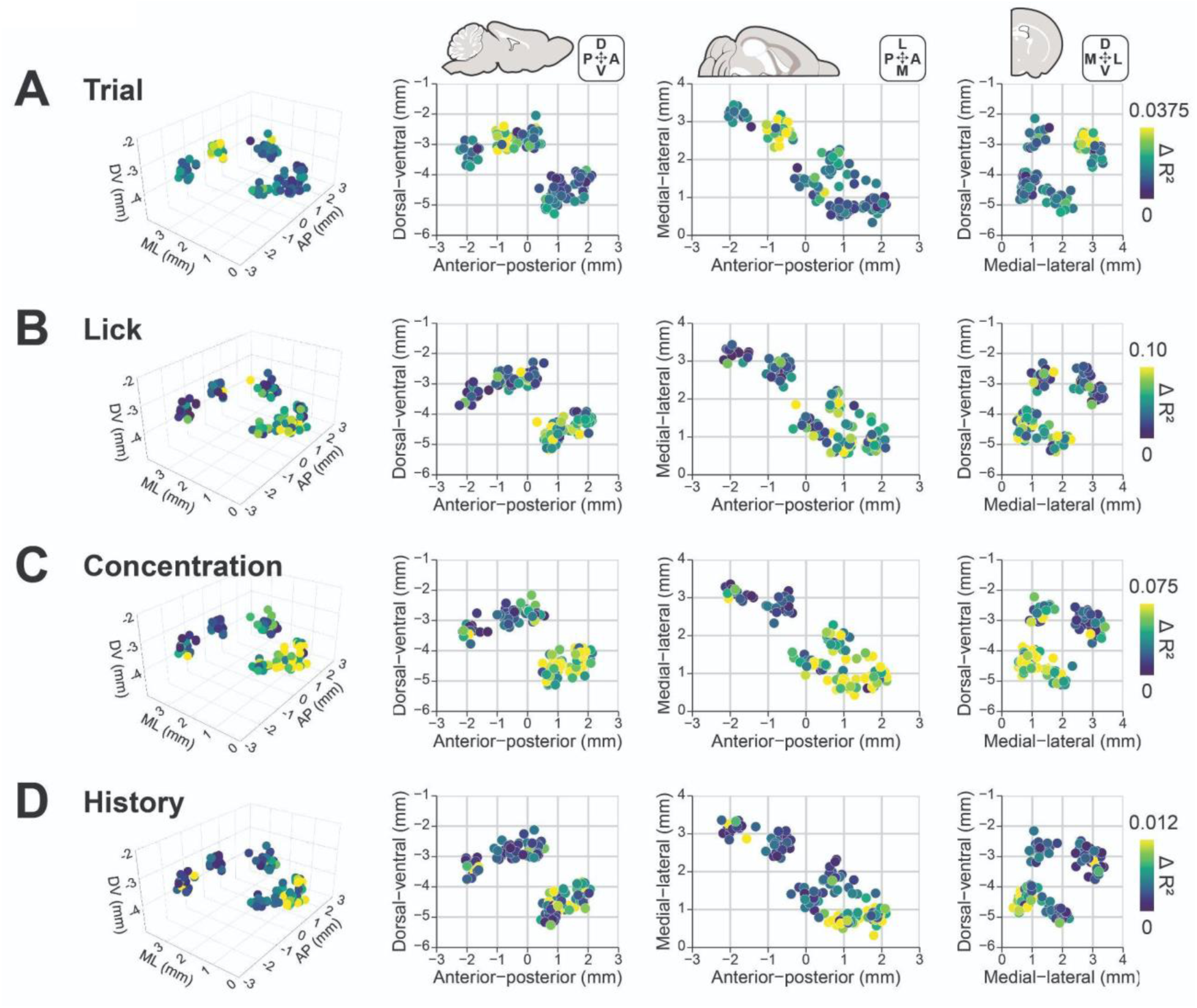
Histological analysis of striatal DA representations of consummatory behavior. Spatial distribution of the results of the general linear model analysis of striatal DA dynamics during consumption under the WR:NaCl restriction/solution configuration shown in Figure 6. (A-D) from left to right: representation of striatal fiber placement in 3d, sagittal, horizontal, and coronal orientations. Rows display each predictor as defined in Figure 6, and color depicts the change in GRAB-DA signal prediction (ΔR^2^) with removal of each predictor (each row has an independent color scale as defined on the right). Fiber positions have been jittered by 0.1mm to reduce overlap.

**Figure 6 Supplement 3:**
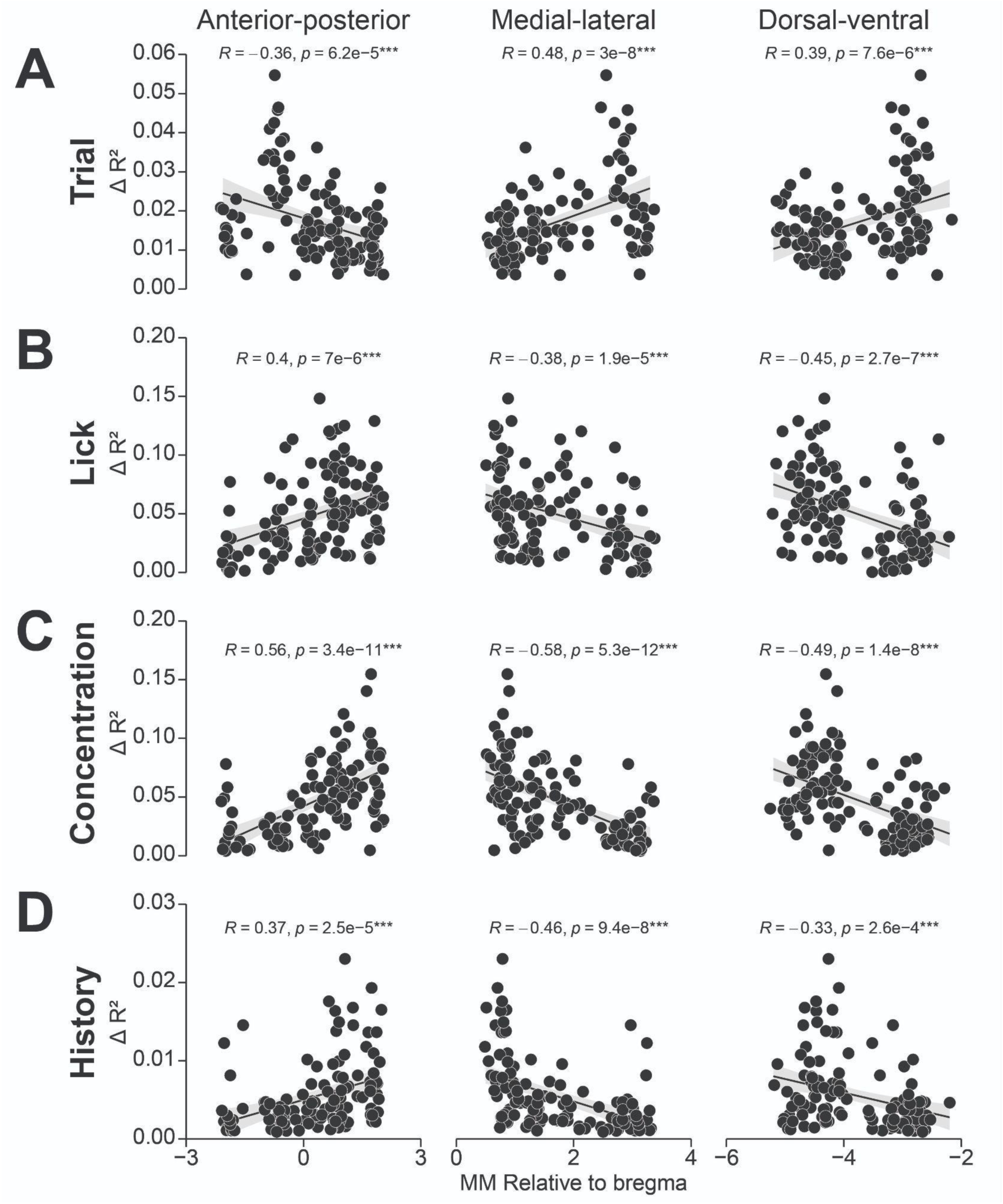
Correlation between atlas axes and DA representations of consummatory behavior. Correlation between the position in anterior-posterior (left), medial-lateral (middle), and dorsal-ventral (right) axes of the mouse brain against the change in GRAB-DA signal prediction (ΔR^2^) with removal of each predictor: trial (A), licking (B), solution concentration (C), and concentration history (D). Text depicts the results of a Pearson’s correlation test. Fiber positions have been jittered by 0.1mm to reduce overlap.

**Figure 6 Supplement 4:**
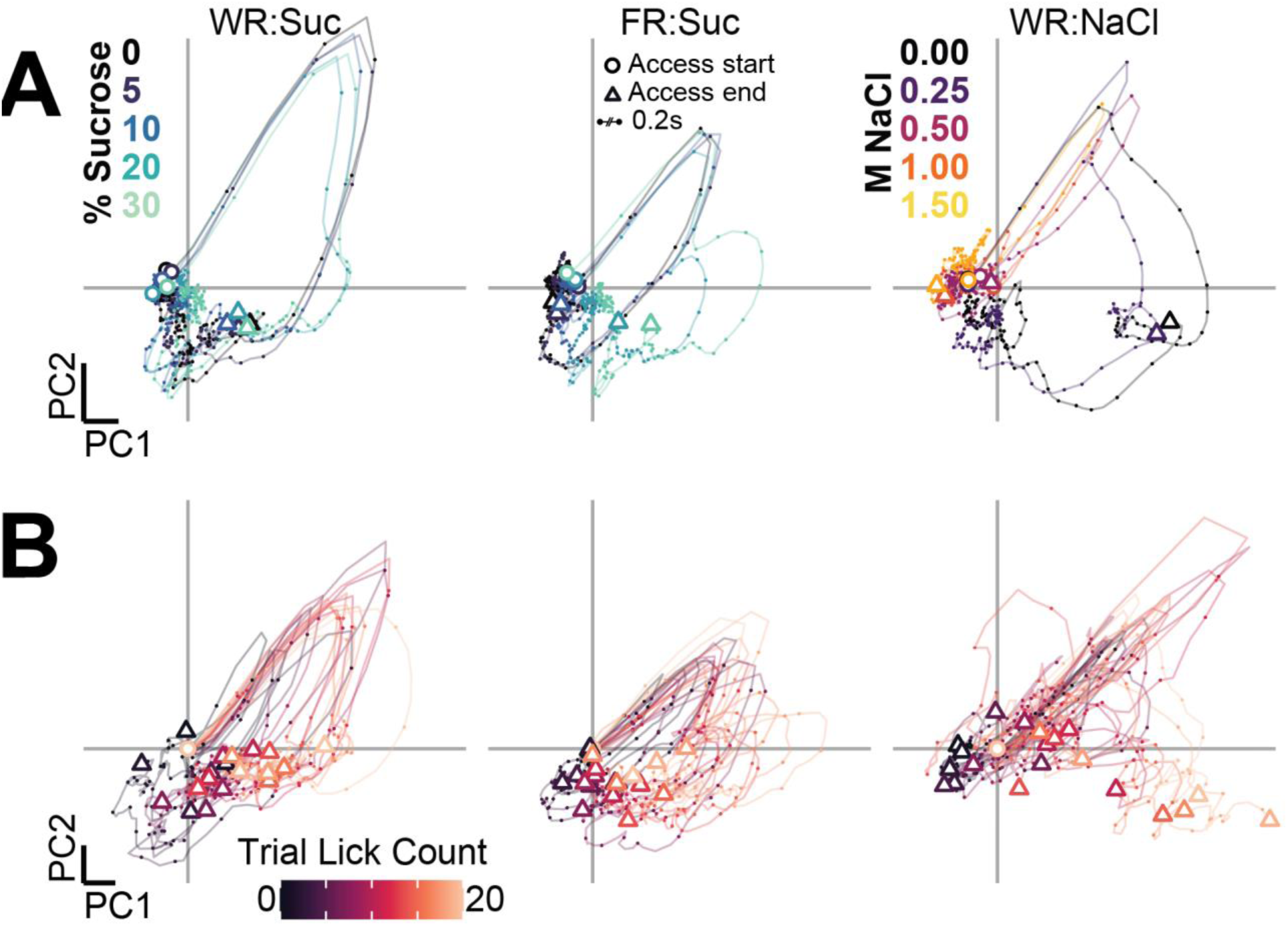
Striatal DA PCA trajectories during consumption for a representative subject. Mean PCA trajectories for a representative subject grouped by solution (A) and lick count bin (B) faceted horizontally by restriction/solution confirmation. Plotting conventions match Figure 6G. *Note: All of the shown trajectories come from the same PCA model that was built using data from all subjects*, ensuring that the principal components capture variance shared across the entire dataset.

**Figure 7 – Supplemental 1.**
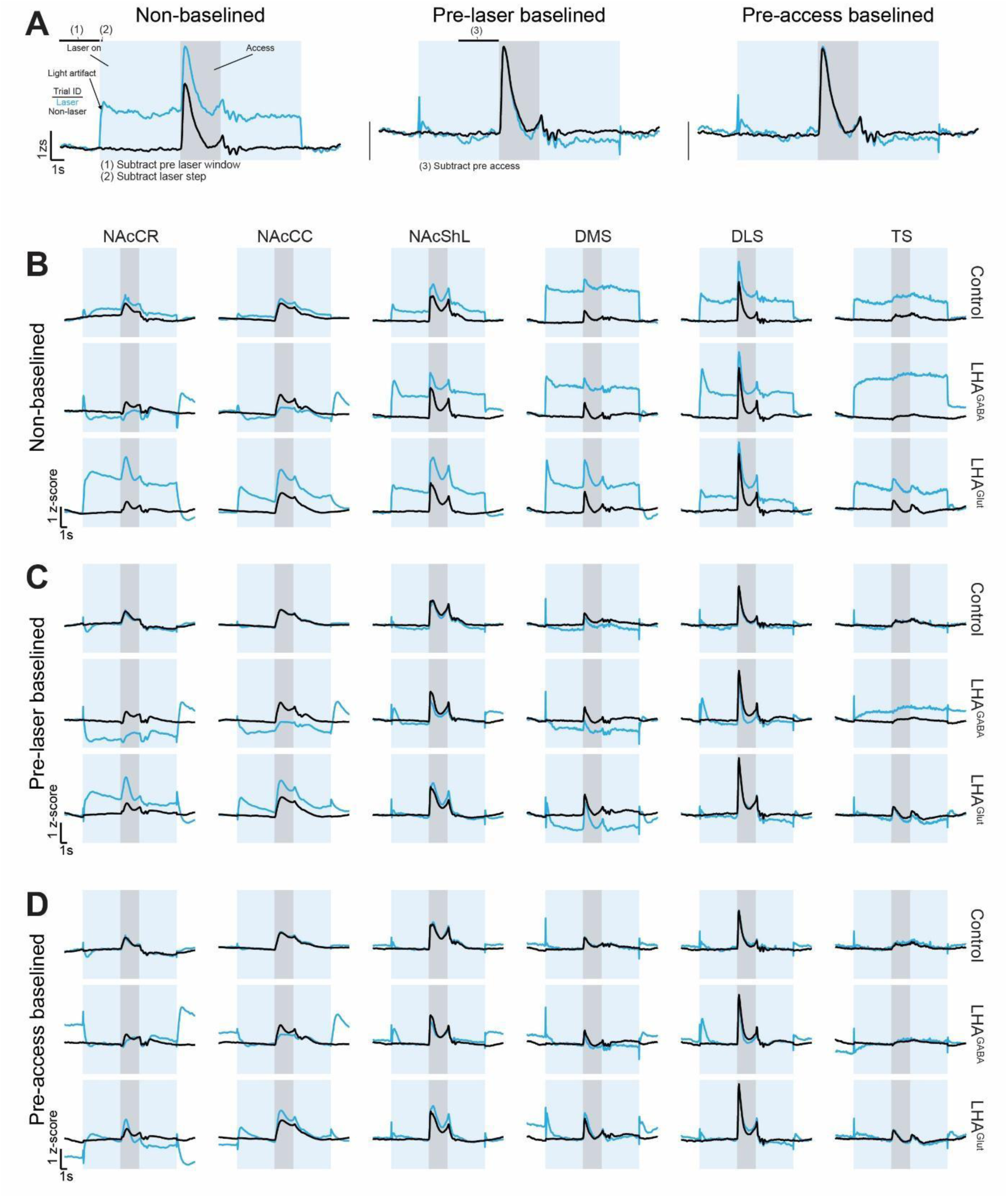
Pre-processing for simultaneous optogenetics and multi-site fiber photometry. (A) Pre-processing pipeline for artifact correction in a representative control mouse. For data shown in Figure 7E-H and Figure 7 **supplement 2A**, the signal is corrected by subtracting the mean signal during the 3s window prior to the laser onset time (1), then the light artifact is removed by subtracting the mean signal during the 0.2s window immediately following laser onset (2). For data shown in Figure 7 **supplement 2B** the signal is further corrected by subtracting out the 3s window prior to the access onset time. (B-D) Mean GRAB-DA signal at each stage of pre-processing stages across striatal regions (columns) and groups (rows): Non-baselined signals (B), following pre-laser baseline correction (C), and following pre-access baseline correction (D).

**Figure 7 – Supplemental 2.**
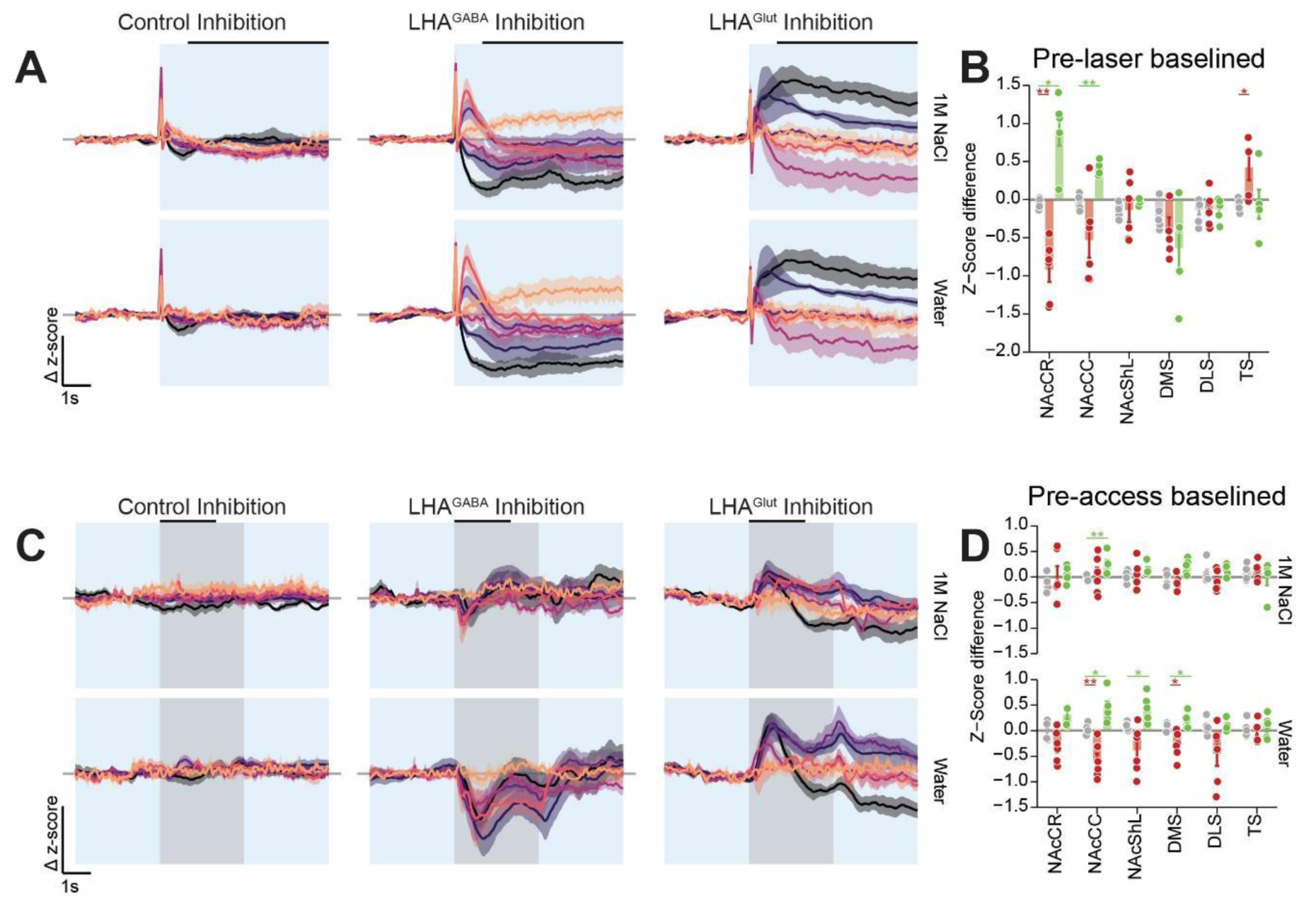
Additional analysis of LHA^GABA^ and LHA^Glut^ regulation of striatal dopamine during consumption. (A) Δzs following pre-laser baseline adjustment (see Figure 7 **supplement 1**) prior to the access period to laser emission in control (left), LHA^GABA^ HR (middle), and LHA^Glut^ HR (right) groups. The blue rectangle indicates the laser emission window, line color indicates striatal region, and horizontal black lines indicate the analysis window that means were calculated from. (B) Mean Δzs response during the analysis window. Asterisks indicate results from a Friedman rank sum test (Bonferroni adjusted for multiple comparisons to the same control). (C) Δzs following pre-access baseline adjustment (see Figure 7 **supplement 1**) prior to the access period to laser emission (same plotting convention as (A)). (D) Mean Δzs response during the analysis window (same plotting convention as (B))

## METHODS

### KEY RESOURCES TABLE

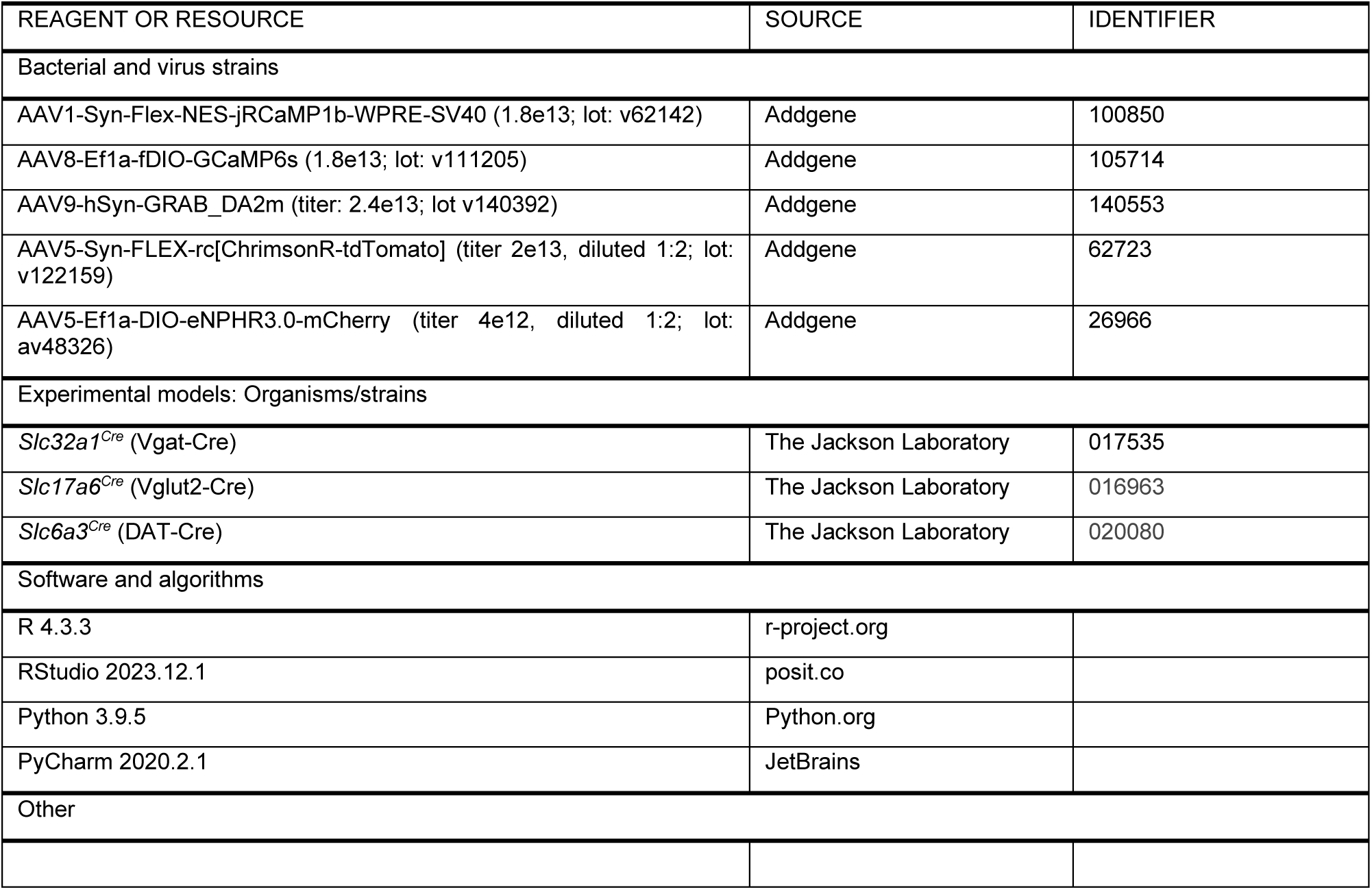

| REAGENT OR RESOURCE | SOURCE | IDENTIFIER |
| --- | --- | --- |
| Bacterial and virus strains |  |  |
| AAV1-Syn-Flex-NES-jRCaMP1b-WPRE-SV40 (1.8e13; lot: v62142) | Addgene | 100850 |
| AAV8-Ef1a-fDIO-GCaMP6s (1.8e13; lot: v111205) | Addgene | 105714 |
| AAV9-hSyn-GRAB_DA2m (titer: 2.4e13; lot v140392) | Addgene | 140553 |
| AAV5-Syn-FLEX-rc[ChrimsonR-tdTomato] (titer 2e13, diluted 1:2; lot: v122159) | Addgene | 62723 |
| AAV5-Ef1a-DIO-eNPHR3.0-mCherry (titer 4e12, diluted 1:2; lot: av48326) | Addgene | 26966 |
| Experimental models: Organisms/strains |  |  |
| *Slc32a1*<sup>Cre</sup> (Vgat-Cre) | The Jackson Laboratory | 017535 |
| *Slc17a6*<sup>Cre</sup> (Vglut2-Cre) | The Jackson Laboratory | 016963 |
| *Slc6a3*<sup>Cre</sup> (DAT-Cre) | The Jackson Laboratory | 020080 |
| Software and algorithms |  |  |
| R 4.3.3 | r-project.org |  |
| RStudio 2023.12.1 | posit.co |  |
| Python 3.9.5 | Python.org |  |
| PyCharm 2020.2.1 | JetBrains |  |
| Other |  |  |

### METHOD DETAILS

#### Animals

All studies were conducted in accordance with the Guide for the Care and Use of Laboratory Animals of the National Institute of Health and were pre-approved by the University of Washington Animal Care and Use Committee (IACUC). Transgenic mice on a C57BL/6J background including Vgat-cre^49^, Vglut2-cre^49^, and DAT-cre^100^ were obtained from Jackson Laboratory and breed with wild-type mice to produce heterozygous mutant animals. Surgery was conducted on mice at least P55 and all mice were singly housed to minimize damage to optical implants. Mice were kept on a reverse light/dark cycle and all experiments were performed during the dark cycle.

#### Surgery

As previously described^22^, mice were anesthetized with isoflurane (5% induction, 1-2% maintenance) and prepared for surgery by shaving the surgical site with an electric trimmer. Mice were placed in a stereotaxic frame (Kopf) outfitted with infrared heating pad to provide heat support and the surgical site with 3 alternating scrubs of betadine and 70% ethanol. Local anesthesia (2% lidocaine) was injected in the skin overlaying the skull and systemic analgesia was provided subcutaneously (5mg/kg carprofen). We prepared the skull using a scalpel to make an incision in the scalp, scrape away tissue overlaying the skull, and score the skull to enhance adhesive bonding. Three guide holes were drilled (2 laterally over the cerebellum, 1 over the olfactory bulb) and stainless-steel skull screws were inserted before the skull was leveled. For the injection of viral vectors, a burr hole was created using a drill, a glass injection pipette was lowered to the brain region of interest (**Table 1**), virus was infused at a rate of 1nL/s (Nanoject III), and the pipette was left in place for 2–5 minutes post-infusion before being retracted. For experiments involving LHA fiber-photometry, a single 400µm optic fiber with 2.5mm stainless steel ferrules was implanted. For experiments involving multi-site striatal recordings, optic fibers with 1.25mm zirconium ferrules (8 x 200µm (6 Striatum, 2 LHA) or 4 x 400µm (Striatum) + 2x 200µm (LHA)) were implanted sequentially. The process of fiber implantation consisted of drilling a burr hole, lowering the fiber to overlay the brain region of interest, and securing the ferrule to the skull using super glue, accelerated with the liquid component of dental cement (Ortho-Jet). Head-fixation components, consisting of stainless-steel head rings(Gordon-Fennell et al., 2023) or head plates (12 x 28 mm), were secured to the skull using super glue and dental cement. Mice were removed from the stereotaxic frame and allowed to fully recover on heat support prior to being single housed.

**Table 1.**
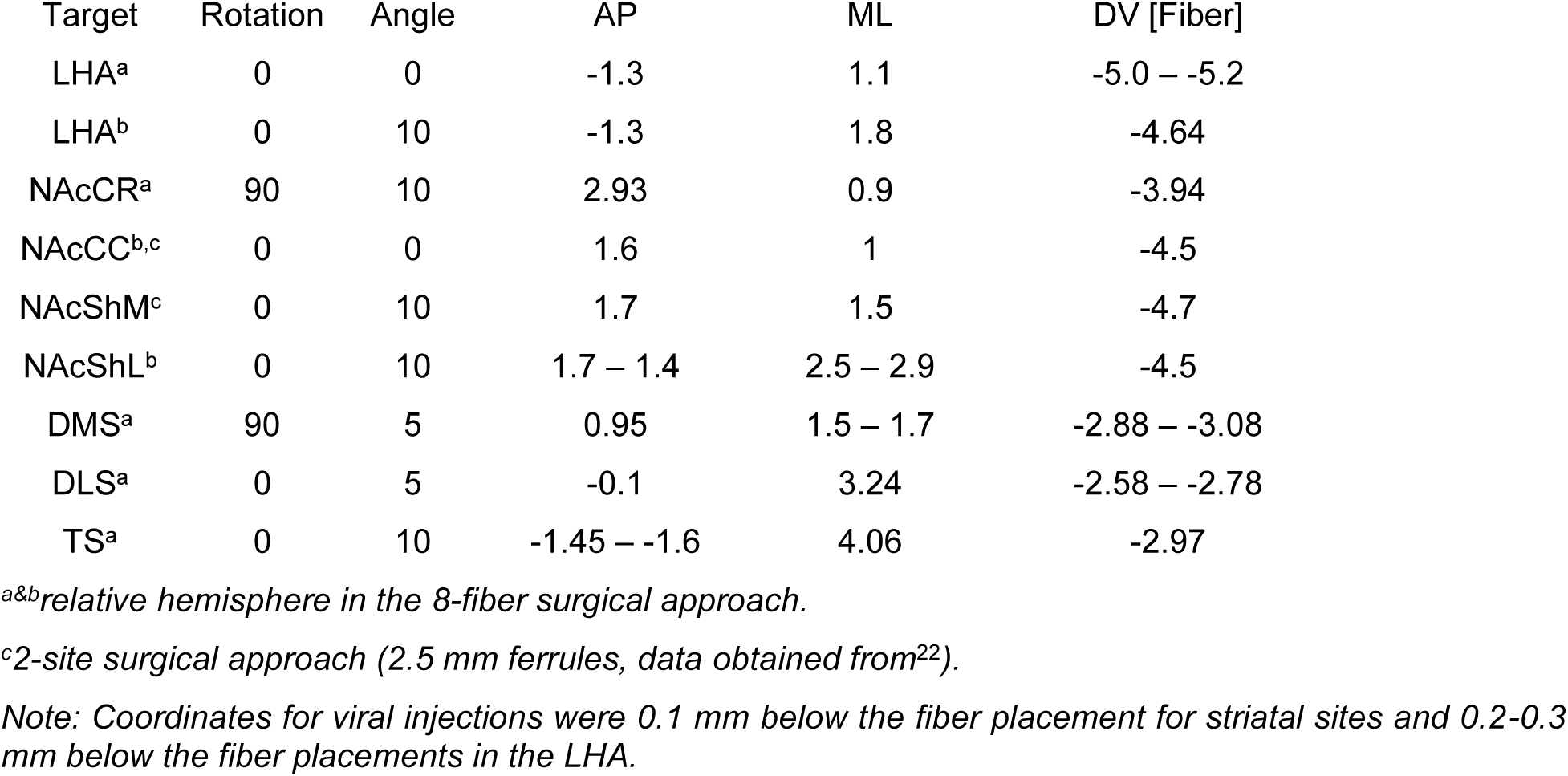
Stereotaxic Coordinates.

#### Brief-Access Taste Task

Mice were tested in the brief-access taste task using an Arduino-based OHRBETS system as previously described^22^. During behavioral sessions, mice were placed in a Falcon centrifuge tube (Millipore Sigma) head-fixed in a 3d printed stage. An Arduino Mega 2560 REV3 served as a microcontroller for controlling behavioral hardware including micro servos (Tower Pro SG92R) and five solenoids (Parker 003-0257-900). Licks were detected either via a capacitive touch sensor (Adafruit MPR121) or a custom voltage detector (see https://github.com/agordonfennell/OHRBETS). The 3D-printed multi-spout assembly used two servos—one driving a linear actuator to retract the spout and another rotating a radial five-spout head— allowing fixed access periods to any one of the five spouts. All task and behavioral events were timestamped and recorded via USB serial communication using a Python program.

Subjects were either food- or water-restricted to 90% of their baseline body weight, habituated to head-fixation for three sessions, and then trained over two sessions to consume 30% sucrose. Each multi-spout session began with a 2–3 min baseline period, followed by 100 trials of three-second access to one of 5 solutions (pseudorandom order with 2 presentations of each solution every 10 trials) separated by an 11– 16 second inter-trial interval (sampled from a uniform distribution). Each lick delivered approximately 1.5 μL; importantly, the identity of the solution on each trial was unknown to the mouse, requiring the mouse to lick and taste to identify the solution. All mice completed the multi-sucrose solution set under one restriction (food or water), transitioned to the alternate restriction state before repeating the multi-sucrose set. Subsequently, mice were water-restricted and performed the multi-NaCl solution set. Each switch in restriction was followed by a three-day acclimation period, and each new solution set or restriction state was followed by at least one training session before recording commenced.

#### Fiber Photometry

Fiber photometry was conducted using either a lock-in demodulated system RZ10X (Tucker Davis Technologies [TDT]) or an interleaved CMOS system, FP3002 (Neurophotometrics [NPM]).

The TDT system was outfitted with 3 LEDs with the following configuration: 465nm at 211Hz / 20-40µW for green indicators (GCaMP and GRAB-DA), 560nm at 449Hz / 40µW for red indicators (jRCaMP1b), and 405nm 331Hz / 20-40µW for isosbestic control. We used a 6-port minicube (Doric, Doric, FMC6_IE(400-410)_E1(460–490)_F1(500–540)_E2(555–570)_F2(580–680)_S) coupled to a 1 meter patch cable that was connected to the animal following head-fixation. Behavioral events were synchronized with fiber photometry signals using TTL inputs from the OHRBETS system to the TDT system.

The NPM system was outfitted with 3 LEDs: 470nm at 45 Hz / 30-70µW for green indicators (GRAB-DA) and 415nm at 45 Hz / 10-70µW for isosbestic control. We recorded up to 4 400µm fibers using a 4x branching patch cable (Doric, BBP(4)_400/440/900-0.37_3m_FCM-4xZF1.25_LAF) or up to 6 200µm fibers using a 12x branching patch cable (BBP(12)_200/220/900-0.37_3m_FCM-12xMF1.25_LAF). Behavioral events were synchronized with fiber photometry signals using serial input from the OHRBETS system to the computer connected to the NPM system.

With both systems, data was initially converted from both recording systems into a uniform data format. Data was smoothed using a 100ms moving mean and resampled to 20Hz. Photobleaching was corrected by fitting and subtracting a 3^rd^ or 4^th^ degree polynomial to each signal individually. Because head-fixation largely eliminates motion artifacts typical to fiber-photometry^7,22^, we used the isosbestic channel to confirm that motion artifacts were absent but did not use it to correct the green or red channels.

#### Optogenetic Characterization

All optogenetic characterization experiments were performed under head-fixed conditions, with mice habituated for at least three days prior to testing. Multi-site fiber photometry was conducted over three sessions, during which fiber branch and recording regions were counterbalanced across both sessions and subjects. To assess the effect of LHA stimulation on striatal DA signaling (**Figure 2A–E**), we used the NPM system to record GRAB-DA signals from 4–6 striatal sites while delivering pulsed laser illumination (3 mW, 5 ms pulses, 3 s train, 2–20 Hz, 10–15 trials per site/frequency) bilaterally to the LHA in mice expressing Chrimson in LHA^GABA^ or LHA^Glut^ neurons or expressing mCherry as a fluorescent protein control, across 1– 3 sessions. To assess the effect of LHA inhibition on striatal DA (**Figure 2F–J**), we employed a TDT system to record GRAB-DA signals from 2 striatal sites while delivering continuous laser illumination (620 nm, 10 mW, 10 s pulse, 20–40 s inter-trial interval, 5 trials per site) bilaterally to the LHA in mice expressing halorhodopsin in LHA^GABA^ or LHA^Glut^ neurons or expressing mCherry as a fluorescent protein control, across 3 sessions. The TDT system was used to minimize artifacts from continuous red laser illumination. Finally, to assess the effect of presynaptic terminal stimulation (**Figure 3**), we again used the NPM system to record GRAB-DA signals from four striatal sites while delivering laser illumination (635 nm, 0.25 mW, 5 ms pulses, 3 s train, 1–20 Hz, 10 trials per site/frequency) directly to one striatal site in mice expressing Chrimson in midbrain DA neurons, across four sessions.

#### Optogenetic Brief-Access Taste Task

Mice were first exposed to the standard optogenetic inhibition protocol and brief-access taste task, as outlined above, and then proceeded to the optogenetic brief-access taste task (**Figure 7**). Animals were water-restricted and completed four sessions consisting of 100 trials each, with three-second access to either water or 1M NaCl (presented in equal proportions). Continuous laser illumination (620 nm, 10 mW) began six seconds prior to access onset and persisted until six seconds following access offset (15 seconds total) and there was an 11-16 second inter-trial interval between laser offset and the following laser onset. This design enabled us to evaluate the effect of laser illumination alone on the fiber photometry signal and perform appropriate re-baselining procedures (**Figure 7 supplement 2**).

#### Histology

The location of optic fibers and viral expression were determined following the conclusion of each experiment as previously described^22^. Briefly, mice were transcardially perfused with PBS and 4% PFA before being decapitated and storing skulls with cranial implants intact for 24-72 hours in 4% PFA (72 hours was preferred for 200µm optic fibers). Brains were removed from the skull, post fixed in 4% PFA for 24 hours, and then stored in 30% sucrose in PBS on a shaker for 48 hours. Brains were serially sectioned at 40 µm on a cryostat (Leica) and tissue was stored in PBS with sodium azide before being mounted on slides. Brain sections that included fiber tips were imaged using epifluorescence (Zeiss, ApoTome2) or confocal (Olympus, FV3000) microscopes. The location of fiber tips and viral expression were determined using manual alignment with an atlas^101^. Data was excluded from analysis if the fiber tip could not be located or was located outside of the target brain region. Fiber positions were visualized in 3d using Plotly in R and in 2D using a combination of R and Adobe Illustrator.

### QUANTIFICATION AND STATISTICAL ANALYSIS

#### General statistical methods

All scripts for analysis can be found in our git repository (*link will be included with publication*). All statistical analyses were performed in R, except those for the GLM were performed in Python. All ANOVAs were computed using the package “afex” (version 1.3-1); HSD post-hoc tests were computed using “emmeans” (version 1.10.0); Pearson correlations, Student’s t-test, Wilcox rank sum test, Friedman rank sum test, and Bonferroni post-hoc corrections were all computed using base R. Statistical test information and results from main effects are outlined in figure legends, the results from every post-hoc test are listed in the supplemental statistical table (**Table S1**). Sample sizes are displayed in the figure and the identity of samples are listed in the figure legend. Unless otherwise specified in the figure legend, the center represents the mean and the dispersion represents the SEM. Significance was defined as p<0.05 for all statistical tests. Subjects were pseudo-randomly assigned to experimental or control viral groups to balance sex and age across groups.

#### Exclusions

For all experiments involving optogenetic regulation of the LHA, mice were excluded if fibers intended for the LHA were misplaced. For all photometry experiments, individual recording sites were excluded if the fiber was determined to be outside of the intended region. Mice with a single fiber misplacement were excluded from the analysis of cross-regulation of DA release (**Figure 3**) and correlated dopamine dynamics (e.g. **Figure 4I-K**). During branching fiber-photometry recording, individual subregions within individual sessions were excluded if there were over 5% of samples clipped, which rarely occurred with poor fiber connection and/or low transmission branches paired with low brightness sites.

#### Analysis of perievent time-histogram fiber photometry

We generated perievent time-histograms relative to the onset of each access period. Fiber-photometry signals were first z-scored using the 3s pre-event window for the mean and standard deviation across all trials within a session and were then baseline corrected by subtracting the mean signal in the 3s pre-event window on each trial unless otherwise noted. The LHA^Ratio^ was computed by adding a small constant offset to the LHA^GABA^ and LHA^Glut^ z-scores to avoid division by zero and sign reversals from negative denominators. The offset LHA^GABA^ and LHA^Glut^ signals were then divided, and the resulting ratio was subsequently z-scored and baseline corrected. For analysis of fiber photometry signals during the brief-access taste task, data was filtered to trials with licking.

For statistical analysis, mean signals during different periods of the access window were calculated for each combination of subject, signal ID (e.g. region, or LHA subpopulation), and solution. The absolute range in fiber photometry z-score (**Figure 1, Figure 5**) was calculated by taking the absolute difference between the fiber photometry response during trials with the highest and lowest concentrations. The absolute range in consumption was calculated by taking the absolute difference between the mean number of licks during trials with the highest and lowest concentrations.

The effect of LHA inhibition on striatal DA signaling during consumption were assessed in the optogenetic brief-access taste task using a Δ z-score (**Figure 7 A-H, Figure 7-supplement 1**). First, perievent time-histograms were computed as described above using the 3s pre-laser period for computing the z-score. The pre-laser baselined signal was calculated by removing the laser artifact by subtracting the mean signal during the first 0.2s following laser onset. The pre-access baselined signal was calculated by subtracting the mean z-score during the 3s prior to access onset. The Δ z-score for these two baselined signals were then calculated by taking the difference between the mean z-score during trials with and without laser illumination.

#### Analysis of spontaneous DA dynamics

Spontaneous GRAB-DA dynamics were assessed in 119 striatal sites across 22 mice during 2-minute baseline periods while mice were head-fixed prior to optogenetic regulation or brief-access taste task sessions (mean session count per subject: 19; SD: 1.56). Spontaneous transients were determined by first z-scoring the baseline of each session using the mean and SD from the entire 2-minute baseline. We then calculated the moving z-score with a 3s preceding window and then threshold searching using the “findpeaks” function from the “ggpmisc” library (version 2.4.4) with minimum peak amplitude of 1.5 z-score and minimum inter-peak interval of 3s, and we then found the corresponding peaks in the z-scored baseline time-series. Perievent time histograms were generated centered on each peak which were used to estimate tau (**Figure 4F**) and peak amplitude (**Figure 4G**). Autocorrelations were calculated for each session individually before averaging for each subject x striatal site (**Figure 4E**). Correlations were analyzed in a subset of subjects that had all 6 sites confirmed to be in the correct location, all sessions were concatenated before calculating the pairwise correlation of GRAB-DA activity at every site (**Figure 4I-K**). For each site, the mean Pearson correlation (Mean R) was calculated by averaging R for all pairwise correlations with all other striatal sites (**Figure 4J**).

#### Generalized linear models

To quantify how task variables and behavior contribute to photometry signals during consumption, we constructed a separate GLM for each fiber in each solution/restriction condition (**Figure 6A**). For each trial, we extracted a peri-event window spanning 3 seconds before to 3 seconds after solution access, sampled at 20 Hz (180 time points per trial). The dependent variable (*Y*) was the z-scored, baseline-subtracted GRAB-DA photometry signal, and therefore has length 180n, where n is the number of trials with licking. The predictor matrix (*X*) has dimension 180n x 551, and is composed of four sets of predictors. The first three predictor sets represent “time within trial,” “solution concentration,” and “solution history" consist of 180 predictors (one per timepoint in trial). The fourth predictor set has 11 coefficients. In greater detail, the 4 predictor sets are defined as follows:

1. 180 predictors representing the “time within trial”. Within each trial, these are represented as a 180x180 identity matrix, independent of the solution and behavior. We included these 180 predictors to allow the effect of uniform sensory information (e.g. spout extension, presence of the spout, etc.) to vary across the timepoints within each trial.
2. 180 predictors representing the “solution concentration”. Within each trial, these are represented as a multiple of the 180x180 identity matrix, with diagonal elements equal to the scaled solution concentration, ranging from 0 (lowest solution value) to 1 (highest solution value). We included these 180 predictors to allow for the effect of “solution concentration” to vary across the timepoints within each trial.
3. 180 predictors representing the “solution history”. Within each trial, these are represented as a multiple of the 180x180 identity matrix, with diagonal elements equal to the mean of the scaled solution concentration over the last three trials. We included these 180 predictors to allow for the effect of “solution history” to vary across the timepoints within each trial.
4. 11 predictors representing the “licking” behavior. Within each trial, these are represented as a 180x11 matrix constructed by convolving a binary lick time series with a half-normal kernel (σ = 2.5 s), and then expanding the predictor to include five time-lagged versions in both time directions (-0.25s, -0.2s, -0.15s, -0.1s, -0.05s, 0.05s, 0.1s, 0.15s, 0.2s, 0.25s). We included these five time-lagged versions to capture neural activity that may precede or follow the licking behavior.

To visualize model performance for a representative fiber, we conducted 10-fold cross-validation: in each fold, we fit the model to 90% of the trials and tested on the held-out 10%, ensuring that all trials were used in exactly one test fold. We then plotted the model predictions against the observed GRAB-DA photometry signal (**Figure 6B**).

Next, to assess the contribution of each predictor to the variance in the photometry signals, we computed the difference in proportion of variance explained (ΔR²) between a model containing all the predictors, and a model that omits one of the four predictor sets. We then compared the ΔR² values from all fibers across striatal subregions using ANOVA followed by an HSD post-hoc test (**Figure 6C-F, Figure 6 supplement 1**).

#### Principal component analysis (PCA)

PCA on GRAB-DA signals across striatal subregions was performed on the subset of subjects for whom all six recording sites were verified to be in the correct location (n=13). Data was filtered to include only trials with licking. Because we recorded from the same six striatal subregions in all subjects, we were able to concatenate the peri-event time histograms across all trials from the thirteen subjects before applying PCA.

The matrix used for PCA contained six columns (one column per striatal subregion) and 3,161,070 rows (derived from 11,330 trials × 279 timepoints per trial, spanning −2.95 s to +10.95 s relative to access onset). On average, each of thirteen subjects contributed 872 trials (SD: 134).

For each of the first two principal components, our analysis proceeded as follows. First, we centered the PC scores within each trial by subtracting out the score at the onset of the access period. Then, we computed the mean score for each timepoint across all of the trials corresponding to a given grouping of interest (e.g., 0% sucrose under WR:NaCl [**Figure 6G**], or 5-6 licks per trial under WR:NaCl [**Figure 6J**]). That is, for each grouping, we identified all of the trials corresponding to this grouping, and averaged the score values across those trials for each of the 279 timepoints in order to obtain a vector of length 279 for that grouping; in what follows, we refer to these vectors of length 279 as “averaged PCA trajectories”.

For groupings defined by solution (e.g. 0% sucrose under WR:NaCl), we display the resulting averaged PCA trajectories in **Figure 6G**, for the 0-10.95s time window. For groupings defined by lick rate (e.g. 5-6 licks per trial), we display the resulting averaged PCA trajectories in **Figure 6J**, for the 0-3s window. Note that for groupings defined by lick rate, the lick rate was binned into bins of 2 licks per trial (i.e. 1-2 licks per trial were in one bin, 3-4 licks per trial were in another bin, etc.).

Next, we restricted analysis to the 10 (of 13) mice for which no restriction/solution configurations have missing lick bins (e.g. if a mouse has no trials with 5-6 licks during WR:NaCl, then it was omitted). We then quantified trajectory similarity in PC space across different solution/restriction configurations. That is, for each subject, for each time point in the trial, and for each binned lick trajectory, we computed the Euclidean distance of the PC trajectory points between the different restriction/solution configurations (as there are three restriction/solution configurations, we computed three pairwise Euclidean distances). This resulted in 10 vectors (10 bins of lick counts) of length 279 (279 time points in the trial) for each of the 10 subjects in each of the three pairs of restriction/solution configurations. For the sake of visualizing the similarity in trajectories over the access window, we averaged these curves across the 10 bins of licks as well as across the 10 subjects, resulting in a vector of length 279 for each of the pairs of restriction/solution configurations. (**Figure 6K**).

## REFERENCES

1. Zhu, Z., Gong, R., Rodriguez, V., Quach, K.T., Chen, X., and Sternson, S.M. (2025). Hedonic eating is controlled by dopamine neurons that oppose GLP-1R satiety. Science 387, eadt0773. 10.1126/science.adt0773.

2. Palmiter, R.D. (2007). Is dopamine a physiologically relevant mediator of feeding behavior? Trends in neurosciences 30, 375–381. 10.1016/j.tins.2007.06.004.

3. Wise, R.A. (2004). Dopamine, learning and motivation. Nature Reviews Neuroscience 5, 483–494. 10.1038/nrn1406.

4. Salamone, J.D., and Correa, M. (2012). The mysterious motivational functions of mesolimbic dopamine. Neuron 76, 470–485. 10.1016/j.neuron.2012.10.021.

5. Berke, J.D. (2018). What does dopamine mean? Nat. Neurosci. 21, 787–793. 10.1038/s41593-018-0152-y.

6. Schultz, W. (1998). Predictive reward signal of dopamine neurons. Journal of neurophysiology 80, 1– 27.

7. Menegas, W., Babayan, B.M., Uchida, N., and Watabe-Uchida, M. (2017). Opposite initialization to novel cues in dopamine signaling in ventral and posterior striatum in mice. Elife 6, e21886. 10.7554/elife.21886.

8. Cohen, J.Y., Haesler, S., Vong, L., Lowell, B.B., and Uchida, N. (2012). Neuron-type-specific signals for reward and punishment in the ventral tegmental area. Nature 482, 85–88. 10.1038/nature10754.

9. Stuber, G.D., and Wise, R.A. (2016). Lateral hypothalamic circuits for feeding and reward. Nature Neuroscience 19, 198–205. 10.1038/nn.4220.

10. Castro, D.C., Cole, S.L., and Berridge, K.C. (2015). Lateral hypothalamus, nucleus accumbens, and ventral pallidum roles in eating and hunger: interactions between homeostatic and reward circuitry. Frontiers in Systems Neuroscience 9, 90. 10.3389/fnsys.2015.00090.

11. Bernardis, L.L., and Bellinger, L.L. (1996). The lateral hypothalamic area revisited: Ingestive behavior. Neurosci. Biobehav. Rev. 20, 189–287. 10.1016/0149-7634(95)00015-1.

12. Nieh, E.H., Matthews, G.A., Allsop, S.A., Presbrey, K.N., Leppla, C.A., Wichmann, R., Neve, R., Wildes, C.P., and Tye, K.M. (2015). Decoding neural circuits that control compulsive sucrose seeking. Cell 160, 528–541. 10.1016/j.cell.2015.01.003.

13. Faget, L., Osakada, F., Duan, J., Ressler, R., Johnson, A.B., Proudfoot, J.A., Yoo, J., Callaway, E.M., and Hnasko, T.S. (2016). Afferent Inputs to Neurotransmitter-Defined Cell Types in the Ventral Tegmental Area. Cell Reports 15, 2796–2808. 10.1016/j.celrep.2016.05.057.

14. Watabe-Uchida, M., Zhu, L., Ogawa, S.K., Vamanrao, A., and Uchida, N. (2012). Whole-brain mapping of direct inputs to midbrain dopamine neurons. Neuron 74, 858–873. 10.1016/j.neuron.2012.03.017.

15. Beier, K.T., Steinberg, E.E., DeLoach, K.E., Xie, S., Miyamichi, K., Schwarz, L., Gao, X.J., Kremer, E.J., Malenka, R.C., and Luo, L. (2015). Circuit Architecture of VTA Dopamine Neurons Revealed by Systematic Input-Output Mapping. Cell 162, 622–634. 10.1016/j.cell.2015.07.015.

16. Mickelsen, L.E., Bolisetty, M., Chimileski, B.R., Fujita, A., Beltrami, E.J., Costanzo, J.T., Naparstek, J.R., Robson, P., and Jackson, A.C. (2019). Single-cell transcriptomic analysis of the lateral hypothalamic area reveals molecularly distinct populations of inhibitory and excitatory neurons. Nat. Neurosci. 22, 642– 656. 10.1038/s41593-019-0349-8.

17. Bonnavion, P., Mickelsen, L.E., Fujita, A., Lecea, L. de, and Jackson, A.C. (2016). Hubs and spokes of the lateral hypothalamus: cell types, circuits and behaviour. J Physiology 594, 6443–6462. 10.1113/jp271946.

18. Rossi, M.A., Basiri, M.L., McHenry, J.A., Kosyk, O., Otis, J.M., Munkhof, H.E. van den, Bryois, J., Hübel, C., Breen, G., Guo, W., et al. (2019). Obesity remodels activity and transcriptional state of a lateral hypothalamic brake on feeding. Science 364, 1271–1274. 10.1126/science.aax1184.

19. Wang, Y., Eddison, M., Fleishman, G., Weigert, M., Xu, S., Wang, T., Rokicki, K., Goina, C., Henry, F.E., Lemire, A.L., et al. (2021). EASI-FISH for thick tissue defines lateral hypothalamus spatio-molecular organization. Cell 184, 6361–6377.e24. 10.1016/j.cell.2021.11.024.

20. Jennings, J.H., Ung, R.L., Resendez, S.L., Stamatakis, A.M., Taylor, J.G., Huang, J., Veleta, K., Kantak, P.A., Aita, M., Shilling-Scrivo, K., et al. (2015). Visualizing Hypothalamic Network Dynamics for Appetitive and Consummatory Behaviors. Cell 160, 516–527. 10.1016/j.cell.2014.12.026.

21. Nieh, E.H., Weele, C.M., Matthews, G.A., Presbrey, K.N., Wichmann, R., Leppla, C.A., Izadmehr, E.M., and Tye, K.M. (2016). Inhibitory Input from the Lateral Hypothalamus to the Ventral Tegmental Area Disinhibits Dopamine Neurons and Promotes Behavioral Activation. Neuron 90, 1286–1298. 10.1016/j.neuron.2016.04.035.

22. Gordon-Fennell, A., Barbakh, J.M., Utley, M.T., Singh, S., Bazzino, P., Gowrishankar, R., Bruchas, M.R., Roitman, M.F., and Stuber, G.D. (2023). An open-source platform for head-fixed operant and consummatory behavior. eLife 12, e86183. 10.7554/elife.86183.

23. Garcia, A., Coss, A., Luis-Islas, J., Puron-Sierra, L., Luna, M., Villavicencio, M., and Gutierrez, R. (2021). Lateral Hypothalamic GABAergic Neurons Encode and Potentiate Sucrose’s Palatability. Front Neurosci-switz 14, 608047. 10.3389/fnins.2020.608047.

24. Stamatakis, A.M., Swieten, M., Basiri, M.L., Blair, G.A., Kantak, P., and Stuber, G.D. (2016). Lateral Hypothalamic Area Glutamatergic Neurons and Their Projections to the Lateral Habenula Regulate Feeding and Reward. The Journal of Neuroscience 36, 302–311. 10.1523/jneurosci.1202-15.2016.

25. Barbano, M.F., Zhang, S., Chen, E., Espinoza, O., Mohammad, U., Alvarez-Bagnarol, Y., Liu, B., Hahn, S., and Morales, M. (2024). Lateral hypothalamic glutamatergic inputs to VTA glutamatergic neurons mediate prioritization of innate defensive behavior over feeding. Nat. Commun. 15, 403. 10.1038/s41467-023-44633-w.

26. Rossi, M.A., Basiri, M.L., Liu, Y., Hashikawa, Y., Hashikawa, K., Fenno, L.E., Kim, Y.S., Ramakrishnan, C., Deisseroth, K., and Stuber, G.D. (2021). Transcriptional and functional divergence in lateral hypothalamic glutamate neurons projecting to the lateral habenula and ventral tegmental area. Neuron 109, 3823–3837.e6. 10.1016/j.neuron.2021.09.020.

27. Stamatakis, A.M., and Stuber, G.D. (2012). Activation of lateral habenula inputs to the ventral midbrain promotes behavioral avoidance. Nature Neuroscience 15, 1105–1107. 10.1038/nn.3145.

28. Huang, H., Liu, X., Wang, L., and Wang, F. (2024). Whole-brain connections of glutamatergic neurons in the mouse lateral habenula in both sexes. Biol. Sex Differ. 15, 37. 10.1186/s13293-024-00611-5.

29. Ma, W., Li, L., Kong, L., Zhang, H., Yuan, P., Huang, Z., and Wang, Y. (2023). Whole-brain monosynaptic inputs to lateral periaqueductal gray glutamatergic neurons in mice. CNS Neurosci. Ther. 29, 4147–4159. 10.1111/cns.14338.

30. Li, Y., Zeng, J., Zhang, J., Yue, C., Zhong, W., Liu, Z., Feng, Q., and Luo, M. (2018). Hypothalamic Circuits for Predation and Evasion. Neuron 97, 911–924.e5. 10.1016/j.neuron.2018.01.005.

31. Ogawa, S.K., Cohen, J.Y., Hwang, D., Uchida, N., and Watabe-Uchida, M. (2014). Organization of monosynaptic inputs to the serotonin and dopamine neuromodulatory systems. Cell reports 8, 1105– 1118. 10.1016/j.celrep.2014.06.042.

32. Pollak Dorocic, I., Fürth, D., Xuan, Y., Johansson, Y., Pozzi, L., Silberberg, G., Carlén, M., and Meletis, K. (2014). A Whole-Brain Atlas of Inputs to Serotonergic Neurons of the Dorsal and Median Raphe Nuclei. Neuron 83, 663–678. 10.1016/j.neuron.2014.07.002.

33. Hahn, J.D., Gao, L., Boesen, T., Gou, L., Hintiryan, H., and Dong, H. (2022). Macroscale connections of the mouse lateral preoptic area and anterior lateral hypothalamic area. J Comp Neurol. 10.1002/cne.25331.

34. Menegas, W., Bergan, J.F., Ogawa, S.K., Isogai, Y., Venkataraju, K.U., Osten, P., Uchida, N., and Watabe-Uchida, M. (2015). Dopamine neurons projecting to the posterior striatum form an anatomically distinct subclass. eLife 4, e10032. 10.7554/elife.10032.

35. Menegas, W., Akiti, K., Amo, R., Uchida, N., and Watabe-Uchida, M. (2018). Dopamine neurons projecting to the posterior striatum reinforce avoidance of threatening stimuli. Nat Neurosci, 1–10. 10.1038/s41593-018-0222-1.

36. Verharen, J.P., Zhu, Y., and Lammel, S. (2020). Aversion hot spots in the dopamine system. Curr. Opin. Neurobiol. 64, 46–52. 10.1016/j.conb.2020.02.002.

37. Hajnal, A., Smith, G.P., and Norgren, R. (2004). Oral sucrose stimulation increases accumbens dopamine in the rat. Am J Physiology-regulatory Integr Comp Physiology 286, R31–R37. 10.1152/ajpregu.00282.2003.

38. Eshel, N., Bukwich, M., Rao, V., Hemmelder, V., Tian, J., and Uchida, N. (2015). Arithmetic and local circuitry underlying dopamine prediction errors. Nature 525, 243–246. 10.1038/nature14855.

39. Tobler, P.N., Fiorillo, C.D., and Schultz, W. (2005). Adaptive Coding of Reward Value by Dopamine Neurons. Science 307, 1642–1645. 10.1126/science.1105370.

40. Yuan, L., Dou, Y.-N., and Sun, Y.-G. (2019). Topography of Reward and Aversion Encoding in the Mesolimbic Dopaminergic System. J. Neurosci. 39, 6472–6481. 10.1523/jneurosci.0271-19.2019.

41. Brown, H.D., McCutcheon, J.E., Cone, J.J., Ragozzino, M.E., and Roitman, M.F. (2011). Primary food reward and reward-predictive stimuli evoke different patterns of phasic dopamine signaling throughout the striatum. The European journal of neuroscience 34, 1997–2006. 10.1111/j.1460-9568.2011.07914.x.

42. Smith, J.C. (2001). The history of the “Davis Rig.” Appetite 36, 93–98. 10.1006/appe.2000.0372.

43. Chen, Z., Chen, G., Zhong, J., Jiang, S., Lai, S., Xu, H., Deng, X., Li, F., Lu, S., Zhou, K., et al. (2022). A circuit from lateral septum neurotensin neurons to tuberal nucleus controls hedonic feeding. Mol Psychiatr, 1–18. 10.1038/s41380-022-01742-0.

44. Tesmer, A.L., Viskaitis, P., Donegan, D., Bracey, E.F., Grujic, N., Patriarchi, T., Peleg-Raibstein, D., and Burdakov, D. (2024). Orexin population activity precisely reflects net body movement across behavioral and metabolic states. bioRxiv, 2024.08.13.607750. 10.1101/2024.08.13.607750.

45. Farrell, K., Lak, A., and Saleem, A.B. (2022). Midbrain dopamine neurons signal phasic and ramping reward prediction error during goal-directed navigation. Cell Rep. 41, 111470. 10.1016/j.celrep.2022.111470.

46. Sharpe, M.J., Marchant, N.J., Whitaker, L.R., Richie, C.T., Zhang, Y.J., Campbell, E.J., Koivula, P.P., Necarsulmer, J.C., Mejias-Aponte, C., Morales, M., et al. (2017). Lateral Hypothalamic GABAergic Neurons Encode Reward Predictions that Are Relayed to the Ventral Tegmental Area to Regulate Learning. Current Biology. 10.1016/j.cub.2017.06.024.

47. Sharpe, M.J., Batchelor, H.M., Mueller, L.E., Gardner, M.P.H., and Schoenbaum, G. (2021). Past experience shapes the neural circuits recruited for future learning. Nat Neurosci 24, 391–400. 10.1038/s41593-020-00791-4.

48. Sharpe, M.J. (2024). The cognitive (lateral) hypothalamus. Trends Cogn. Sci. 28, 18–29. 10.1016/j.tics.2023.08.019.

49. Vong, L., Ye, C., Yang, Z., Choi, B., Chua, S., and Lowell, B.B. (2011). Leptin Action on GABAergic Neurons Prevents Obesity and Reduces Inhibitory Tone to POMC Neurons. Neuron 71, 142–154. 10.1016/j.neuron.2011.05.028.

50. Dana, H., Mohar, B., Sun, Y., Narayan, S., Gordus, A., Hasseman, J.P., Tsegaye, G., Holt, G.T., Hu, A., Walpita, D., et al. (2016). Sensitive red protein calcium indicators for imaging neural activity. Elife 5, e12727. 10.7554/elife.12727.

51. Chen, T.-W., Wardill, T.J., Sun, Y., Pulver, S.R., Renninger, S.L., Baohan, A., Schreiter, E.R., Kerr, R.A., Orger, M.B., Jayaraman, V., et al. (2013). Ultrasensitive fluorescent proteins for imaging neuronal activity. Nature 499, 295–300. 10.1038/nature12354.

52. Klapoetke, N.C., Murata, Y., Kim, S.S., Pulver, S.R., Birdsey-Benson, A., Cho, Y.K., Morimoto, T.K., Chuong, A.S., Carpenter, E.J., Tian, Z., et al. (2014). Independent optical excitation of distinct neural populations. Nat Methods 11, 338–346. 10.1038/nmeth.2836.

53. Sun, F., Zhou, J., Dai, B., Qian, T., Zeng, J., Li, X., Zhuo, Y., Zhang, Y., Wang, Y., Qian, C., et al. (2020). Next-generation GRAB sensors for monitoring dopaminergic activity in vivo. Nat Methods 17, 1156–1166. 10.1038/s41592-020-00981-9.

54. Gradinaru, V., Zhang, F., Ramakrishnan, C., Mattis, J., Prakash, R., Diester, I., Goshen, I., Thompson, K.R., and Deisseroth, K. (2010). Molecular and Cellular Approaches for Diversifying and Extending Optogenetics. Cell 141, 154–165. 10.1016/j.cell.2010.02.037.

55. Haber, S.N., Fudge, J.L., and McFarland, N.R. (2000). Striatonigrostriatal Pathways in Primates Form an Ascending Spiral from the Shell to the Dorsolateral Striatum. J. Neurosci. 20, 2369–2382. 10.1523/jneurosci.20-06-02369.2000.

56. Haber, S.N., and Knutson, B. (2010). The Reward Circuit: Linking Primate Anatomy and Human Imaging. Neuropsychopharmacol 35, 4–26. 10.1038/npp.2009.129.

57. Belin, D., and Everitt, B.J. (2008). Cocaine Seeking Habits Depend upon Dopamine-Dependent Serial Connectivity Linking the Ventral with the Dorsal Striatum. Neuron 57, 432–441. 10.1016/j.neuron.2007.12.019.

58. Alexander, G.E., DeLong, M.R., and Strick, P.L. (1986). Parallel Organization of Functionally Segregated Circuits Linking Basal Ganglia and Cortex. Annu. Rev. Neurosci. 9, 357–381. 10.1146/annurev.ne.09.030186.002041.

59. Mohebi, A., Wei, W., Pelattini, L., Kim, K., and Berke, J.D. (2024). Dopamine transients follow a striatal gradient of reward time horizons. Nat. Neurosci. 27, 737–746. 10.1038/s41593-023-01566-3.

60. Salinas, A.G., Lee, J.O., Augustin, S.M., Zhang, S., Patriarchi, T., Tian, L., Morales, M., Mateo, Y., and Lovinger, D.M. (2023). Distinct sub-second dopamine signaling in dorsolateral striatum measured by a genetically-encoded fluorescent sensor. Nat. Commun. 14, 5915. 10.1038/s41467-023-41581-3.

61. Calipari, E.S., Huggins, K.N., Mathews, T.A., and Jones, S.R. (2012). Conserved dorsal–ventral gradient of dopamine release and uptake rate in mice, rats and rhesus macaques. Neurochem. Int. 61, 986–991. 10.1016/j.neuint.2012.07.008.

62. Cragg, S.J., Hille, C.J., and Greenfield, S.A. (2000). Dopamine Release and Uptake Dynamics within Nonhuman Primate Striatum In Vitro. J. Neurosci. 20, 8209–8217. 10.1523/jneurosci.20-21-08209.2000.

63. Vu, M.-A.T., Brown, E.H., Wen, M.J., Noggle, C.A., Zhang, Z., Monk, K.J., Bouabid, S., Mroz, L., Graham, B.M., Zhuo, Y., et al. (2024). Targeted micro-fiber arrays for measuring and manipulating localized multi-scale neural dynamics over large, deep brain volumes during behavior. Neuron 112, 909–923.e9. 10.1016/j.neuron.2023.12.011.

64. Hamid, A.A., Frank, M.J., and Moore, C.I. (2021). Wave-like dopamine dynamics as a mechanism for spatiotemporal credit assignment. Cell 184, 2733–2749.e16. 10.1016/j.cell.2021.03.046.

65. Simpson, E.H., Akam, T., Patriarchi, T., Blanco-Pozo, M., Burgeno, L.M., Mohebi, A., Cragg, S.J., and Walton, M.E. (2024). Lights, fiber, action! A primer on in vivo fiber photometry. Neuron 112, 718–739. 10.1016/j.neuron.2023.11.016.

66. Namboodiri, V.M.K., Otis, J.M., Heeswijk, K. van, Voets, E.S., Alghorazi, R.A., Rodriguez-Romaguera, J., Mihalas, S., and Stuber, G.D. (2019). Single-cell activity tracking reveals that orbitofrontal neurons acquire and maintain a long-term memory to guide behavioral adaptation. Nat Neurosci 22, 1110–1121. 10.1038/s41593-019-0408-1.

67. Engelhard, B., Finkelstein, J., Cox, J., Fleming, W., Jang, H.J., Ornelas, S., Koay, S.A., Thiberge, S.Y., Daw, N., Tank, D.W., et al. (2019). Specialized coding of sensory, motor, and cognitive variables in VTA dopamine neurons. Nature 570, 509–513. 10.1038/s41586-019-1261-9.

68. Parker, N.F., Cameron, C.M., Taliaferro, J.P., Lee, J., Choi, J., Davidson, T.J., Daw, N.D., and Witten, I.B. (2016). Reward and choice encoding in terminals of midbrain dopamine neurons depends on striatal target. Nature Neuroscience 19. 10.1038/nn.4287.

69. Steinmetz, N.A., Zatka-Haas, P., Carandini, M., and Harris, K.D. (2019). Distributed coding of choice, action and engagement across the mouse brain. Nature 576, 266–273. 10.1038/s41586-019-1787-x.

70. Park, I.M., Meister, M.L.R., Huk, A.C., and Pillow, J.W. (2014). Encoding and decoding in parietal cortex during sensorimotor decision-making. Nat. Neurosci. 17, 1395–1403. 10.1038/nn.3800.

71. Li, J.X., Yoshida, T., Monk, K.J., and Katz, D.B. (2013). Lateral Hypothalamus Contains Two Types of Palatability-Related Taste Responses with Distinct Dynamics. J. Neurosci. 33, 9462–9473. 10.1523/jneurosci.3935-12.2013.

72. Aponte, Y., Atasoy, D., and Sternson, S.M. (2011). AGRP neurons are sufficient to orchestrate feeding behavior rapidly and without training. Nature Neuroscience 14, 351–355. 10.1038/nn.2739.

73. Krashes, M.J., Koda, S., Ye, C., Rogan, S.C., Adams, A.C., Cusher, D.S., Maratos-Flier, E., Roth, B.L., and Lowell, B.B. (2011). Rapid, reversible activation of AgRP neurons drives feeding behavior in mice. J. Clin. Investig. 121, 1424–1428. 10.1172/jci46229.

74. Betley, J.N., Cao, Z.F.H., Ritola, K.D., and Sternson, S.M. (2013). Parallel, Redundant Circuit Organization for Homeostatic Control of Feeding Behavior. Cell 155, 1337–1350. 10.1016/j.cell.2013.11.002.

75. Leib, D.E., Zimmerman, C.A., Poormoghaddam, A., Huey, E.L., Ahn, J.S., Lin, Y.-C., Tan, C.L., Chen, Y., and Knight, Z.A. (2017). The Forebrain Thirst Circuit Drives Drinking through Negative Reinforcement. Neuron 96, 1272–1281.e4. 10.1016/j.neuron.2017.11.041.

76. Matsuda, T., Hiyama, T.Y., Niimura, F., Matsusaka, T., Fukamizu, A., Kobayashi, K., Kobayashi, K., and Noda, M. (2017). Distinct neural mechanisms for the control of thirst and salt appetite in the subfornical organ. Nature Neuroscience 20. 10.1038/nn.4463.

77. Miselis, R.R. (1981). The efferent projections of the subfornical organ of the rat: A circumventricular organ within a neural network subserving water balance. Brain Res. 230, 1–23. 10.1016/0006-8993(81)90388-7.

78. Jennings, J.H., Rizzi, G., Stamatakis, A.M., Ung, R.L., and Stuber, G.D. (2013). The Inhibitory Circuit Architecture of the Lateral Hypothalamus Orchestrates Feeding. Science 341, 1517–1521. 10.1126/science.1241812.

79. O’Connor, E.C., Kremer, Y., Lefort, S., Harada, M., Pascoli, V., Rohner, C., and Lüscher, C. (2015). Accumbal D1R Neurons Projecting to Lateral Hypothalamus Authorize Feeding. Neuron 88, 553–564. 10.1016/j.neuron.2015.09.038.

80. Rinaman, L. (2010). Ascending projections from the caudal visceral nucleus of the solitary tract to brain regions involved in food intake and energy expenditure. Brain Res. 1350, 18–34. 10.1016/j.brainres.2010.03.059.

81. Pauli, J.L., Chen, J.Y., Basiri, M.L., Park, S., Carter, M.E., Sanz, E., McKnight, G.S., Stuber, G.D., and Palmiter, R.D. (2022). Molecular and anatomical characterization of parabrachial neurons and their axonal projections. eLife 11, e81868. 10.7554/elife.81868.

82. Roman, C.W., Derkach, V.A., and Palmiter, R.D. (2016). Genetically and functionally defined NTS to PBN brain circuits mediating anorexia. Nat. Commun. 7, 11905. 10.1038/ncomms11905.

83. Gutierrez, R., and Simon, S.A. (2022). Comprehensive Physiology. Compr. Physiol. 11, 2489–2523. 10.1002/cphy.c210002.

84. Gehrlach, D.A., Weiand, C., Gaitanos, T.N., Cho, E., Klein, A.S., Hennrich, A.A., Conzelmann, K.-K., and Gogolla, N. (2020). A whole-brain connectivity map of mouse insular cortex. eLife 9, e55585. 10.7554/elife.55585.

85. Burdakov, D., and Karnani, M.M. (2020). Ultra-sparse Connectivity within the Lateral Hypothalamus. Curr Biol 30, 4063–4070.e2. 10.1016/j.cub.2020.07.061.

86. Gordon-Fennell, A., and Stuber, G.D. (2021). Illuminating subcortical GABAergic and glutamatergic circuits for reward and aversion. Neuropharmacology 198, 108725. 10.1016/j.neuropharm.2021.108725.

87. Jong, J.W. de, Afjei, S.A., Dorocic, I.P., Peck, J.R., Liu, C., Kim, C.K., Tian, L., Deisseroth, K., and Lammel, S. (2018). A Neural Circuit Mechanism for Encoding Aversive Stimuli in the Mesolimbic Dopamine System. Neuron. 10.1016/j.neuron.2018.11.005.

88. Grove, J.C.R., Gray, L.A., Medina, N.L.S., Sivakumar, N., Ahn, J.S., Corpuz, T.V., Berke, J.D., Kreitzer, A.C., and Knight, Z.A. (2022). Dopamine subsystems that track internal states. Nature 608, 374–380. 10.1038/s41586-022-04954-0.

89. Augustine, V., Lee, S., and Oka, Y. (2020). Neural Control and Modulation of Thirst, Sodium Appetite, and Hunger. Cell 180, 25–32. 10.1016/j.cell.2019.11.040.

90. Robbins, T.W., and Everitt, B.J. (1992). Functions of dopamine in the dorsal and ventral striatum. Semin. Neurosci. 4, 119–127. 10.1016/1044-5765(92)90010-y.

91. Amo, R., Matias, S., Yamanaka, A., Tanaka, K.F., Uchida, N., and Watabe-Uchida, M. (2022). A gradual temporal shift of dopamine responses mirrors the progression of temporal difference error in machine learning. Nat Neurosci, 1–11. 10.1038/s41593-022-01109-2.

92. Abe, K., Kambe, Y., Majima, K., Hu, Z., Ohtake, M., Momennezhad, A., Izumi, H., Tanaka, T., Matunis, A., Stacy, E., et al. (2024). Functional diversity of dopamine axons in prefrontal cortex during classical conditioning. eLife 12, RP91136. 10.7554/elife.91136.

93. Stuber, G.D., Schwitzgebel, V.M., and Lüscher, C. (2025). The neurobiology of overeating. Neuron. 10.1016/j.neuron.2025.03.010.

94. Schultz, W. (2016). Dopamine reward prediction-error signalling: a two-component response. Nature reviews. Neuroscience 17, 183–195. 10.1038/nrn.2015.26.

95. Mohebi, A., Pettibone, J.R., Hamid, A.A., Wong, J.-M.T., Vinson, L.T., Patriarchi, T., Tian, L., Kennedy, R.T., and Berke, J.D. (2019). Dissociable dopamine dynamics for learning and motivation. Nature 570, 65–70. 10.1038/s41586-019-1235-y.

96. Threlfell, S., Lalic, T., Platt, N.J., Jennings, K.A., Deisseroth, K., and Cragg, S.J. (2012). Striatal dopamine release is triggered by synchronized activity in cholinergic interneurons. Neuron 75, 58–64. 10.1016/j.neuron.2012.04.038.

97. Sulzer, D., Cragg, S.J., and Rice, M.E. (2016). Striatal dopamine neurotransmission: Regulation of release and uptake. Basal Ganglia 6, 123–148. 10.1016/j.baga.2016.02.001.

98. Cragg, S.J. (2003). Variable Dopamine Release Probability and Short-Term Plasticity between Functional Domains of the Primate Striatum. J. Neurosci. 23, 4378–4385. 10.1523/jneurosci.23-10-04378.2003.

99. Ma, P., Chen, P., Tilden, E.I., Aggarwal, S., Oldenborg, A., and Chen, Y. (2024). Fast and slow: Recording neuromodulator dynamics across both transient and chronic time scales. Sci. Adv. 10, eadi0643. 10.1126/sciadv.adi0643.

100. Zhuang, X., Masson, J., Gingrich, J.A., Rayport, S., and Hen, R. (2005). Targeted gene expression in dopamine and serotonin neurons of the mouse brain. J. Neurosci. Methods 143, 27–32. 10.1016/j.jneumeth.2004.09.020.

101. Paxinos, G., and Franklin, K.B.J. (2001). The Mouse Brain in Stereotaxic Coordinates. 2nd Edition (Academic Press).

